# High-resolution transcriptomic profiling of the heart during chronic stress reveals cellular drivers of cardiac fibrosis and hypertrophy

**DOI:** 10.1101/854026

**Authors:** Micheal A. McLellan, Daniel A. Skelly, Malathi S.I. Dona, Galen T. Squiers, Gabriella E. Farrugia, Taylah L. Gaynor, Charles D. Cohen, Raghav Pandey, Henry Diep, Antony Vinh, Nadia A. Rosenthal, Alexander R. Pinto

**Affiliations:** The Jackson Laboratory, Bar Harbor, ME, USA.; Sackler School of Graduate Biomedical Sciences, Tufts University, Boston, MA, USA.; Baker Heart and Diabetes Research Institute, Melbourne, Victoria, Australia.; Centre for Cardiovascular Biology and Disease Research, La Trobe University, Melbourne, Victoria, Australia.

## Abstract

**Background:** Cardiac fibrosis is a key antecedent to many types of cardiac dysfunction including heart failure. Physiological factors leading to cardiac fibrosis have been recognized for decades. However, the specific cellular and molecular mediators that drive cardiac fibrosis, and the relative impact of disparate cell populations on cardiac fibrosis, remain unclear.

**Methods:** We developed a novel cardiac single-cell transcriptomics strategy to characterize the cardiac cellulome—the network of cells that forms the heart. This method was utilized to profile the cardiac cellular ecosystem in response to two weeks of continuous administration of Angiotensin II, a pro-fibrotic stimulus which drives pathological cardiac remodeling.

**Results:** This analysis uncovered multiple cell populations contributing to pathological remodeling of the extracellular matrix of the heart. Two phenotypically distinct fibroblast populations emerged after induction of tissue stress to promote fibrosis in the absence of smooth muscle actin-expressing myofibroblasts, a key pro-fibrotic cell population. Further, the cellular responses to Angiotensin II and the relative abundance of fibrogenic cells were sexually dimorphic.

**Conclusions:** These results offer a valuable resource for exploring the cardiac cellular landscape in health and after chronic cardiovascular stress. These data provide insights into the cellular and molecular mechanisms that promote pathological remodeling of the mammalian heart, highlighting early transcriptional changes which precede chronic cardiac fibrosis.

## Introduction

The mammalian heart is composed of a complex and heterogeneous network of cells. While cardiomyocytes (CMs)—the contractile cells of the heart—constitute the majority of cardiac cell mass, non-myocytes far outnumber CMs [1]. Little is known about how the individual cellular components of the heart operate together as an integrated unit, how this cellular ecosystem is altered in the context of physiological stress, and how disparate cell populations may contribute to the functional decline that accompanies cardiac injury, aging, hypertension or obesity. Moreover, while biological sex is recognized as an important variable in cardiovascular homeostasis and disease [2], little is known about how biological sex affects gene expression in disparate cardiac cell populations following chronic tissue stress.

To study the cardiac cellulome—the network of cells that forms the heart—in the context of chronic stress, we used an Angiotensin II (AngII) cardiac hypertrophy model and applied a novel method for simultaneous transcriptional profiling of single CM nuclei and non-myocyte cells. Our study revealed shared and distinct cellular pathways driving AngII-induced cardiac fibrosis. Fibrosis developed in a manner independent of myofibroblasts, a cell type conventionally recognized as a key mediator of cardiac fibrosis. This study also documents extensive sexual dimorphism in the gene expression profiles of cardiac cells, underscoring how biological sex influences responses to tissue stress. This work provides a valuable resource for cardiac cell biology research and offers important insights into the orchestrated cellular and molecular mechanisms that drive cardiac fibrosis and heart failure.

## Results

Previous large-scale single cell transcriptomic studies exploring cardiac cellular diversity have failed to profile key cardiac cell populations including CMs [3–6] due to a reliance on droplet-based sequencing approaches that are incompatible with large cell isolation. Moreover, extraction of CMs from rodent hearts typically requires retro-aortic perfusion with proteases to liberate cells from the extracellular matrix; this is a relatively time-consuming process that has limited applicability for cell preparation in transcriptomic studies [7].

To overcome these limitations, we developed a novel experimental framework to isolate and prepare all cardiac cells for single cell RNA-seq (scRNA-seq; Figure 1A). The goal of this approach was to profile all cardiac cell types with maximum possible throughput and accuracy, subject to limitations based on biological characteristics and the specifications of current scRNA-seq instrumentation. Based on recent innovations in CM isolation [8], we developed a perfusion-based tissue dissociation protocol that enables simultaneous isolation of cardiac cells from multiple hearts (see Methods). To address the challenge of CM size, CMs were denuded of their cell bodies to liberate nuclei for droplet-based transcriptional profiling. As proof of concept, both CM nuclei and non-myocyte nuclei were isolated independently and pooled for sequencing (Online Figure 1). Isolation of cells using this methodology yielded similar non-myocyte cell proportions to protocols previously employed [4] although total cell yield and the number of extracted endothelial cells was lower (Online Figure 2). Although isolation of nuclei overcame critical limitations related to CM size, for all further experimentation the non-myocyte fraction was preserved as whole cells to provide access to total cellular RNA and to allow control of input cell proportions for scRNA-seq by fluorescence-activated cell sorting using cell surface markers (Figure 1A).

**Figure 1.**
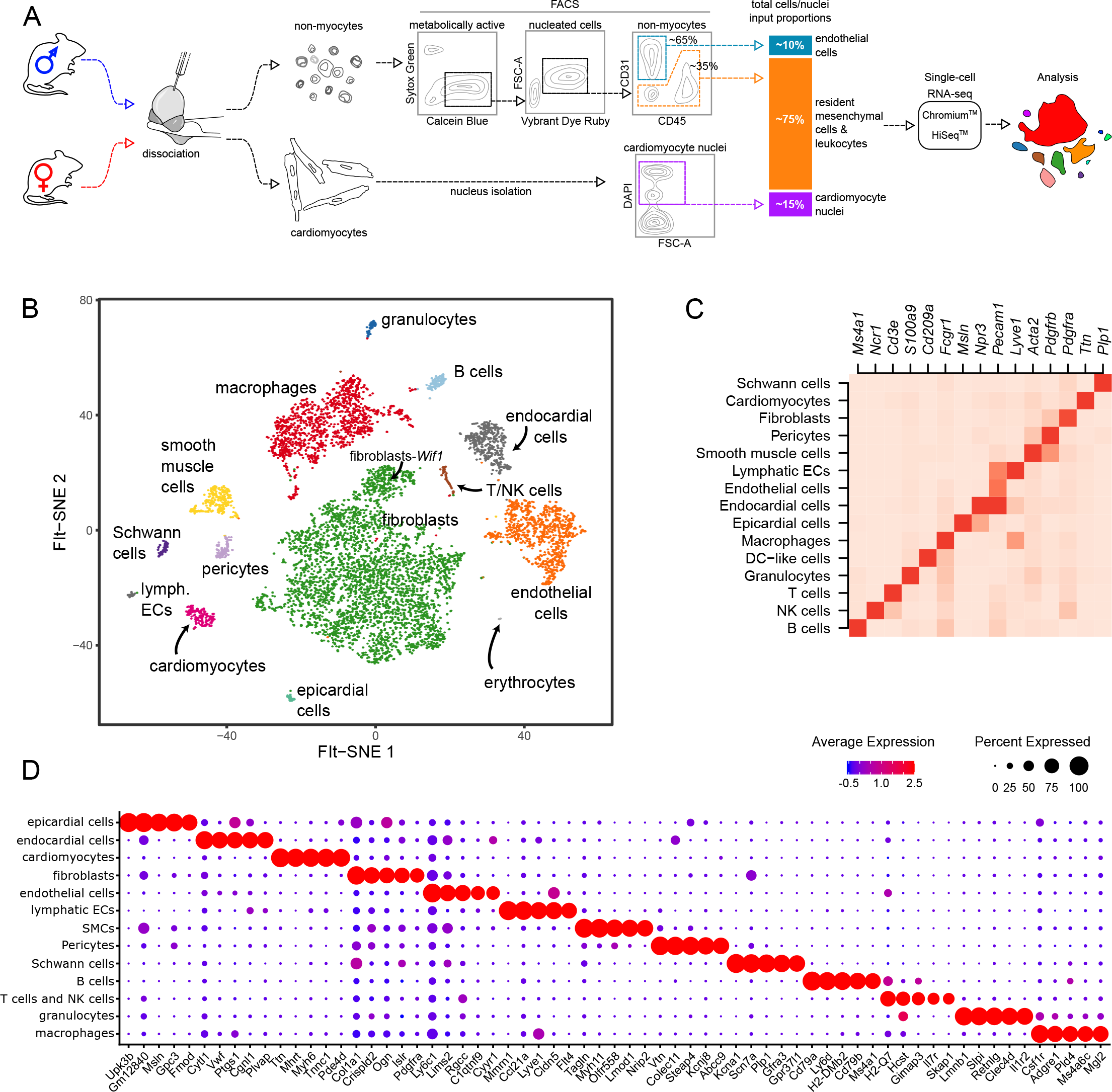
Isolation and analysis of entire cardiac cellulome by scRNA-seq. A) Schematic outline of the experimental procedure for the isolation and analysis of adult mouse cardiomyocytes and non-myocytes by scRNA-seq (see Methods). B) t-SNE projection of cardiac cells analyzed by scRNA-seq. Cells are colored by distinct cell populations as indicated. C) Heatmap of relative expression of canonical cell markers in each major cardiac cell population. D) Top five distinct genes identified for each cell population. Dot color and size indicate the relative average expression level and the proportion of cells expressing the gene, respectively, within each cell population.

**Figure 2.**
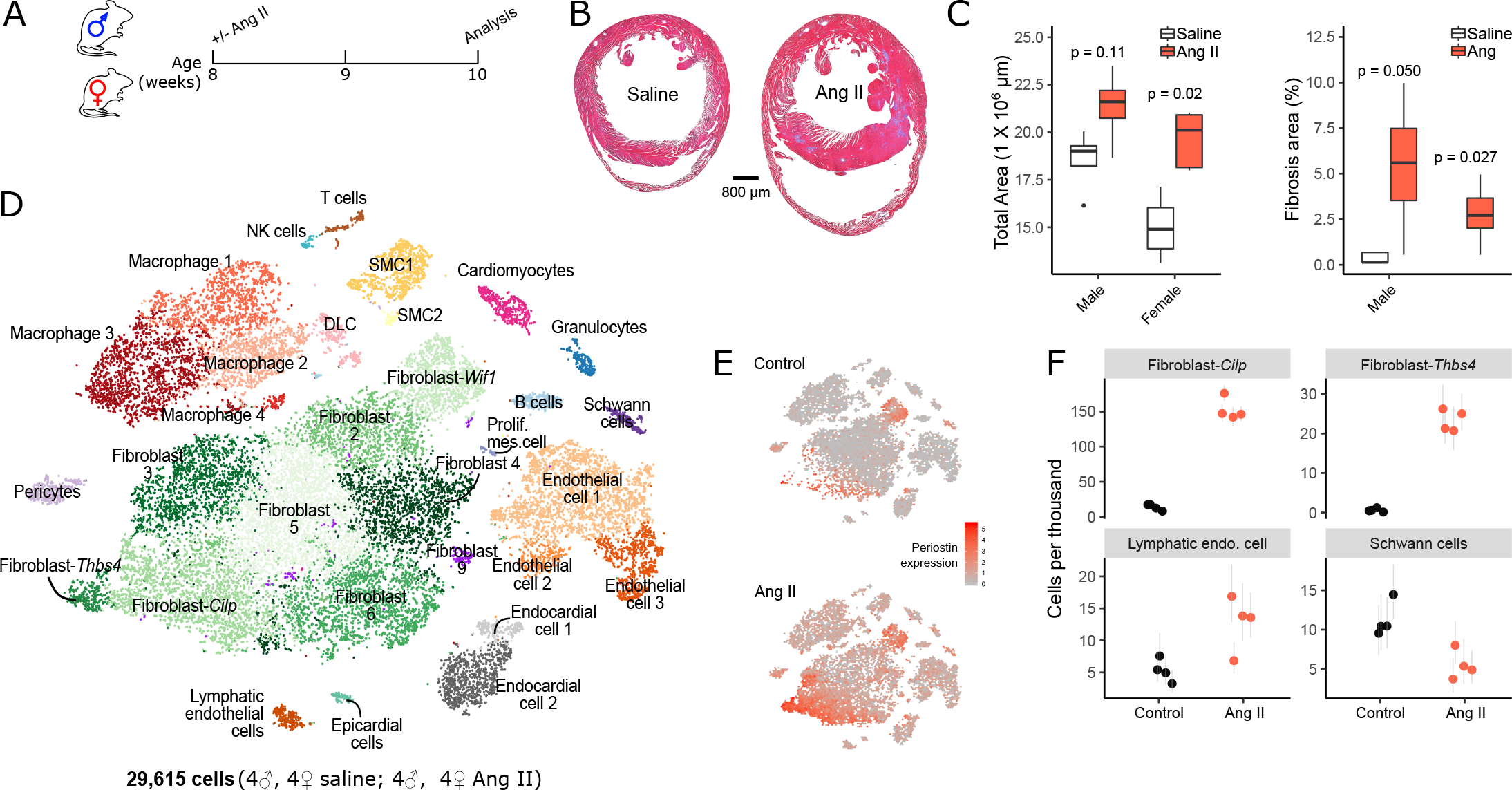
Histological and cellular changes in the heart following induction of cardiac fibrosis. A) Experimental schema for AngII mediated induction of cardiac fibrosis. Female and male mice were infused with AngII or saline (control) continuously for two weeks. B) Representative micrographs of trichrome-stained cardiac sections from mice infused with saline or AngII (left and right images, respectively). C) Change in total cardiac area and cardiac fibrosis with and without induction of fibrosis in females and males (*N*=4-5 animals per group). *P-*values shown on plot are derived from Wilcoxon test for differences in mean. D) FIt-SNE plot of cardiac cell clusters identified using single cell transcriptional profiles of cells from mouse heart with and without induction of fibrosis. E) FIt-SNE projections of cardiac cells isolated from control (top plot) and from fibrosis-induced (bottom plot) mice. Cells are colored according to Periostin (*Postn*) gene expression (red=high, gray=low). In plot, reduced dimensionality space is divided into bins and hexagons are colored according to the mean values of cells in each bin to avoid distortion of patterns through over-plotting. F) Change in proportion of selected cell populations in the scRNA-seq dataset with and without fibrosis induction. Control mice include mice implanted with saline osmotic pumps (see Methods) and those that had not undergone any surgical interventions. Online Figure 5 shows results for all cell populations.

Following isolation of both CM nuclei and non-myocyte cells, scRNA-seq was performed on a mixture of non-myocytes and myocytes from homeostatic female and male hearts. We limited the relative number of endothelial cells and CM nuclei to minimize oversampling of these abundant cell populations at the expense of resolving less frequent cell types. Following clustering of 7,474 cells (Figure 1B), examination of established marker genes and top cell population-enriched genes (Figure 1C-D) revealed a wide array of cell types that were present in all samples (Online Figure 3). These include fibroblasts (*Pdgfra, Col1a1*), pericytes (*Pdgfrb, Vtn*), smooth muscle cells (SMCs; *Acta2, Myh11*), Schwann cells (*Plp1, Kcna1*), endothelial cells (*Pecam, Ly6c1*), macrophages (*Fcgr1, Csf1r*), and other immune cell populations (granulocytes, B cells, T cells and NK cells). A subset of fibroblasts (Fibroblast-*Wif1*) showed distinct gene expression patterns including active Wnt signaling and specific expression of *Wif1*, in line with our previous report [4] and the findings of Farbehi et al [5]. Additional less well-characterized cell populations included CMs (*Ttn, Myh6*), epicardial cells (*Msln, Upk3b*), lymphatic endothelial cells (*Lyve1, Cldn5*) and endocardial cells (*Npr3, Cytl1*) (Figure 1B-D).

**Figure 3.**
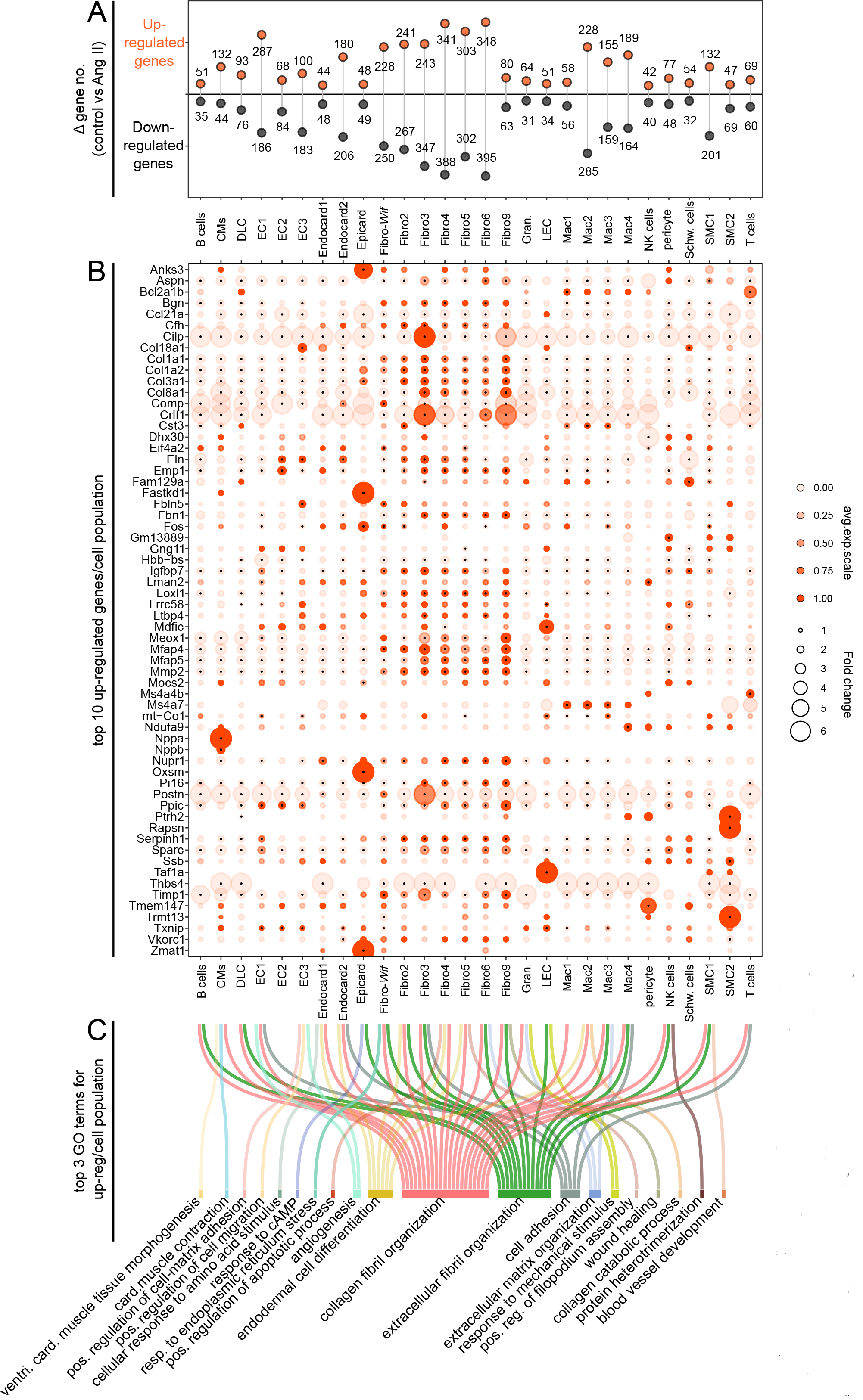
Gene expression changes in cardiac cell populations. A) Lollipop plot summarizing number of up- and downregulated genes (*p* < 0.01) in AngII-treated mouse heart cells relative to control cohorts. B) Dot plot summarizing expression of genes identified as within the top ten genes upregulated in response to AngII administration, for each cell type. Dot color and size are proportional to average expression within each cell cluster and fold increase in AngII cells relative to control cells, respectively. Black points at centers of some dots indicate a significant difference in gene expression in response to AngII (uncorrected *p* < 0.01). C) Sankey plot summarizing the top three GO terms for upregulated genes of each cell type. Connections indicate GO terms associated with each cell type. Note: (i) that many cell types upregulate genes that are associated with same GO terms, consistent with a specific and choreographed response to a cardiac stressor; (ii) not all cell types have three GO terms associates with their upregulated genes, (iii) Fibroblast-*Cilp* and Fibroblast-*Thbs4* are excluded from this differential analysis as they are not represented in controls conditions.

Cardiac tissue from mice at a relatively early stage of disease development (two weeks of chronic AngII administration) was analyzed to capture early cellular and molecular changes that lead to pathological remodeling of the heart (Figure 2A). An increase in heart mass and fibrotic area was noted in both female and male mice treated with AngII, however, hemodynamic changes and functional impairment were principally restricted to male mice that had a reduction of ejection fraction (Online Figure 4), in line with trends in human populations [9]. Histological analyses of hypertrophic mouse hearts showed a greater cross-sectional area and higher levels of fibrosis in both female and male hearts (Figure 2B-C). Together these observations describe the pathological remodeling of the heart with AngII-induced pathological hypertrophy in both female and male contexts.

**Figure 4.**
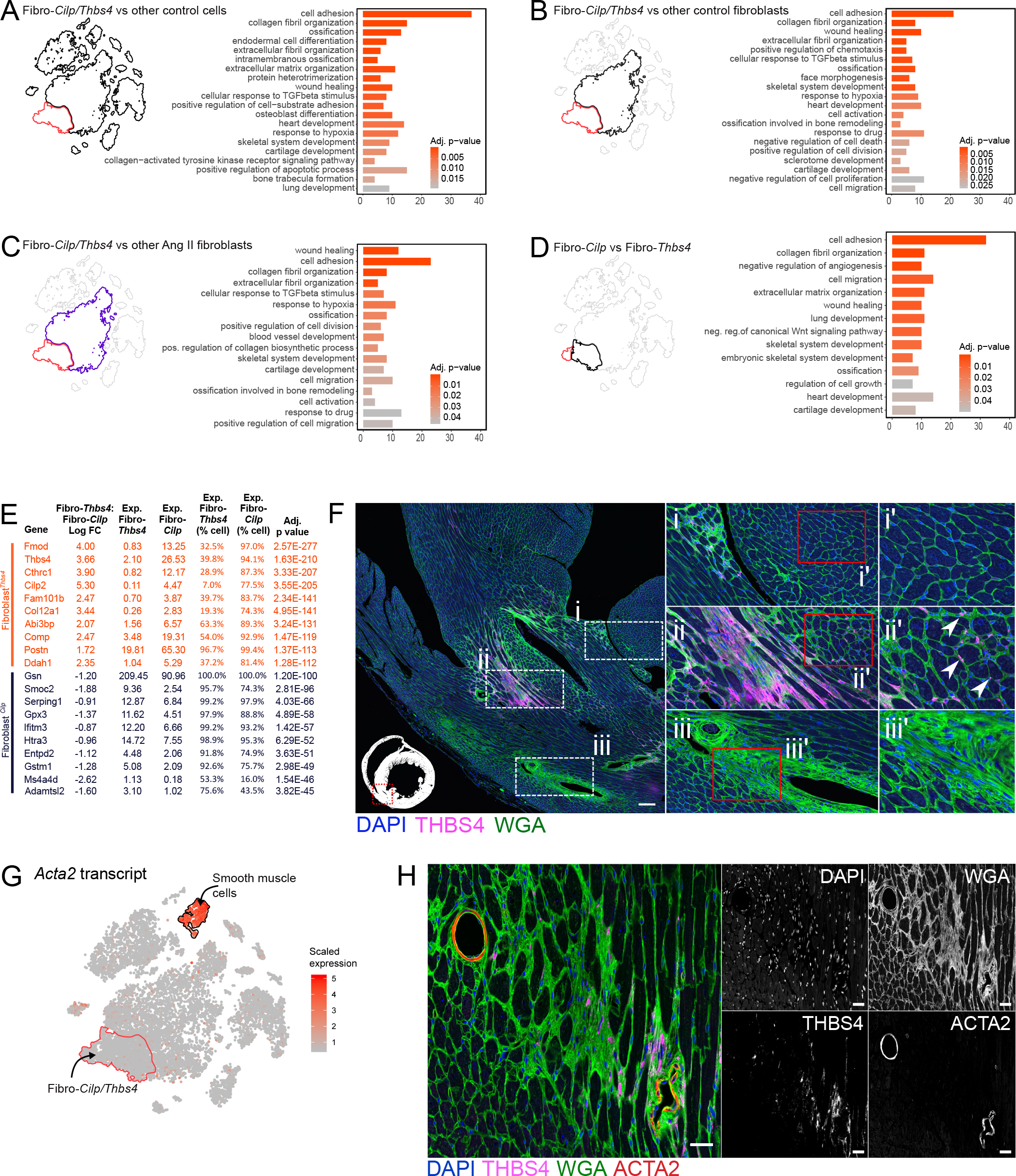
Distinguishing features of fibrosis-associated fibroblasts. A) GO terms enriched in a set of genes derived from comparing gene expression differences between Fibroblast-*Cilp* and Fibroblast-*Thbs4* vs. all other control cells. X-axis indicates the number of genes mapped to each GO term and color indicates the adjusted p-value from GO enrichment analysis. Outline of FIt-SNE projection shown on Figure 2D (left) indicates Fibroblast-*Cilp* and Fibroblast-*Thbs4* population (red) being compared to other cells (black). B) Presentation of GO terms as in (A) but associated with a comparison of Fibroblast-*Cilp* and Fibroblast-*Thbs4* vs. fibroblasts from control mice. C) Presentation of GO terms as in (A) but associated with a comparison of Fibroblast-*Cilp* and Fibroblast-*Thbs4* vs. fibroblasts from AngII-treated mice. Purple FIt-SNE outline indicates AngII-fibroblasts being compared to Fibroblast-*Cilp* and Fibroblast-*Thbs4.* D) Presentation of GO terms as in (A) but associated with a comparison of Fibroblast-*Cilp* to Fibroblast-*Thbs4.* E) Top 10 genes enriched for either Fibroblast-*Cilp* (dark blue) and Fibroblast-*Thbs4* (orange). F) Micrograph showing AngII treated mouse heart section stained with anti-THBS4 antibody, wheat-germ agglutinin (WGA; to identify cell boundaries and fibrosis) and DAPI (nuclei). Figure insets indicate regions without fibrosis (i and ii’); dense THBS4+ fibrosis (ii) and perivascular fibrosis without THBS4 staining (iii and iii’). Region with discrete THBS4+ cells is also shown (ii’). Scale bar indicates 100 µm. G) FIt-SNE plot with cells colored according to *Acta2* transcript abundance (red=high, gray=low). Red outline indicates Fibroblast-*Cilp* and Fibroblast-*Thbs4* population. H) Micrograph labeled as in F, however with additional marker for ACTA2. Monochromatic images show intensity levels of individual fluorescence channels. Scale bar indicates 25 µm.

To examine cellular and molecular changes that accompany induction of fibrosis, we performed scRNA-seq on isolated myocytes and non-myocytes in a total of 12 hearts from sham and AngII-treated mice. Combining these data with cells from the four homeostatic female and male hearts described above, we obtained a total of 29,615 cells and performed clustering to identify a diverse array of cardiac cell types and subtypes (Figure 2D). Examination of cell abundances within this scRNA-seq dataset revealed several cell populations that changed in relative prevalence in response to AngII (Figure 2E-F). Most notably, two fibroblast subpopulations which were not present at an appreciable number in unstressed hearts (Figure 2E-F; Online Figure 5) increased dramatically in hearts of mice treated with AngII and expressed high levels of periostin (Figure 2E). We refer to these cell populations as Fibroblast-*Cilp* and Fibroblast-*Thbs4*—reflecting markers that are highly expressed within these populations—with Fibroblast-*Cilp* being the largest cell population in this fibrosis dataset.

**Figure 5.**
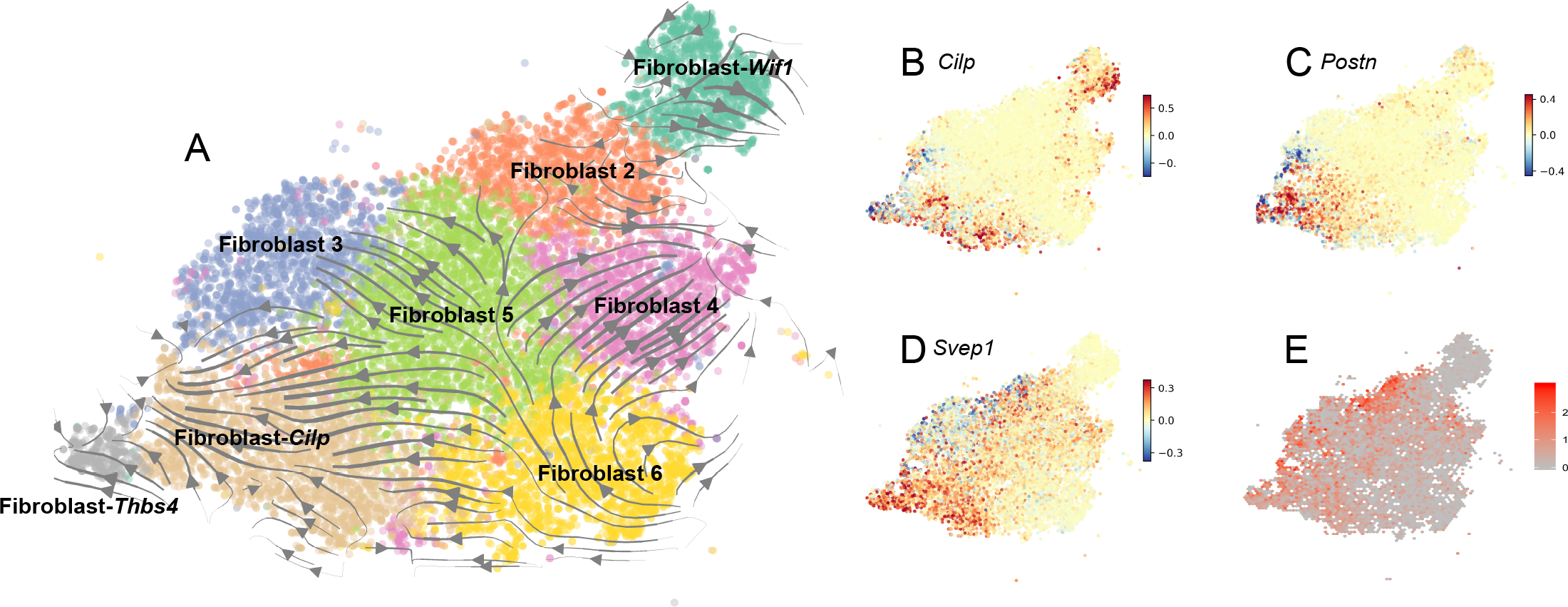
Shifts in fibroblast cell state in response to chronic stress imposed by AngII treatment. A) Stream plot depicting patterns of RNA velocity in fibroblasts from mice administered AngII. Clustering of fibroblasts and their placement in two-dimensional space are identical to Figure 2D. RNA velocities are projected into this FIt-SNE embedding for visualization purposes. Lines with arrows depict predominating changes in velocity and their associated directionality. B) Estimation of RNA velocity for *Cilp* in fibroblasts from AngII-treated mice. Relative velocities are colored according to scale on right. C) As in (B) but for *Postn.* D) As in (B) but for *Svep1.* E) Gene expression of *Svep1* in fibroblasts from AngII-treated mice.

To determine whether subpopulations of fibroblasts, endothelial/endocardial cells or macrophages/dendritic-like cells (DLCs) change in abundance following AngII treatment, we further clustered these cell populations in isolation before examining relative abundances of any new clusters (Online Figure 6). Sub-clustering fibroblasts did not significantly fragment fibroblast subpopulations further. In contrast, sub-clustering endothelial cells revealed two sub-clusters that decrease or increase in abundance following AngII treatment (Online Figure 6). Moreover, sub-clustering divided dendritic-like cells to three subpopulations, with one decreasing significantly upon hypertrophy induction. Macrophages did not fragment further compared to original clusters (Online Figure 6).

**Figure 6.**
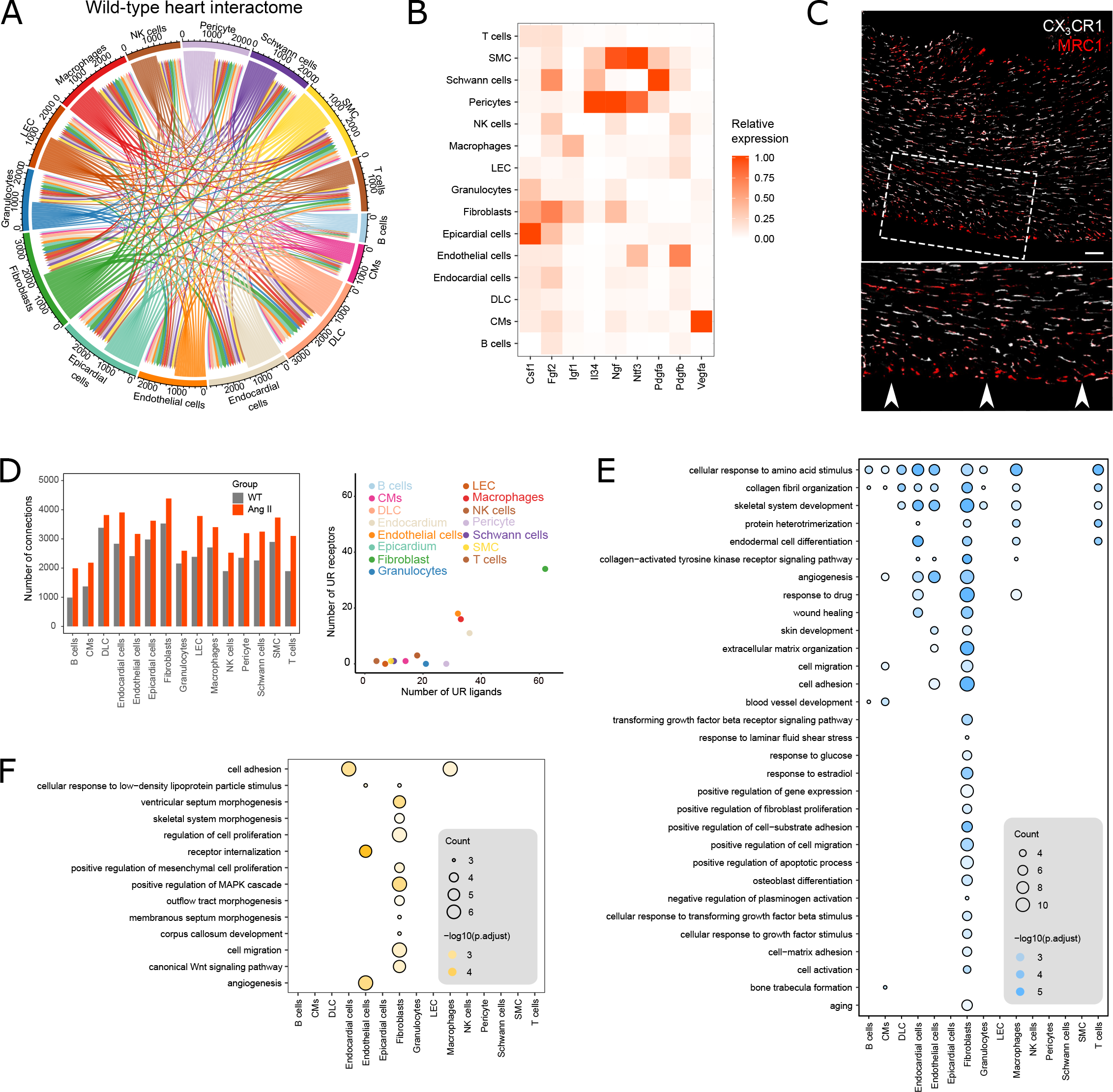
Intercellular connection network of cardiac cellulome. A) Chord plot summarizing interconnections between different cardiac cell types from hearts of control mice. B) Relative expression of a selection of essential growth factors across major cardiac cell types. Gene expression is normalized to the expression level of the cell type with the greatest mean expression. C) Spatial distribution of MRC1^+^ and MRC1^-^ macrophages relative to the epicardium in *Cx3cr1^gfp/+^* mouse hearts. GFP (white) labels both MRC1+ and MRC1-macrophages. Scale bar indicates 100 µm. Inset shows magnified view of the epicardium (indicated by white arrows). D) Number of receptors and ligands that may mediate cell-cell communication for each cell population with and without AngII treatment. Bar plot shows total number of connections (receiving and transmitted signals) made by each cell type without (grey bars) and with AngII treatment (orange bars). Scatter plot summarizes number of upregulated receptors and ligands for each cell population. Note: dots corresponding to Schwann cells, epicardial cells and B cells overlap since they have the same number of upregulated ligands and receptors. E) GO terms enriched in a set of genes that encode ligands upregulated following fibrosis induction. GO terms are ordered by their frequency of significant enrichment in different cardiac cell populations. F) GO terms enriched in a set of genes that encode receptors upregulated following fibrosis induction. GO terms are ordered by their frequency of significant enrichment in different cardiac cell populations.

The limited input of specific cell populations for scRNA-seq analysis might restrict the detection of shifts in abundances of certain cell populations. To examine cell composition using an orthogonal method, we performed flow cytometric analysis of control and AngII-treated mouse hearts (Online Figure 7). Macrophages significantly increased in abundance in males, compared to controls. However, there was no significant increase in fibroblasts detected by flow cytometry (Online Figure 7), suggesting that Fibroblast-*Cilp* and Fibroblast-*Thbs4* represent a cell state arising primarily from resident fibroblasts rather than infiltration or proliferation of cells. Indeed, single cell transcriptional profiling of cell cycle states [10] revealed no significant increase in proliferation of fibroblast populations or in other cell compartments in response to AngII (Online Figure 8).

**Figure 7.**
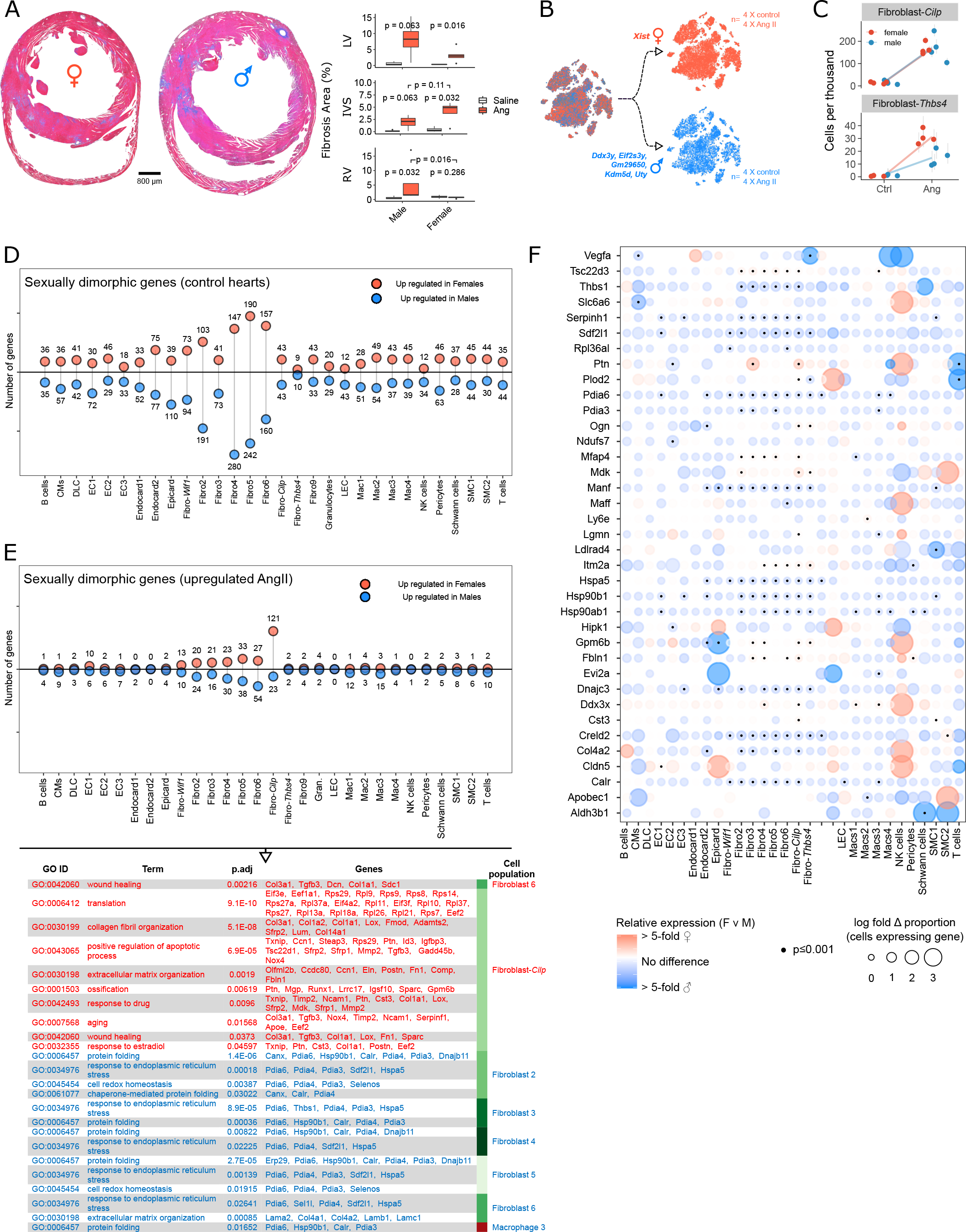
Sexual dimorphic remodeling and cellular responses. A) Trichrome stained sections from female and male hearts (left, micrographs) and quantification of fibrosis (boxplots, right). Plots summarize fibrosis in the left ventricles (LV), intraventricular septums (IVS) and right ventricles (RV) of male and female hearts. Uncorrected *p*-values determined using Wilcoxon test. B) Illustration of the desegregation of female and male cells based on sex-specific gene expression. Cells are visualized as points in FIt-SNE space, identically to Figure 2D. As expected, female and male cells are each present within each cell cluster. C) Cellular abundances of Fibroblast-*Cilp* and Fibroblast-*Thbs4* populations in females and males with and without fibrosis induction. Light grey lines indicate binomial proportion confidence intervals and are calculated using the Jeffreys interval [45]. Individual female and male mice are indicated by red and blue dots, respectively. Y-axis shows cell abundance and regression lines derived separately from female and male samples are shown in red and blue, respectively. D) Number of sexually dimorphic genes in cardiac cell types from hearts without (control) fibrosis induction. Number of genes at significantly higher levels in females and males is indicated for each cell type by red and blue circles, respectively.F) Sexually dimorphic gene expression in genes upregulated cardiac cells following AngII-treatment. See Methods and *Online* Figure 19 for details. A summary of GO terms of genes upregulated in female cells (red text) or male cells (blue text) is shown in the table below. Fibroblast-*Cilp* and Fibroblast-*Thbs4* are not shown as this analysis was restricted to cells present during both homeostasis and AngII-treatment. F) Dot plot summarizing the expression of genes changing in a sexually dimorphic manner in response to AngII. Plot shows genes that are upregulated after AngII treatment and differentially expressed between corresponding female and male cells (*p* <0.001). Dot color and size are proportional to the relative expression and relative proportion of cells expressing compared between females and males, respectively.

AngII treatment induced gene expression changes in all cardiac cell populations (Figure 3A). Most notably, fibroblast populations up- or downregulated the greatest numbers of genes, with macrophage and endothelial cell subsets also exhibiting high reactivity. Endothelial and smooth muscle cell sub-clusters (particularly endothelial1 and SMC1) and CMs exhibited highly asymmetrical patterns, with a predominance of either activating or repressive changes in gene expression (Figure 3A).

Examination of the top 10 upregulated genes for each cell type revealed several patterns relevant to hypertrophy (Figure 3B). Among the top genes upregulated upon AngII treatment were transcripts for *Cilp, Comp, Crlf1, Postn,* and *Thbs4*, some of which have been previously implicated in human heart failure [11–13]. Notably, these genes exhibit a subtle but significant increase in expression across almost all heart cell types under AngII treatment. Genes that were upregulated in clusters of cells or across a limited number of cell populations included *Ms4a7* (macrophages) and *Gm13889* (pericytes and SMCs). Upregulated genes highly restricted to one cell type included *Ccl21a* (lymphatic endothelial cells), and *Nppb* and *Nppa* (in CMs), which are well-established biomarkers of heart failure [14]. Gene ontology (GO) analysis of AngII-upregulated genes revealed that the top three GO terms in most cardiac cell types contribute to remodeling of the cardiac ECM (Figure 3C).

Of the top 10 downregulated genes, epicardial cells exhibited the greatest number of cell-specific downregulated genes (*B4galt5*, *Cst9*, and *Has1*; Online Figure 9A) with others such as *Spic1* (Mac1), *Cd207* (DLCs), *Ptpdc1* (Fibroblast 9), *Thap8* (Mac1), *Arhgap17* (NK cells), *Letmd1* (NK cells), and *Atp1a2* (T cells) restricted to single cell types (Online Figure 9A). Multiple genes related to ATP synthesis and mitochondrial activity (e.g. *mt-Co2*, *mt-Nd4l*) were also downregulated, particularly in macrophages and endothelial cell populations (Online Figure 9A). GO analysis of downregulated genes revealed multiple coherent gene programs induced by AngII (Online Figure 9B), the most frequent of which were those associated with protein translation, particularly amongst fibroblasts, as well as pathways related to inflammation and infiltration of monocytes and other leukocytes.

Abundances of Fibroblast-*Cilp* and Fibroblast-*Thbs4* changed most dramatically upon treatment with AngII (Figure 2F). To determine the impact of these cell populations on cardiac remodeling, we first compared differentially expressed genes between Fibroblast-*Cilp* and Fibroblast-*Thbs4* compared to all other control cells within this dataset, cardiac fibroblasts from control animals, and fibroblasts from AngII-treated animals (Figures 4A-C, respectively). Features common to basic fibroblast biology defined Fibroblast-*Cilp* and Fibroblast-*Thbs4* when compared to all cardiac cells (e.g. collagen and ECM organization). When compared to all fibroblasts from control animals or AngII-treated cohorts (Figure 4B and C, respectively), collagen remodeling genes were the primary distinguishing features of Fibroblast-*Cilp* and Fibroblast-*Thbs4.* Comparison of Fibroblast-*Cilp* and Fibroblast-*Thbs4* also showed higher levels of ECM remodeling genes in Fibroblast-*Cilp* compared to Fibroblast-*Thbs4* (Figure 4D). Indeed, examination of total transcripts corresponding to ECM remodeling genes suggests that Fibroblast-*Cilp* and Fibroblast-*Thbs4* are key contributors to ECM remodeling, with the Fibroblast-*Cilp* population contributing greater summed transcript abundance than all other cell populations in AngII-treated mice (Online Figure 10).

Fibroblast-*Cilp* and Fibroblast-*Thbs4* were also distinguished by the upregulation of genes such as *Thbs4*, *Fmod, Cthrc1* and *Cilp*2 (Figure 4E). No genes were uniquely and highly expressed in Fibroblast-*Cilp* that were not also expressed in Fibroblast-*Thbs4*. THBS4^+^ cells in AngII-treated hearts (Figure 4F) were aggregated within foci and could also be identified as discrete cells (Figure 4F, *i* and *ii*, respectively; Online video 1). An absence of THBS4^+^ cells surrounding perivascular fibrotic lesions (Figure 4F, *iii*), suggested Fibroblast-*Thbs4* are principally involved in interstitial fibrosis. Finally, in accordance with transcriptomic analyses, we confirmed that neither Fibroblast-*Cilp* nor Fibroblast-*Thbs4* correspond to ACTA2^+^ myofibroblasts (Figure 4G and H) using mice from two independent mouse colonies. These findings suggest that Fibroblast-*Cilp* and Fibroblast-*Thbs4* may be transcriptionally similar to the matrifibrocyte, a specialized fibroblast state found in the mature scar after MI [15]. While Fibroblast-*Cilp/Thbs4* had some expression of the matrifibrocyte markers *Comp*, *Sfrp2,* and *Wisp2* [6, 15], these genes are less specific for Fibroblast-*Cilp* and Fibroblast-*Thbs4* than *Cilp*, *Ddah1*, or *Thbs4* (Online Figure 11).

To explore the origins of Fibroblast-*Cilp* and Fibroblast-*Thbs4*, we examined dynamic patterns of transcriptional activation and repression in detail in all fibroblasts using RNA velocity [16], as implemented in scVelo [17]. This analysis suggested that the Fibroblast-*Thbs4* population predominantly arises from Fibroblast-*Cilp*, while Fibroblast-*Cilp* has broader origins from Fibro5 and Fibro6 (Figure 5A). Use of two independent pseudotime-based methods yielded results that were also concordant with these conclusions (Online Figure 12). Some transcripts expressed in both Fibroblast-*Cilp* and Fibroblast-*Thbs4*, such as *Cilp* and *Postn*, showed gradients in velocity across Fibroblast-*Cilp* into Fibroblast-*Thbs4* (Figure 5B-C), suggesting that Fibroblast-*Thbs4* cells represent a cell state characterized by transcriptional activation of a unique set of genes. Certain genes that were not clearly fibroblast subpopulation-specific yet appeared to play a role in AngII-induced remodeling included *Svep1,* the second-ranked gene differing in velocity between Fibroblast-*Cilp*/Fibroblast-*Thbs4* and the remaining fibroblast subpopulations (the top-ranked gene, *Adcy7*, is likely involved in cross-talk with AngII signaling pathways in cardiac fibroblasts [18]). Figure 5D demonstrates that *Svep1* is strongly activated in Fibroblast-*Cilp*/Fibroblast-*Thbs4*, consistent with the recognized role of *Svep1* as a GWAS-implicated hypertension risk gene [19]. Notably, steady-state gene expression levels alone did not clearly link *Svep1* expression to cell populations specific to AngII-treatment (Figure 5E).

To determine the impact of AngII treatment on cardiac intercellular signaling, we mapped ligands and cognate receptors expressed by various cell populations (Figure 6). As previously reported [4], a similar analysis revealed extensive intercellular communication in homeostatic hearts. Signaling by fibroblasts, along with macrophages, represented key features of the interstitial cardiac niche at homeostasis (Figure 6A; Online Figure 13). DLCs, epicardial cells and endocardial cells were also significant populations forming extensive potential signaling networks within a diverse array of cell types (Figure 6A, Online Figure 13). In contrast, consistent with previous analyzes [4] patrolling cell populations—granulocytes, NK cells and lymphocytes—were amongst the least interactive cell populations. Few interactions observed for CMs may be attributable to a lower number of genes and reads in this cell population due to transcriptional profiling of nuclei rather than whole cells.

Examination of factors essential for supporting specific cell populations points to important trophic intercellular interactions (Figure 6B). In addition to previously reported expression patterns for *Igf1*, *Il34*, *Ngf* and *Ntf3* (Figure 6B; [4]), epicardial cells were identified as the most highly enriched for *Csf1* transcripts, required for macrophage vitality, while CMs were the primary producers of *Vegfa* that supports endothelial cell growth (Figure 6B). Although the role of epicardial cells in supporting macrophage trafficking in fetal development has been previously established [20], this result suggests that epicardial cells are the most macrophage-trophic cell population by *Csf1/Csf1r* signaling (Figure 6B). Indeed, the spatial distribution of macrophage subsets relative to the epicardium is distinct (Figure 6C), and further accentuated in older animals [21], underscoring the epicardium and sub-epicardial space as a macrophage-trophic environment in the adult heart.

Cell-cell signaling in response to AngII-induced hypertrophy was increased in all cell types studied, with fibroblasts increasing the greatest number of connections (Figure 6D). GO enrichment analysis of upregulated connections between cell types revealed that the most frequently increased ligands involved *cellular response to amino acid stimulus* and *collagen fibril organization* (Figure 6E); upregulated receptors included those interacting with ECM (Figure 6F). Notably, ligands involved in angiogenesis were upregulated in multiple cell types, while endothelial cells upregulated cognate receptors (Figure 6F). Together these analyzes indicate a concerted effort by multiple cell types in the cardiac cellulome to alter the cardiac niche and drive ECM remodeling.

Conversely, GO terms related to downregulated ligands showed a clear dampening of inflammation in the cardiac microenvironment (Online Figure 14), with almost all enriched GO terms related to inflammation or leukocytes/monocyte trafficking. Intriguingly, other GO terms enriched within downregulated ligands and receptors were associated with angiogenesis, suggesting the up- and downregulation of receptors within endothelial cells may reflect changes in cellular phenotype and shifts in sensitivity to angiotrophic cues.

Analysis of hemodynamic and cardiac functional parameters, as well as quantification of tissue fibrosis, suggested extensive sexual dimorphism in the development of pathological remodeling in response to AngII. Many upregulated genes were modulated by estradiol (Online Figure 15). Sexually dimorphic differences in cardiac cellularity after AngII treatment were detected (Online Figure 7) including development of fibrosis in specific anatomical loci (Figure 7A).

To explore sex-specific gene expression patterns within specific populations of the cardiac cellulome, we isolated female and male cells from control and AngII-treated groups based on the expression of genes expected to differ between sexes (chromosome X and Y genes outside of the pseudoautosomal region; Figure 7B; Online Figure 16). Comparison of proportions of female and male cell populations in control and AngII-treated hearts revealed no differences in cell abundance except within the Fibroblast-*Thbs4* population, which was approximately two-fold higher abundance of these cells in females compared to males (Figure 7C).

Comparison of sex-specific gene expression differences within cell populations from both control and AngII-treated hearts revealed highly sexually dimorphic gene expression in almost all cell populations, with the greatest level of sexually dimorphic gene expression occurring within fibroblast subpopulations 2 and 4-6 in control hearts and fibroblast subpopulations 2-6 and Fibroblast-*Cilp* of AngII-treated mice (Figure 7D-E). This is in line with the high enrichment of sex hormone receptors expressed by these cells (Online Figure 17). Consistent with a positive relationship between cell number and power to detect sexually dimorphic gene expression, we observed a slight yet statistically significant correlation between cell population size and the number of sexually dimorphic genes identified (Online Figure 18).

Sexual dimorphism in expression of genes upregulated following AngII treatment (Figure 7E-F, Online Figure 19) were detected in fibroblast populations (with the exception of Fibroblast-*Wif1* and Fibroblast-*Thbs4*), with very few genes differentially regulated in other cell types. (Figure 7E). Very few dimorphisms in expression were identified amongst genes downregulated following cardiac fibrosis (Online Figure 20). Examination of genes with 2-fold or greater difference between sexes in cells of control hearts revealed a heterogeneous collection of genes (Online Figure 21A). These included genes encoding proteins such as GNGT2, CAMLG, TDP1, and NEK7 that have been previously identified as sexually dimorphic in human heart failure [22]. GO enrichment analysis of sexually dimorphic genes in control hearts showed an enrichment of genes corresponding to antigen processing in multiple female cell populations and protein folding, stress and angiogenesis in multiple male cell types (Online Figure 21B). Consistent with sexually dimorphic regulation of the endothelium, flow cytometric analysis of endothelial cells showed greater numbers of endothelial cells in male mouse hearts (Online Figure 22). Genes such as *Vegfa* were upregulated in male (but not female) CMs and Fibroblast-*Thbs4* populations from AngII-treated mice, further suggesting dimorphic angiogenic stimulation in females and males (Figure 7E). Amongst genes upregulated or downregulated in either sex (Online Figure 19), female hearts had an enrichment of ECM remodeling genes amongst the upregulated genes in Fibro6 and Fibroblast-*Cilp*, whereas males showed upregulation of genes involved in protein folding or responses to endoplasmic reticulum stress (Figure 7E). We also observed upregulation of ECM-organizing genes in male Fibro6 cells that were distinct from those upregulated in female Fibro6 or Fibroblast-*Cilp* populations, underscoring a potential difference in the quality of ECM generated in the two sexes upon tissue stress. GO enrichment analysis revealed no coherent biological themes shared among downregulated genes, principally in fibroblasts, whose expression was sexually dimorphic.

To determine whether Fibroblast-*Cilp* and Fibroblast-*Thbs4* populations—those that emerge following induction of cardiac stress—are also present in the context of human pathological remodeling, we analyzed bulk cardiac RNA-seq data from 85 female and 172 male humans between the ages of 40 and 69 profiled by the Genotype-Tissue Expression (GTEx) Project. To segregate humans without and with putative hypertrophy *NPPB* expression was examined; this gene encodes the preprohormone preproBNP from which the cardiac hypertrophy biomarker B-type natriuretic peptide is derived [23]. Samples with the highest (top 20%) *NPPB* expression (without regard to age or sex) were classified as putatively hypertrophic (Online Figure 23A). Expression of *POSTN*, *TGFβ1* and collagens was significantly elevated in all hypertrophy groups compared to corresponding ‘no-hypertrophy’ controls (uncorrected *p* < 1e-07 for all genes; Online Figure 23B) confirming that human ‘hypertrophy’ cohorts also exhibit other markers of cardiac stress. Expression of *THBS4*, *DDAH1, FMOD, CTHRC1*, and *CILP,* five genes that are enriched in Fibroblast-*Cilp* and Fibroblast-*Thbs4,* was significantly increased in hypertrophic human hearts (1.4e-09 < *p* < 2e-03 for all genes, uncorrected; Online Figure 23). The overall concordance of these observations is striking due to the limitations of this underpowered statistical analysis, including imperfect status of *NPPB* transcript as a biomarker for heart failure, the inconsistency of human heart sample isolation, and age-dependent changes in the systemic milieu such as the onset of menopause. Nevertheless, our observations suggest that cells corresponding to Fibroblast-*Cilp* and Fibroblast-*Thbs4* may be present in the context of human cardiac stress and motivate future studies of the detailed molecular mechanisms initiating the emergence of these cells as well as the physiological impacts of modifying their function.

## Discussion

While there exists a great deal of data concerning individual cellular components which contribute to cardiac dysfunction, less is known about how these components interact during concerted pathological responses to tissue stress. To clarify the roles of distinct heart cell populations in the context of chronic maladaptive cardiac stress responses leading to fibrosis, we developed a strategy enabling unbiased analysis of myocyte and non-myocyte transcriptomes in parallel using scRNA-seq, producing a high-resolution atlas of the cardiac cellulome in the context of homeostasis and fibrosis. Virtually all cell types in the heart have detectable transcriptional responses to tissue stress, with most cells participating in remodeling of the ECM. Specifically, two activated fibroblast populations emerge in response to stress, presumably to remodel the extracellular environment by direct ECM deposition and paracrine regulation of the heart cell niche. The extensive sexual dimorphism of cell-specific gene expression in resting and maladaptive hearts implicates sex-specific mechanisms driving cardiac physiology and stress responses. Overall, this study provides insight into the cellular and molecular drivers of cardiac pathological remodeling and fibrosis.

As a resource for researchers, the extensive dataset presented here comprises a deep and comprehensive map of the cardiac cellulome. These data were derived from eight individual scRNA-seq libraries, with each library comprised of both female and male cells. Cells from the 16 individual biological specimens can be isolated *in silico* to moderate effects of outliers and to examine the effect of biological sex on cardiac homeostasis and disease, an under-examined area in both cardiac cell biology and physiology [2, 24].

Our analysis attained a detailed portrait of the intercellular circuitry supporting the heart cell ecosystem at homeostasis and in response to the induction of pathological hypertrophy. For example, this dataset implicates epicardial cells as critically important for supporting macrophage vitality and trafficking in the adult heart. Fibroblast-derived *Csf1* and mural-cell-derived *Il34* are both growth factors that support macrophages which we previously detected in the heart [4], however, the current analysis found that epicardial cells are the most macrophage-trophic cardiac cell population as a result of their high production of *Csf1*. Moreover, we identified distinct spatial distributions of macrophage subsets at the epicardium, suggesting additional unidentified regulation of macrophage positioning and/or trafficking. These results further support the important role of epicardial cells in macrophage homeostasis, in accordance with previous studies demonstrating the role of the epicardium in macrophage colonization of the heart [20]. Also notable was the identification of CMs as the principal producers of *Vegfa,* which is a dominant endothelial and angiogenesis growth factor. While we have not examined the precise isoforms of *Vegfa* expressed by cardiac cells [25], our analysis indicates that the cell type perhaps most dependent on tissue perfusion is the key driver of vascular innervation of the heart.

Examination of ligand genes upregulated following induction of hypertrophy revealed the entire cardiac cellular niche is altered to support ECM remodeling and angiogenesis. This corresponded with upregulation of receptor genes mediating cell adhesion in a wide variety of cells, whereas angiogenesis-related receptors were upregulated in endothelial cells. GO enrichment analysis of all genes upregulated following induction of fibrosis indicated that a primary response of most cell types is the activation of genes involved in ECM remodeling. However, Fibroblast-*Cilp* and Fibroblast-*Thbs4*, which principally emerge following cardiac stress, appear to be the most fibrogenic cell populations based on expression of transcripts related to ECM remodeling GO terms. Their fibrogenic phenotype is further underscored when compared to other fibroblast populations from AngII-treated mice that extensively activate ECM remodeling and collagen genes.

Fibroblast-*Cilp* and Fibroblast-*Thbs4* do not constitute classically defined myofibroblast populations—widely accepted as the main protagonists driving fibrosis [26]. Recent scRNA-seq studies examining MI have readily detected *Acta2+* myofibroblasts [5,6,27,28]. The experimental conditions of these reports represent examples of tissue trauma where myofibroblast activation may mediate wound healing. In contrast, *Acta2* transcript was not detected in Fibroblast-*Cilp* nor in Fibroblast-*Thbs4* above background transcriptomic levels, or at the protein level by histology. A lack of smooth muscle α-actin would be consistent with the absence of a need for structural support of a necrotic area. Indeed, fibrosis without myofibroblast conversion has been reported in a model of diabetic cardiomyopathy [29] and this fibrosis dataset underscores that accumulation of extensive fibrosis in the heart is not dependent on ACTA2^+^ myofibroblasts.

Indeed, Fibroblast-*Cilp* and Fibroblast-*Thbs4* more closely resemble matrifibrocytes that reside in a mature scar after MI [15]. Although expression of key matrifibrocyte marker genes— *Comp*, *Sfrp2* or *Wisp2*—were not highly enriched in cells defined as Fibroblast-*Cilp* and Fibroblast-*Thbs4* in this study, the functional inferences made of these cell populations are comparable. Therefore, cells analogous to Fibroblast-*Cilp* and Fibroblast-*Thbs4* may promote fibrosis in multiple cardiac stress contexts.

Extending on these findings, we examined expression of transcriptional signatures resembling Fibroblast-*Cilp* and Fibroblast-*Thbs4* in the human context. Specifically, using human heart bulk RNA-seq data from normal and hypertrophic cardiac samples, we found that genes with restricted expression to Fibroblast-*Cilp* and Fibroblast-*Thbs4* are upregulated in hypertrophy and correlate with the cardiac stress marker *Nppb*. This finding suggests that future studies targeting Fibroblast-*Cilp* and Fibroblast-*Thbs4*, or genes they specifically activate, may yield therapeutically relevant outcomes for addressing human disease.

Analysis of downregulated genes indicates that a key feature of fibrosis induction is dampening of inflammation and leukocyte infiltration. Downregulated genes included Ccl ligands, which are key monocyte chemoattractants. Although AngII infusion caused an increase in macrophage abundance using flow cytometry, this was not accompanied by significant increases in pro-inflammatory mediators. This suggests that the cardiac cellulome functions to dampen inflammation in order to regulate inflammatory responses. Further, we found mitochondrial bioenergetics are subdued in many cell types, particularly in endothelial cells and macrophages, in accordance with the association between mitochondrial activity and heart failure.

The current study also identified numerous cell type-specific genes that were robustly upregulated in hypertrophy (such as *Rapsn* in SMCs or *Fastkd1* in epicardial cells) with unknown implications for cardiac pathology. Similar relationships were observed amongst downregulated genes (for example *Thap8* in macrophages or *Has1* in the epicardium). These genes constitute appealing subjects for further research to explore their expression in further detail, and to their validity as potential therapeutic targets.

Analysis of the abundance of Fibroblast-*Cilp* and Fibroblast-*Thbs4* cell populations suggest sex differences in cellular abundance (Figure 7C), which is of particular interest in light of known sexual dimorphisms in cardiac stiffness [30]. Indeed, analysis of cardiac fibrosis in mice treated with AngII revealed differences in fibrotic area in addition to the distribution of fibrotic loci. Females were characterized by less fibrosis in the left ventricle wall and the absence of fibrosis in the right ventricle walls, as well as less perivascular fibrosis. In contrast, greater fibrosis accumulated in the female interventricular septum compared to males, although a functional decline was only observed in males.

In both the non-stressed and stressed contexts, sexual dimorphisms were observed in cardiac gene expression across almost all cell types, principally present in fibroblasts that are the major sex hormone-receptor expressing cells in the heart. In the context of cardiac fibrosis, female and male cardiac fibroblasts regulated distinct sets of genes that are associated with ECM remodeling, suggesting cardiac ECM quality is biologically distinct in the two sexes.

A key sexually dimorphic characteristic is the upregulation of genes related to unfolded protein responses (UPR) and endoplasmic reticulum (ER) stress in multiple male cell populations. Indeed, UPR may at least partly explain two sexually dimorphic features in this fibrosis dataset. First, an increase in UPR may contribute to the increased fibrosis observed in male hearts (Figure 7A). ER stress is observed in multiple models of tissue fibrosis including multiple models of maladaptive cardiac remodeling [31] and attenuation of UPR inhibits fibrosis [32–34]. Second, UPR may contribute to downregulation of translation-related genes in males. UPR may elicit this by multiple means including IRE1-mediated translational control of the transcription factor *Xbp1*, and regulation of multiple genes including *eukaryotic initiation factor* (*Eif*) genes and others involved in protein translation [31]. Incidentally, Xbp1 is upregulated in multiple male cardiac cell types, and is associated with tissue fibrosis in the heart and elsewhere [35, 36].

In addition to the link between protein translation and UPR, translation is also supported by estrogens in several species and in both neoplastic and non-neoplastic tissues [37–39]. In tumor cells, inhibition of *estrogen receptor α* induces UPR [37]. Moreover, estrogen has been recently shown to regulate *Eif* genes to promote protein synthesis [40]. Many *Eif* genes upregulated after AngII infusion across multiple female cell populations in addition to many ribosomal protein genes. Together, these observations suggest female cells are better primed for protein synthesis as well as avoidance of tissue fibrosis and maladaptive remodeling.

Conversely, the enrichment of genes implicated in angiogenesis was observed in multiple male cardiac cell populations including Fibro 2, Fibro 6, Fibroblast-*Cilp* and Fibroblast-*Thbs4* from non-stressed mice. Accordingly, increased numbers of endothelial cells were found in male hearts compared to females. In addition, significantly higher levels of *Vegfa* expression was detected in male CMs and Fibroblast-*Thbs4* cells after fibrosis induction. While estrogen has been reported to enhance angiogenesis in a range of contexts [41], more recent research suggests that loss of estrogen receptor promotes angiogenesis [42]. Additionally, testosterone induces angiogenesis in a sex-dependent manner, further underscoring the importance of the hormonal milieu [43]. Finally, UPR is also strongly linked to upregulation of pro-angiogenic genes [44], resulting in a more pro-angiogenic environment in male hearts.

The findings of this study offer unique insights into the multifaceted mechanisms driving maladaptive remodeling of the heart, including important components of this process that are sexually dimorphic. They also highlight the utility of the single cell transcriptomics data and the overall approach presented in this study for exploring the cardiac cellulome and its changes in the context of chronic tissue stress. Further analysis of cardiac cell networks, with integration of data from different developmental and disease states, time points, and genetic backgrounds, will further advance our understanding of cardiac cellularity in health and disease.

## METHODS

### Animals

All experimental procedures performed on C57BL/6J mouse strain following protocols approved by Jackson Laboratory Animal Research ethics committee. Angiotensin II (1.5 or 1.44 mg/kg/day) or saline infusion was performed on 8-week-old female and male mice over a period of two weeks before subsequent analyses.

### Cardiac single-cell preparation, sequencing and analysis

Cardiac single-cell and single-nucleus suspensions of non-myocytes and cardiomyocyte nuclei, respectively, were prepared using a modified from Ackers-Johnson et al., 2016 [8] before subsequent processing as outlined in Figure 1A. Single-cell RNA-seq was performed using the 10X Genomics Chromium Single Cell 3’ Library and Gel bead Kit v2.

For additional information see Online Materials and Methods.

## Non-conventional acronyms and abstractions

scRNA-seq: single-cell RNA sequencing
Fibroblast-*Cilp*: fibroblasts with high expression of *Cilp*
Fibroblast-*Thbs4*: fibroblasts with high expression of *Thbs4*
Fibroblast-*Wif1*: fibroblasts with high expression of *Wif1*
SMC: smooth muscle cell
LEC: lymphatic endothelial cells
CM: cardiomyocyte
ECM: extracellular matrix
ER: endoplasmic reticulum
UPR: unfolded protein response

## ACKNOWLEDGMENTS

We acknowledge the use of JAX Flow Cytometry Core, Microscopy Core, Surgical Services, Histology Service, and Single-Cell Biology Laboratory. JAX Cores are supported by the Jackson Laboratory Cancer Center Core grant and the Leducq Foundation Transatlantic Network of Excellence in Cardiac Research to NR. The Genotype-Tissue Expression (GTEx) Project was supported by the Common Fund of the Office of the Director of the National Institutes of Health, and by NCI, NHGRI, NHLBI, NIDA, NIMH, and NINDS. The data used for the analyses described in this manuscript were obtained from the GTEx Portal on 02/21/2019. We acknowledge the use Monash Micro Imaging facility at the Alfred Research Alliance for provision of microscopy instrumentation and training.

**Online Figure 1. Isolation and analysis of cardiac cell nuclei for scRNA-seq. A)** Example images of isolated cardiomyocytes using the perfusion based cardiac single cell preparation method (see Online Materials and Methods for more detail) (top, phase contrast micrograph scale bar indicates 100 µm) and isolated cardiomyocyte nuclei imaged using the Amnis ImageStream system (bottom) showing DAPI and phase images of single nuclei. **B)** tSNE projection displaying cardiomyocyte nuclei (indicated) and non-myocyte nuclei analyzed by single-nucleus RNA-seq, as proof of concept for single-nucleus sequencing from live cardiac tissue (see Online Materials and Methods).

**Online Figure 2. Comparison of two cardiac cell isolation techniques on cellular composition and yield. A)** Flow cytometry gating strategy to identify relative proportions of endothelial cells (ECs), resident mesenchymal cells (RMCs) and leukocytes. Note: only non-myocytes are included in this analysis. **B)** Quantified proportions of three major cardiac cell types (% Calcein^+^ cells). Endothelial cell (EC) yield is significantly lower when performing the perfusion protocol, however significantly greater proportions of leukocytes are liberated by perfusion. Resident mesenchymal cell proportions are not altered by either protocol. Statistical analysis was conducted using an unpaired t-test. FSC-H = forward scatter height, FSC-W = forward scatter width, FSC-A = forward scatter area.

**Online Figure 3. Cellular composition of mouse hearts at homeostasis.** Cellular composition of the sampling of sequenced cells derived from mouse hearts at homeostasis. Cells from two samples of wild type mice are shown in columns (batch). Each sample consisted of cells from female and male mice (Materials and Methods). Colors indicate cell populations and are identical to Figure 1B. Heights of individual boxes comprising each bar represent the proportion of cells classified as each cell population. Cellular composition shown here is not directly comparable to a homeostatic mouse heart due to the sorting strategy employed (Figure 1A).

**Online Figure 4. Physiological assessment of cardiac function after AngII reveals sexual dimorphism in functional cardiac decline. A-D)** Echocardiographic assessment of experimental mice prior to AngII stimulus (Male/Female-pre) and after two weeks AngII treatment, or saline control. Echocardiographs were used to calculate (A) ejection fraction percentage, (B) fractional shortening percentage, (C) diastolic volume, and (D) systolic volume. Statistical analysis was conducted using an unpaired t-test, *P< 0.05, **P< 0.005, ***P< 0.0005. **E)** Conscious blood pressure assessment after AngII, or saline control, in males and females. *P< 0.05, **P< 0.005, ***P< 0.0005. **F)** Heart-weight to body-weight ratio after AngII, or saline control, in males and females. ***P< 0.0005

**Online Figure 5. Cellular composition of mouse hearts at homeostasis and after AngII treatment. A)** Cellular composition of the sampling of sequenced cells derived from mouse hearts at homeostasis, after administration of saline (two weeks), and after administration of AngII (two weeks). Cells from individual samples are shown in columns (batch). Each sample consisted of cells from female and male mice (Materials and Methods). Colors indicate cell populations and sub-populations and are used throughout this study. Heights of individual boxes comprising each bar represent the proportion of cells classified as each cell population. Cellular composition shown here is not directly comparative to hearts isolated from treated and untreated mice due to the sorting strategy employed (Online Figure 1A). **B)** As in (A) but showing individual points representing observed cell abundances (in units of cells per thousand sequenced) of each population. Individual AngII-treated or control (wild type or saline administration) samples mice are indicated by black and red dots, respectively. Light grey lines indicate binomial proportion confidence intervals and are calculated using Jeffreys interval.

**Online Figure 6. Sub-clustering of endothelial cells and macrophages.** Top two tSNE projections display endothelial cell and macrophage/DC-like cell populations (left and right, respectively) with cells colored based on clustering shown on Figure 2D. Bottom two tSNE projections display endothelial cells and macrophage populations (left and right, respectively) with cells colored based on new clusters identified following re-clustering. Dot-and-whisker plots summarize relative abundances (in percentage) of new clusters identified following re-clustering of cell populations. The cluster each plot corresponds to is indicated by the number above each plot. Statistical analysis was performed using Wilcoxon-test.

**Online Figure 7. Sexual dimorphism in cardiac cellular composition changes in response to AngII detected by flow cytometry A)** Flow cytometry gating strategies to identify relative proportions of live, metabolically active (C+V+) cardiac cell types. **B)** Quantified proportions of each cell type normalized to the mean of each WT control (WT = 1). ECs were significantly elevated in male mice given AngII compared to females given AngII. Leukocytes were also significantly increased in male AngII mice compared to male WT controls within the major cardiac cell compartment. Both vascular and lymphatic ECs are elevated in male AngII mice compared to male WT controls and female AngII counterparts. No changes were observed within the resident mesenchymal cell compartment. However, in the leukocyte compartment male AngII mice exhibited significantly elevated myeloid cells compared to both male WT mice as well as female AngII mice. Consequently, these aforementioned increases in leukocyte proportions were observed in macrophages including their polarization sub-types MHCII^hi^ and MHCII^lo^. Statistical analysis was conducted using an unpaired t-test, where significance was determined as P< 0.05. FSC-H = forward scatter height, FSC-W = forward scatter width, FSC-A = forward scatter area, SSC-H = side scatter height.

**Online Figure 8. Cell cycle scoring to identify dividing cardiac cells from mice with and without AngII treatment.** Cell cycle scores are determined by considering expression levels G2/M and S phase marker genes and is based on the strategy by Tirosh et al., 2016 [10].

**Online Figure 9. Downregulated genes after induction of fibrosis. A)** Dot plot summarizing the top 10 downregulated genes for each cell population. Duplicate entries of genes have been removed. Fold change (FC) in gene expression is indicated by circle size. Circle color indicates relative expression level in cells from control samples (expression level before reduction). Red dots indicate reduction in gene expression where p≤0.01 (uncorrected). **B)** Enrichment of GO terms in downregulated genes. Number of genes downregulated (circle size) and adjusted p-value (circle color) are indicated.

**Online Figure 10. Total transcript corresponding to ECM-related GO terms.** Dot plot showing expression of genes classified within the GO categories shown on y-axis. Circle size represents the sum of transcripts for genes corresponding to each GO term within each cell population.

**Online Figure 11. Markers of matrifibrocytes and Fibroblast-*Cilp* and Fibroblast-*Thbs4*.** Figures show FItSNE projections for cardiac cell populations (as shown in Figure 2D) with relative gene expression indicated by color (red=high, gray=low).

**Online Figure 12. Shifts in fibroblast cell state in response to chronic stress imposed by AngII treatment.** Left-Pseudotime trajectories of fibroblasts isolated from AngII-treated mice inferred using monocle3. Cells are visualized using UMAP dimensionality reduction and colored according to fibroblast cluster. Gray line indicates inferred pseudotime trajectory. Right-Pseudotime trajectories of fibroblasts isolated from AngII-treated mice inferred using slingshot. Cells are visualized using UMAP dimensionality reduction and colored according to fibroblast cluster. Black line indicates inferred pseudotime trajectory.

**Online Figure 13. Transmitted and received signals by cardiac cells with and without AngII treatment. A)** Number of signals transmitted as determined by ligand encoding genes which are expressed by cells which have cognate receptors expressed by other cardiac cell populations. **B)** Number of signals received as determined by receptor encoding genes expressed by cells which have cognate ligands expressed by other cardiac cell populations.

**Online Figure 14. GO terms corresponding to downregulated ligand and receptor genes. A)** GO terms enriched in downregulated ligand genes. **B)** GO terms enriched in downregulated receptor genes.

**Online Figure 15. Summed transcript levels of genes annotated as responsive to estradiol that are upregulated in cardiac cells following AngII treatment** A bar plot which summarizes the total transcript abundance for genes that are upregulated in cardiac cells following AngII treatment and included in the ‘response to estradiol’ GO term.

**Online Figure 16. Distribution of males and female cells based on expression of Y chromosome-related genes and *Xist*.** Blue and red dots indicate male and female cells, respectively. Grey dots represent cells which could not be discriminated. Genes considered are outside of the pseudoautosomal regions of the X and Y chromosomes (see Online Materials and Methods).

**Online Figure 17. Sex hormone receptors in cardiac cells from mice with and without AngII treatment.** Violin plots generated with Seurat showing the expression of various hormone receptors across all cardiac cell types. Androgen receptor expression (*Ar*) is greatest in cardiac fibroblasts. Estrogen receptor α and estrogen receptor β (*Esr1*, *Esr2*, respectively) are both lowly expressed, but most highly detected in cardiac fibroblasts. The membrane bound G protein-coupled estrogen receptor 1 (*Gper1*) is also lowly expressed, but greatest in endothelial cells and pericytes.

**Online Figure 18. Relationship between number of sexually dimorphic genes discovered and cell population size.** Top panel summarizes the relationship in cardiac cell populations from control animals. Bottom panel summarizes the relationship in cardiac cell populations from AngII-treated animals.

**Online Figure 19. Strategy for determining sexually dimorphic genes regulation following AngII treatment (related to figure 7D-F).** For each cell population, female and male cells were considered in isolation and genes upregulated or downregulated following AngII treatment determined. For dimorphisms in upregulated genes (results shown Figure 7D-F), a list of genes upregulated in either females or males was generated and sexual dimorphisms in gene expression was examined. For dimorphisms in downregulated genes (results shown Online Figure 20), a list of genes downregulated in either sex was generated and dimorphism in expression was again examined.

**Online Figure 20. Sexual dimorphism in gene expression amongst genes that are downregulated in cardiac cells following AngII treatment.** The number of genes which are downregulated in response to AngII as well as exhibiting sexually dimorphic gene expression, presented for each cardiac cell type. In this context upregulated refers to a gene that is greater in either sex, but overall remains lower after AngII. Strategy to determine sexually dimorphisms is outlined in Online Figure 19.

**Online Figure 21. Sexually dimorphic genes in cell populations from control mouse hearts. A)** A dot plot showing genes differentially expressed at 2-fold or greater between males and females. Red and blue circles represent genes which are more highly expressed in females or males respectively. Circle size represents the difference in the percentage of cells expressing the gene between sexes. Solid black dots indicate a statistically significant difference in expression with p≤0.01 (unadjusted). **B)** GO terms enriched in sexually dimorphic gene sets in female and male cardiac cell populations. Red and blue text indicate GO terms enriched in sets of genes that are upregulated in females and males, respectively.

**Online Figure 22. Abundances of endothelial cells in female and male hearts determined by flow cytometry A)** Flow cytometry contour plots outlining the gating strategy used to determine frequency of endothelial cells. **B)** Absolute number of endothelial cells normalized to heart tissue mass. **C)** Proportion of endothelial cells relative to total non-myocytes.

**Online Figure 23. GTEx bulk RNA-seq analysis from human hearts. A)** Expression of *NPPB* (in units of transcripts per million; tpm) in 257 female and male human hearts as a function of age and sex. Dotted red horizontal line indicates threshold used for classifying samples as putatively hypertrophic (highest 20% *NPPB* expression without regard to age or sex). **B)** Expression of markers of pathological cardiac remodeling in human hearts. Boxplots show identical human samples to (A) classified as positive or negative for putative hypertrophy. To facilitate comparison of patterns for genes that have different absolute expression levels, gene expression levels (in units of tpm) were rescaled to set a maximum of one and minimum of zero. *P-*values provided above each gene are derived from linear models for the effect of hypertrophy classification while controlling for age and sex. **C)** Summed expression of transcripts of genes encoded by mitochondrial DNA for identical human samples to (A) classified as positive or negative for putative hypertrophy. *P-*values provide below the boxplots is derived from a linear model for the effect of hypertrophy classification while controlling for age and sex. **D)** Expression of human orthologs to markers of Fibroblast-*Thbs4* in human hearts. Boxplots show identical human samples to (A) classified as positive or negative for putative hypertrophy. To facilitate comparison of patterns for genes that have different absolute expression levels, gene expression levels (in units of tpm) were rescaled to set a maximum of one and minimum of zero. *P-*values provided above each gene are derived from linear models for the effect of hypertrophy classification while controlling for age and sex.

**Online Figure 24. Cardiac tissue perfusion assembly for parallel perfusion-based dissociation of cardiac cells from multiple hearts. A)** A close-up view of a heart being perfused through the apex by digestion enzymes. **B)** Equipment layout for simultaneous perfusion of multiple hearts.

**Online Video 1. Spatial distribution of Fibroblast-*Thbs4* cells in AngII-treated mouse hearts.** 3D projection video of a fluorescence micrograph from an AngII-treated mouse heart. The tissue is stained with anti-THBS4 antibody (red), wheat-germ-agglutinin (blue), anti-ACTA2 antibody (green) and DAPI (white). Red, blue, green and white represent Fibroblast-*Thbs4*, cell boundaries, ACTA2+ cells (SMCs and myofibroblasts) and cell nuclei, respectively.

## References

1. Pinto AR, Ilinykh A, Ivey MJ, Kuwabara JT, D’antoni ML, Debuque RJ, et al. Revisiting cardiac cellular composition. Circ Res 2016;118:400–409. doi:10.1161/CIRCRESAHA.115.307778.

2. Blenck CL, Harvey PA, Reckelhoff JF, Leinwand LA. The importance of biological sex and estrogen in rodent models of cardiovascular health and disease. Circ Res 2016;118:1294–1312. doi:10.1161/CIRCRESAHA.116.307509.

3. DeLaughter DM, Bick AG, Wakimoto H, McKean D, Gorham JM, Kathiriya IS, et al. Single-Cell Resolution of Temporal Gene Expression during Heart Development. Dev Cell 2016;39:480–490. doi:10.1016/j.devcel.2016.10.001.

4. Skelly DA, Squiers GT, McLellan MA, Bolisetty MT, Robson P, Rosenthal NA, et al. Single-Cell Transcriptional Profiling Reveals Cellular Diversity and Intercommunication in the Mouse Heart. Cell Rep 2018;22. doi:10.1016/j.celrep.2017.12.072.

5. Farbehi N, Patrick R, Dorison A, Xaymardan M, Janbandhu V, Wystub-Lis K, et al. Single-cell expression profiling reveals dynamic flux of cardiac stromal, vascular and immune cells in health and injury. Elife 2019;8. doi:10.7554/eLife.43882.

6. Forte E, Skelly DA, Chen M, Daigle S, Morelli KA, Hon O, et al. Dynamic Interstitial Cell Response During Myocardial Infarction Predicts Resilience to Rupture in Genetically Diverse Mice. SSRN Electron J 2019. doi:10.2139/ssrn.3436216.

7. van den Brink SC, Sage F, Vértesy Á, Spanjaard B, Peterson-Maduro J, Baron CS, et al. Single-cell sequencing reveals dissociation-induced gene expression in tissue subpopulations. Nat Methods 2017;14:935–936. doi:10.1038/nmeth.4437.

8. Ackers-Johnson M, Li PY, Holmes AP, O’Brien SM, Pavlovic D, Foo RS. A Simplified, Langendorff-Free Method for Concomitant Isolation of Viable Cardiac Myocytes and Nonmyocytes from the Adult Mouse Heart. Circ Res 2016;119:909– 920. doi:10.1161/CIRCRESAHA.116.309202.

9. Lee DS, Gona P, Vasan RS, Larson MG, Benjamin EJ, Wang TJ, et al. Relation of disease pathogenesis and risk factors to heart failure with preserved or reduced ejection fraction: Insights from the framingham heart study of the national heart, lung, and blood institute. Circulation 2009;119:3070–3077. doi:10.1161/CIRCULATIONAHA.108.815944.

10. Tirosh I, Izar B, Prakadan SM, Wadsworth MH, Treacy D, Trombetta JJ, et al. Dissecting the multicellular ecosystem of metastatic melanoma by single-cell RNA-seq. Science (80-) 2016;352:189–196. doi:10.1126/science.aad0501.

11. Zhao S, Wu H, Xia W, Chen X, Zhu S, Zhang S, et al. Periostin expression is upregulated and associated with myocardial fibrosis in human failing hearts. J Cardiol 2014;63:373–378. doi:10.1016/j.jjcc.2013.09.013.

12. Van Nieuwenhoven FA, Munts C, Op’T Veld RC, González A, Diéz J, Heymans S, et al. Cartilage intermediate layer protein 1 (CILP1): A novel mediator of cardiac extracellular matrix remodelling. Sci Rep 2017;7. doi:10.1038/s41598-017-16201-y.

13. Tan FL, Moravec CS, Li J, Apperson-Hansen C, McCarthy PM, Young JB, et al. The gene expression fingerprint of human heart failure. Proc Natl Acad Sci U S A 2002;99:11387–11392. doi:10.1073/pnas.162370099.

14. Sergeeva IA, Christoffels VM. Regulation of expression of atrial and brain natriuretic peptide, biomarkers for heart development and disease. Biochim Biophys Acta - Mol Basis Dis 2013;1832. doi:10.1016/j.bbadis.2013.07.003.

15. Fu X, Khalil H, Kanisicak O, Boyer JG, Vagnozzi RJ, Maliken BD, et al. Specialized fibroblast differentiated states underlie scar formation in the infarcted mouse heart. J Clin Invest 2018;128:2127–2143. doi:10.1172/JCI98215.

16. La Manno G, Soldatov R, Zeisel A, Braun E, Hochgerner H, Petukhov V, et al. RNA velocity of single cells. Nature 2018;560:494–498. doi:10.1038/s41586-018-0414-6.

17. Bergen V, Lange M, Peidli S, Wolf FA, Theis FJ. Generalizing RNA velocity to transient cell states through dynamical modeling. BioRxiv 2019:820936. doi:10.1101/820936.

18. Ostrom RS, Naugle JE, Hase M, Gregorian C, Swaney JS, Insel PA, et al. Angiotensin II enhances adenylyl cyclase signaling via Ca 2+/calmodulin: Gq-Ga cross-talk regulates collagen production in cardiac fibroblasts. J Biol Chem 2003;278:24461– 24468. doi:10.1074/jbc.M212659200.

19. Patel RS, Masi S, Taddei S. Understanding the role of genetics in hypertension. Eur Heart J 2017;38:2309–2312. doi:10.1093/eurheartj/ehx273.

20. Stevens SM, von Gise A, VanDusen N, Zhou B, Pu WT. Epicardium is required for cardiac seeding by yolk sac macrophages, precursors of resident macrophages of the adult heart. Dev Biol 2016;413:153–159. doi:10.1016/j.ydbio.2016.03.014.

21. Pinto AR, Godwin JW, Chandran A, Hersey L, Ilinykh A, Debuque R, et al. Age-related changes in tissue macrophages precede cardiac functional impairment. Aging (Albany NY) 2014;6:399–413. doi:10.18632/aging.100669.

22. Fermin DR, Barac A, Lee S, Polster SP, Hannenhalli S, Bergemann TL, et al. Sex and Age Dimorphism of Myocardial Gene Expression in Nonischemic Human Heart Failure. Circ Cardiovasc Genet 2008;1:117–125. doi:10.1161/CIRCGENETICS.108.802652.

23. Moe GW. B-type natriuretic peptide in heart failure. Curr Opin Cardiol 2006;21:208– 214. doi:10.1097/01.hco.0000221582.71619.84.

24. Maric-Bilkan C, Arnold AP, Taylor DA, Dwinell M, Howlett SE, Wenger N, et al. Maric-Bilkan et al NHLBI WG Report on Sex Differences in Cardiovascular Research 803 Choosing Appropriate Experimental Models to Study Sex Differences. Hypertension 2016;67.

25. Kikuchi R, Nakamura K, MacLauchlan S, Ngo DTM, Shimizu I, Fuster JJ, et al. An antiangiogenic isoform of VEGF-A contributes to impaired vascularization in peripheral artery disease. Nat Med 2014;20:1464–1471. doi:10.1038/nm.3703.

26. Gourdie RG, Dimmeler S, Kohl P. Novel therapeutic strategies targeting fibroblasts and fibrosis in heart disease. Nat Publ Gr 2016;15. doi:10.1038/nrd.2016.89.

27. Gladka MM, Molenaar B, de Ruiter H, van der Elst S, Tsui H, Versteeg D, et al. Single-Cell Sequencing of the Healthy and Diseased Heart Reveals Ckap4 as a New Modulator of Fibroblasts Activation. Circulation 2018:CIRCULATIONAHA.117.030742. doi:10.1161/CIRCULATIONAHA.117.030742.

28. Kretzschmar K, Post Y, Bannier-Hélaouët M, Mattiotti A, Drost J, Basak O, et al. Profiling proliferative cells and their progeny in damaged murine hearts. Proc Natl Acad Sci U S A 2018:201805829. doi:10.1073/pnas.1805829115.

29. Alex L, Russo I, Holoborodko V, Frangogiannis NG. Characterization of a mouse model of obesity-related fibrotic cardiomyopathy that recapitulates features of human heart failure with preserved ejection fraction. Am J Physiol Heart Circ Physiol 2018;315:H934–H949. doi:10.1152/ajpheart.00238.2018.

30. Redfield MM, Jacobsen SJ, Borlaug BA, Rodeheffer RJ, Kass DA. Age- and Gender-Related Ventricular-Vascular Stiffening. Circulation 2005;112:2254–2262. doi:10.1161/CIRCULATIONAHA.105.541078.

31. Kropski JA, Blackwell TS. Endoplasmic reticulum stress in the pathogenesis of fibrotic disease. J Clin Invest 2018;128:64–73. doi:10.1172/JCI93560.

32. Ayala P, Montenegro J, Vivar R, Letelier A, Urroz PA, Copaja M, et al. Attenuation of endoplasmic reticulum stress using the chemical chaperone 4-phenylbutyric acid prevents cardiac fibrosis induced by isoproterenol. Exp Mol Pathol 2012;92:97–104. doi:10.1016/j.yexmp.2011.10.012.

33. Groenendyk J, Lee D, Jung J, Dyck JRB, Lopaschuk GD, Agellon LB, et al. Inhibition of the unfolded protein response mechanism prevents cardiac fibrosis. PLoS One 2016;11. doi:10.1371/journal.pone.0159682.

34. Kassan M, Galán M, Partyka M, Saifudeen Z, Henrion D, Trebak M, et al. Endoplasmic reticulum stress is involved in cardiac damage and vascular endothelial dysfunction in hypertensive mice. Arterioscler Thromb Vasc Biol 2012;32:1652–1661. doi:10.1161/ATVBAHA.112.249318.

35. Duan Q, Ni L, Wang P, Chen C, Yang L, Ma B, et al. Deregulation of XBP1 expression contributes to myocardial vascular endothelial growth factor-A expression and angiogenesis during cardiac hypertrophy in vivo. Aging Cell 2016;15:625–633. doi:10.1111/acel.12460.

36. Kim RS, Hasegawa D, Goossens N, Tsuchida T, Athwal V, Sun X, et al. The XBP1 Arm of the Unfolded Protein Response Induces Fibrogenic Activity in Hepatic Stellate Cells Through Autophagy. Sci Rep 2016;6:39342. doi:10.1038/srep39342.

37. Andruska ND, Zheng X, Yang X, Mao C, Cherian MM, Mahapatra L, et al. Estrogen receptor α inhibitor activates the unfolded protein response, blocks protein synthesis, and induces tumor regression. Proc Natl Acad Sci U S A 2015;112:4737–4742. doi:10.1073/pnas.1403685112.

38. Briz V, Baudry M. Estrogen regulates protein synthesis and actin polymerization in hippocampal neurons through different molecular mechanisms. Front Endocrinol (Lausanne) 2014;5. doi:10.3389/fendo.2014.00022.

39. Noteboom WD, Gorski J. An early effect of estrogen on protein synthesis. Proc Natl Acad Sci United States 1963;50:250–255. doi:10.1073/pnas.50.2.250.

40. Cuesta R, Berman AY, Alayev A, Holz MK. Estrogen receptor promotes protein synthesis by fine-tuning the expression of the eukaryotic translation initiation factor 3 subunit f (eIF3f). J Biol Chem 2019;294:2267–2278. doi:10.1074/jbc.RA118.004383.

41. Losordo DW, Isner JM. Estrogen and angiogenesis: A review. Arterioscler Thromb Vasc Biol 2001;21:6–12. doi:10.1161/01.ATV.21.1.6.

42. Likhite N, Yadav V, Milliman EJ, Sopariwala DH, Lorca S, Narayana NP, et al. Loss of Estrogen-Related Receptor Alpha Facilitates Angiogenesis in Endothelial Cells. Mol Cell Biol 2019;39. doi:10.1128/mcb.00411-18.

43. Sieveking DP, Lim P, Chow RWY, Dunn LL, Bao S, McGrath KCY, et al. A sex-specific role for androgens in angiogenesis. J Exp Med 2010;207:345–352. doi:10.1084/jem.20091924.

44. Binet F, Sapieha P. ER Stress and Angiogenesis. Cell Metab 2015; 22:560–575. doi:10.1016/j.cmet.2015.07.010.

45. Brown LD, Cai TT, Das Gupta A. Interval estimation for a binomial proportion. Stat Sci 2001;16:101–117. doi:10.1214/ss/1009213286.

