## Supplementary material for "High-resolution transcriptomic profiling of the heart during chronic stress reveals cellular drivers of cardiac fibrosis and hypertrophy": Online Methods and Figures

### Online Materials and Methods

**Animals and physiological assessment**—The experiments described in this study were conducted using 8-14 week old female and male wild-type or *Cx3cr1*$^{gfp/+}$ mice in the C57BL/6J background. Mice were purchased from The Jackson Laboratory (Bar Harbor ME, USA) or the Walter and Eliza Hall Institute (Melbourne, Australia). All experiments performed were approved by The Jackson Laboratory Institutional Animal Care and Use Committee (IACUC) or La Trobe University Animal Ethics Committee (AEC16-93).

Hypertension-induced cardiac hypertrophy and fibrosis experiments were achieved by subcutaneous implantation of osmotic pumps (Alzet, model 2002) loaded with angiotensin II or vehicle. Similar protocols were performed at the two sites, except Ang II was administered at either 1.5 or 1.44 mg/kg/day at The Jackson Laboratory or La Trobe University, respectively. Surgical procedures conducted at the Jackson Laboratory were undertaken by The Jackson Laboratory Surgical Services platform. Echocardiography was performed on the Vevo3100 system at The Center for Biometric Analysis at The Jackson Laboratory. For analysis, B-mode and M-mode measurements were utilized in Vevo LAB software (VisualSonics). Conscious blood pressure detection was performed using a MC4000 tail cuff apparatus (Hatteras Instruments, NC, USA) at the La Trobe Animal Research and Training Facility (LARTF).

**Histology and immunostaining**—Hearts were fixed in formalin supplemented with 30mM KCl (SigmaAldrich) and subsequently were paraffin-embedded (FFPE) then sectioned at 10 µm thickness for histological and immunohistochemical analyses. Masson's trichrome staining was performed to visualize fibrosis following standard protocols. For fluorescence imaging, sections were stained with antibodies targeting thrombospondin IV (THBS4, 1:200, MAB7860SP, R&D Systems) and smooth muscle actin (ACTA2, 1:500, 55135-1-AP, Proteintech Group). Sections were additionally stained with Wheat Germ Agglutinin (WGA, 1:100, 29022-1, Biotium) and DAPI (1 µg/mL, D9542, Sigma Aldrich) before acquiring confocal micrographs. Brightfield or fluorescence micrographs were subsequently analyzed using Fiji software for quantification of tissue area and fibrosis (Masson trichrome images) and generation of composite fluorescence images.

**Cardiac tissue thick section immunostaining—**Cardiac thick sections were prepared and stained based on previously described protocols [1]. Briefly, hearts were fixed in 4% (w/v) formaldehyde and 150 µm sections prepared. After permeabilization (0.5% Triton X-100/PBS) and blocking, sections were stained with antibodies targeting *mannose receptor C-type 1* (Mrc1;1:200, CO68C2, BioLegend) and GFP (1:200, ab13970, Abcam). Sections were additionally stained with Wheat Germ Agglutinin (WGA, 1:100, 29022-1, Biotium) and DAPI (1 µg/mL, D9542, Sigma Aldrich) before acquiring confocal micrographs. Micrographs were subsequently processed using Imaris software (Bitplane). Online video 1 was prepared using Imaris and annotated using Inkscape (https://inkscape.org) and Blender (http://www.blender.org) software packages.

**Simultaneous single cell and single nucleus isolation and sequencing**—For simultaneous isolations of cardiomyocytes and non-myocytes, we adapted a protocol previously described for perfusion-based dissociation of the heart [2]. Recipes for all buffers used can be found in the aforementioned protocol with the following modifications, 2-3, butanedione monoxime (BDM) was replaced with 5mM blebbistatin (SigmaAldrich) and [0.1U/µL] murine RNase inhibitor (NEB) was also supplemented. One-hour prior to tissue harvest, mice were administered an intraperitoneal heparin injection at 1 mg/kg bodyweight. Following euthanasia of mice by $CO_2$ asphyxiation, thoracic cavities were opened and inferior vena cava cut to allow exsanguination, before injecting 7 ml of *EDTA buffer* le. Following perfusion, aortas were clamped using a hemostat and cut to extract hearts. The hemostat was subsequently placed on a 3D-printed platform (Online Figure 24) enabling hearts to be suspended above a waste fluid collection reservoir, with apex of the heart pointing upwards. To perfuse fluid, a hypodermic needle, connected to a peristaltic pump through the 3D-printed platform (Online Figure 24) which enabled it to be orientated above the heart, was inserted to the left ventricular chamber through the cardiac apex. Once the hearts were cannulated, *EDTA buffer* was pumped through using a peristaltic pump (~1ml/heart/min) until blood cleared (~5 min) before perfusion with *enzyme digest cocktail* for 15 minutes, during which time cardiac tissue was basted with fresh digestion buffer to prevent desiccation of the tissue. All buffers were maintained at 37°C during perfusion.

Following tissue digestion, cardiac ventricles were isolated by removing aorta and atria, and the ventricular digestion was finished by gently pulling apart the tissue before a 5-minute incubation at 37$^{o}$C followed by trituration with a 1000 µL pipette. The cell suspension was then passed through a 250 µM filter to remove large undigested debris and brought to 15 mL with cold *perfusion buffer* supplemented with 5% FBS in a conical tube. Three consecutive minute-long centrifugation sedimentations at 50*g*, 4$^{o}$C were performed to separate cardiomyocytes and non-myocytes based on cell mass. The supernatant from each sedimentation step was collected and pooled in a 50 mL tube which was then centrifuged at 400*g*, 4$^{o}$C for 5 minutes to pellet the non-myocyte fraction of cells. Whole intact non-myocyte cells then underwent antibody and viability dye staining before flow cytometry or fluorescence-activated cell sorting (FACS), as described previously [3].

**Nuclei extraction**—Cardiomyocyte nuclei were isolated by swelling cardiomyocytes in a hypotonic solution (91 mM NaCl, 5.3 mM KCl, 0.5 mM $MgCl_2$, 0.293 mM $CaCl_2$, 10 mM glucose, 10 mM HEPES at pH7.4) [4] before triturating cells using a 30-gauge hypodermic needle to mechanically fragment cells. Nuclei were then isolated by filtration through a 20-µm cell strainer (pluriStrainer) to further fragment cells and exclude large debris. Finally, cell fragments were centrifuged at 1000*g* for 5 minutes before staining with DAPI, followed by FACS for isolation of nuclei. For the proof-of-concept single-nucleus sequencing experiment (Online Figure 1), nuclei from both groups were isolated independently after cardiomyocyte and non-myocyte isolation via centrifugation and subsequently pooled at designated proportions.

**Single-cell transcriptomic library preparation and sequencing**—Intact non-myocyte cells and cardiomyocyte nuclei from each heart were mixed (mixing ratios for specific experiments indicated in Figure 1 and Online Figure 1) and used for single cell transcriptomic library preparation on the Chromium controller (10X Genomics). Approximately 12,000 total cells/nuclei were loaded into each reaction of the Chromium Single Cell 3' v2 reagent kit (10X Genomics). Following capture and lysis, we synthesized cDNA and amplified for 12 cycles as per manufacturer's protocol (10X Genomics). The amplified cDNA was used to construct

Illumina sequencing libraries that were each sequenced on one lane of an Illumina HiSeq 4000 to an approximate depth of 100,000 reads per cell. Single-cell isolation was performed in two batches on separate days containing at least n=2 from each condition to mitigate batch effects.

**Cardiac Tissue Assessment via Flow Cytometry**—Flow cytometric analysis of cardiac non-myocyte cell proportion and abundance was performed using the distinct cardiac tissue mincing protocol published previously (Pinto et al 2013, 2016). Two cell viability dyes were used, Sytox Green (ThermoFisher Scientific) and Calcein Blue (ThermoFisher Scientific). Antibodies used were MHCII (BD Biosciences clone clone:2G9, BUV395), Lyve1 (eBioscience clone:ALY7, eflour660), CD31 (BD Biosciences clone:390, BV605), CD45 (BioLegend clone:30-F11, APC-Cy7), CD64 (BioLegend clone:X54-5/7.1, PE-Cy7), Mcam (BD Biosciences clone:ME-9F1, BV711), CD3e (BD Biosciences clone:145-2C11, BV650), CD11b (BD Biosciences clone:m1/70, BUV737), and CD59a (BioLegend clone:mCD59.3, PE).

**Analysis of single-cell RNA-Seq data**—We used Cell Ranger version 2.1.1 (10X Genomics) to process raw sequencing data before subsequent analyses. This pipeline converted Illumina basecall files to fastq format, aligned sequencing reads to the mm10 transcriptome using the STAR aligner [5] and quantified the expression of transcripts in each cell. We carried out analyses of processed scRNA-seq data in R version 3.4 or 3.6 [6] using the Seurat suite versions 2.3.4 and 3.0.2 [7,8] and Tidyverse packages [9]. Single nuclei transcriptomes typically contain a higher fraction of reads mapping to introns than single cell transcriptomes [10]. Although retaining intronic reads in transcriptomes from cardiomyocytes would more deeply profile these nuclei, we chose to analyze only exonic reads for both nuclei and whole cell transcriptomes in order to treat data from each cell type equally. We obtained data from 29,682 cells that passed quality control steps implemented in Cell Ranger. As a further quality-control measure, we filtered out cells meeting any of the following criteria: <100 or >15,000 unique genes expressed, >50,000 UMIs, or >30% of reads mapping to mitochondria. These steps removed an additional 67 cells, resulting in a final dataset of 29,615 cells. We quantified gene expression across 27,998 genes and considered for further analysis a total of 17,170 genes that were expressed in at least ten cells in at least one of our eight samples.

To explore transcriptional heterogeneity and to undertake cell clustering, we reduced dimensionality using PCA. We selected 30 PCs that explained more variability than expected by chance using heuristics detailed in vignettes associated with the Seurat software. We used PC loadings as input for a graph-based approach to cluster cells by cell type where we clustered with resolution 1.2, and as input for t-distributed stochastic neighbor embedding (t-SNE) or fast interpolation-based t-SNE (FIt-SNE; [11]) for reduction to two dimensions for visualization purposes. For analyses of cells/nuclei isolated from wild-type homeostatic hearts, we merged transcriptionally similar clusters as previously describe [3]. For hexplot-style visualizations we used the schex package (https://github.com/SaskiaFreytag/schex).

**Differential expression analysis**—In order to identify differentially expressed (DE) genes, we first identified genes expressed in at least 10% of cells in at least one of the groups being compared. To test for differential expression we used MAST [12] including cellular detection rate as a covariate, which performed well in the single cell RNA-Seq differential expression benchmarking study of Soneson and Robinson [13]. We used a threshold of $p < 0.01$ to define genes as statistically significantly and differentially expressed between groups.

**Gene ontology analysis**—The Gene Ontology (GO) enrichment analysis for differentially expressed gene lists was performed using enrichDAVID function from clusterProfiler R package [14] which interacted with DAVID version 6.8 [15]. The Benjamini-Hochberg adjusted *p*-value cut-off of 0.05 was used to determine statistically significant GO terms.

**Ligand-receptor intercellular communication network analysis**—In order to represent potential intercellular communication between cardiac cell populations, we obtained mouse orthologs of human ligand-receptor pairs [16] from BioMart database version 86 using biomaRt R package [17]. A ligand/receptor with non-zero expression in more than 20% of cells in a particular cell population was deemed an "expressed" ligand/receptor. To construct the potential cell-cell communication network, we linked expressed ligands with their corresponding receptors between and within major cell populations. The signaling direction from ligand to receptor is indicated by arrows represented in the chord plot in Figure 6A. The total number of communication signals transmitted and received by a certain cell population is represented by the numbered bands in the circular visualization. We used the Circlize R package [18] to plot the putative intercellular communication network.

**Sex Analysis**—In order to examine sexually dimorphic gene expression patterns, we separated male and female cells based on expression of *Xist* and five Y chromosome genes (*Ddx3y, Eif2s3y, Gm29650, Kdm5d and Uty).* A cell with non-zero expression of *Xist* but zero expression of our Y chromosome genes was classified as a female cell, whereas a cell expressing one or more of the five Y chromosome genes with no expression of *Xist* was classified as a male cell. Because female cells should not contain a Y chromosome but male cells could express *Xist* transcript, we additionally classified cells with low or moderate *Xist* expression and without low expression of Y genes as male. Specifically, we considered cell library size-normalized read counts and classified cells with summed Y expression >1 UMI per thousand (which corresponded to cells not in the lowest 10% of Y gene expression among male cells called above) and *Xist* expression <8 UMI per thousand (which corresponded to cells with less than median *Xist* expression among female cells called above). Cells that expressed neither *Xist* nor Y chromosome genes, or cells that did not meet the criteria defined above, were removed from our sex analysis. We rescued ~75% of cells from our initial cell yield for the downstream sex analysis. This contained 15,181 female and 7,116 male cells (see Fig.7B). We examined sexual dimorphism in gene expression in Control and AngII-treated animals separately using methods for differential gene expression analysis described above.

**RNA velocity and pseudotime analyses**—To explore dynamic patterns of transcriptional changes among fibroblasts in resting and hypertrophic hearts, we isolated 15,446 fibroblasts from our complete dataset. We examined RNA velocity using Velocyto version 0.17.17 [19] to count spliced and unspliced reads and scVelo version 0.1.21 [20] to compute velocity. We ran scVelo with the arguments min_shared_counts=10 and n_top_genes=2000 to the function filter_and_normalize() and carried out analyses using 30 principal components. To identify genes that differed in velocity between Fibroblast-*Cilp* and Fibroblast-*Thbs4* compared to the remaining fibroblasts, we used the function rank_velocity_genes(). We also applied two pseudotime inference algorithms to all fibroblasts isolated from AngII-treated mice. We used slingshot version 1.3.2 [21] with default arguments and cells grouped according to clusters defined in our Seurat analysis (above). We used monocle3 version 0.1.1 [22], preprocessing with 15 principal components and regressing out percent mitochondrial reads as well as number of UMIs per cell.

**Analysis of Genotype Tissue Expression project (GTEx) RNA-Seq data**—We downloaded GTEx version 7 data from https://gtexportal.org/home/datasets (accessed 02/21/2019), including sample attributes, subject phenotypes, and gene TPMs from RNA-Seq. To examine gene expression in human hearts we focused on subjects within the age windows 40-49, 50-59, and 60-69 because these age windows contained greater than 25 samples per window. We classified the samples with highest 20% *NPPB* expression (without regard to age or sex) as putatively hypertrophic. We tested for differences in gene expression between putative hypertrophy/non-hypertrophy cohorts using linear models. Specifically, in our models the response variable was the logarithm of gene expression (TPM) and predictors included age class, sex, hypertrophy class, and sex×hypertrophy interaction, with the final term added because sexual dimorphism in response to hypertrophy was of particular interest. Results in the main text reporting differences in gene expression between hypertrophy classes were calculated with respect to the additive hypertrophy term with two-sided *p*-values.

### References related to Online Materials and Methods


[1] Pinto AR, Paolicelli R, Salimova E, Gospocic J, Slonimsky E, Bilbao-Cortes D, et al. An abundant tissue macrophage population in the adult murine heart with a distinct alternatively-activated macrophage profile. PLoS One 2012;7:e36814. doi:10.1371/journal.pone.0036814.

[2] Ackers-Johnson M, Li PY, Holmes AP, O'Brien SM, Pavlovic D, Foo RS. A Simplified, Langendorff-Free Method for Concomitant Isolation of Viable Cardiac Myocytes and Nonmyocytes from the Adult Mouse Heart. Circ Res 2016;119:909–920. doi:10.1161/CIRCRESAHA.116.309202.

[3] Skelly DADA, Squiers GTGT, McLellan MAMA, Bolisetty MTMTMT, Robson P, Rosenthal NANA, et al. Single-Cell Transcriptional Profiling Reveals Cellular Diversity and Intercommunication in the Mouse Heart. Cell Rep 2018;22:600–610. doi:10.1016/j.celrep.2017.12.072.

[4] Bell JR, Lloyd D, Curl CL, Delbridge LMD, Shattock MJ. Cell volume control in phospholemman (PLM) knockout mice: do cardiac myocytes demonstrate a regulatory volume decrease and is this influenced by deletion of PLM? Exp Physiol 2009;94:330–343. doi:10.1113/expphysiol.2008.045823.

[5] Dobin A, Davis CA, Schlesinger F, Drenkow J, Zaleski C, Jha S, et al. STAR: ultrafast universal RNA-seq aligner. Bioinformatics 2013;29:15–21. doi:10.1093/bioinformatics/bts635.

[6] R Core Team. R: A Language and Environment for Statistical Computing 2019.

[7] Butler A, Hoffman P, Smibert P, Papalexi E, Satija R. Integrating single-cell transcriptomic data across different conditions, technologies, and species. Nat Biotechnol 2018;36:411–420. doi:10.1038/nbt.4096.

[8] Stuart T, Butler A, Hoffman P, Hafemeister C, Papalexi E, Mauck WM, et al. Comprehensive Integration of Single-Cell Data. Cell 2019;177:1888-1902.e21. doi:10.1016/j.cell.2019.05.031.

[9] Wickham H. tidyverse: Easily Install and Load "Tidyverse" Packages 2017.

[10] Habib N, Avraham-Davidi I, Basu A, Burks T, Shekhar K, Hofree M, et al. Massively parallel single-nucleus RNA-seq with DroNc-seq. Nat Methods 2017;14:955–958. doi:10.1038/nmeth.4407.

[11] Linderman GC, Rachh M, Hoskins JG, Steinerberger S, Kluger Y. Fast interpolation-based t-SNE for improved visualization of single-cell RNA-seq data. Nat Methods 2019;16:243–245. doi:10.1038/s41592-018-0308-4.

[12] Finak G, McDavid A, Yajima M, Deng J, Gersuk V, Shalek AK, et al. MAST: a flexible statistical framework for assessing transcriptional changes and characterizing heterogeneity in single-cell RNA sequencing data. Genome Biol 2015;16:278.

doi:10.1186/s13059-015-0844-5.

[13] Soneson C, Robinson MD. Bias, robustness and scalability in single-cell differential expression analysis. Nat Methods 2018;15:255–261. doi:10.1038/nmeth.4612.

[14] Yu G, Wang LG, Han Y, He QY. ClusterProfiler: An R package for comparing biological themes among gene clusters. Omi A J Integr Biol 2012;16:284–287. doi:10.1089/omi.2011.0118.

[15] Huang DW, Sherman BT, Lempicki RA. Systematic and integrative analysis of large gene lists using DAVID bioinformatics resources. Nat Protoc 2009;4:44–57. doi:10.1038/nprot.2008.211.

[16] Ramilowski JA, Goldberg T, Harshbarger J, Kloppman E, Lizio M, Satagopam VP, et al. A draft network of ligand–receptor-mediated multicellular signalling in human. Nat Commun 2015;6:7866. doi:10.1038/ncomms8866.

[17] Durinck S, Spellman PT, Birney E, Huber W. Mapping identifiers for the integration of genomic datasets with the R/Bioconductor package biomaRt. Nat Protoc 2009;4:1184–1191. doi:10.1038/nprot.2009.97.

[18] Gu Z, Gu L, Eils R, Schlesner M, Brors B. Circlize implements and enhances circular visualization in R. Bioinformatics 2014;30:2811–2812. doi:10.1093/bioinformatics/btu393.

[19] La Manno G, Soldatov R, Zeisel A, Braun E, Hochgerner H, Petukhov V, et al. RNA velocity of single cells. Nature 2018;560:494–498. doi:10.1038/s41586-018-0414-6.

[20] Bergen V, Lange M, Peidli S, Wolf FA, Theis FJ. Generalizing RNA velocity to transient cell states through dynamical modeling. BioRxiv 2019:820936. doi:10.1101/820936.

[21] Cao J, Spielmann M, Qiu X, Huang X, Ibrahim DM, Hill AJ, et al. The single-cell transcriptional landscape of mammalian organogenesis. Nature 2019;566:496–502. doi:10.1038/s41586-019-0969-x.

[22] Street K, Risso D, Fletcher RB, Das D, Ngai J, Yosef N, et al. Slingshot: Cell lineage and pseudotime inference for single-cell transcriptomics. BMC Genomics 2018;19. doi:10.1186/s12864-018-4772-0.

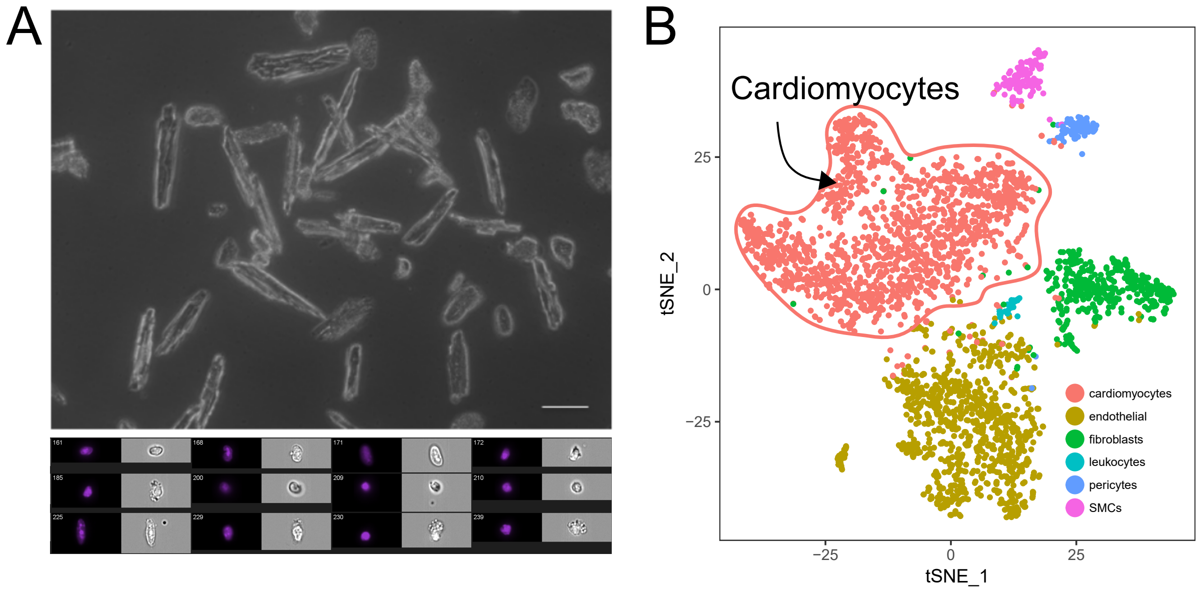


**Online Figure 1. Isolation and analysis of cardiac cell nuclei for scRNA-seq.**
A) Example images of isolated cardiomyocytes using the perfusion based cardiac single cell preparation method (see Online Materials and Methods for more detail) (top, phase contrast micrograph scale bar indicates 100 μm) and isolated cardiomyocyte nuclei imaged using the Amnis ImageStream system (bottom) showing DAPI and phase images of single nuclei. B) tSNE projection displaying cardiomyocyte nuclei (indicated) and non-myocyte nuclei analyzed by single-nucleus RNA-seq, as proof of concept for single-nucleus sequencing from live cardiac tissue (see Online Materials and Methods).

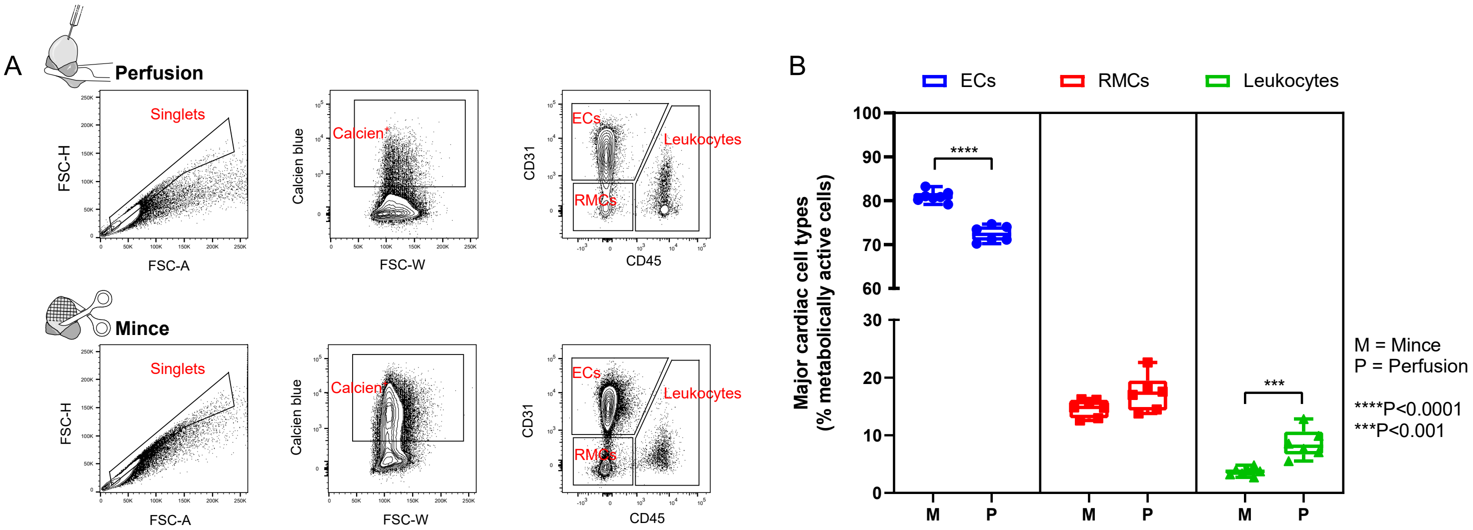


**Online Figure 2. Comparison of two cardiac cell isolation techniques on cellular composition and yield.** A) Flow cytometry gating strategy to identify relative proportions of endothelial cells (ECs), resident mesenchymal cells (RMCs) and leukocytes. Note: only non-myocytes are included in this analysis. B) Quantified proportions of three major cardiac cell types (% Calcein+ cells). Endothelial cell (EC) yield is significantly lower when performing the perfusion protocol, however significantly greater proportions of leukocytes are liberated by perfusion. Resident mesenchymal cell proportions are not altered by either protocol. Statistical analysis was conducted using an unpaired t-test. FSC-H = forward scatter height, FSC-W = forward scatter width, FSC-A = forward scatter area.

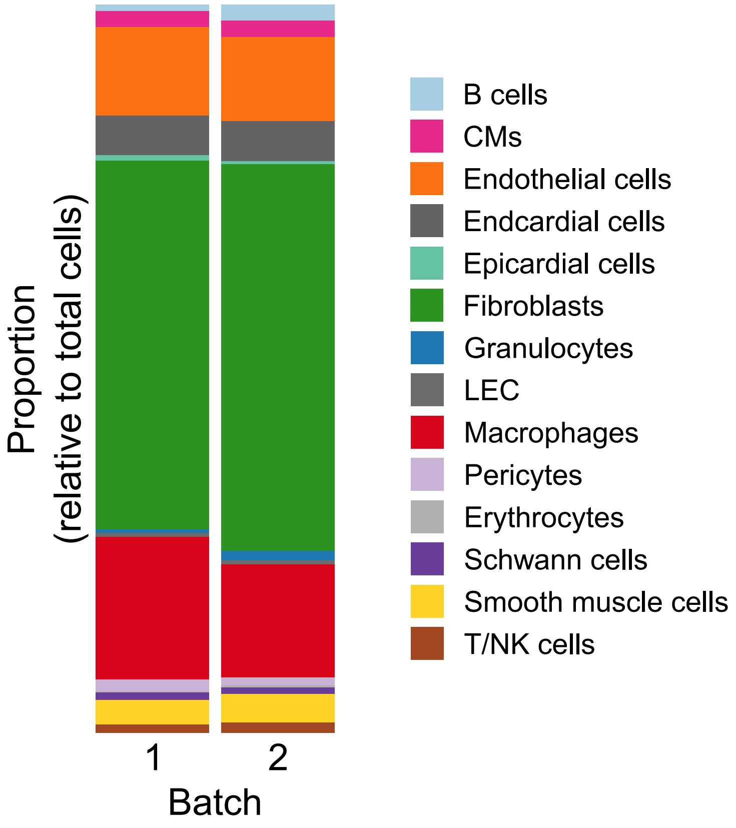


**Online Figure 3. Cellular composition of mouse hearts at homeostasis.**
Cellular composition of the sampling of sequenced cells derived from mouse hearts at homeostasis. Cells from two samples of wild type mice are shown in columns (batch). Each sample consisted of cells from female and male mice (Materials and Methods). Colors indicate cell populations and are identical to Figure 1B. Heights of individual boxes comprising each bar represent the proportion of cells classified as each cell population. Cellular composition shown here is not directly comparable to a homeostatic mouse heart due to the sorting strategy employed (Figure 1A).

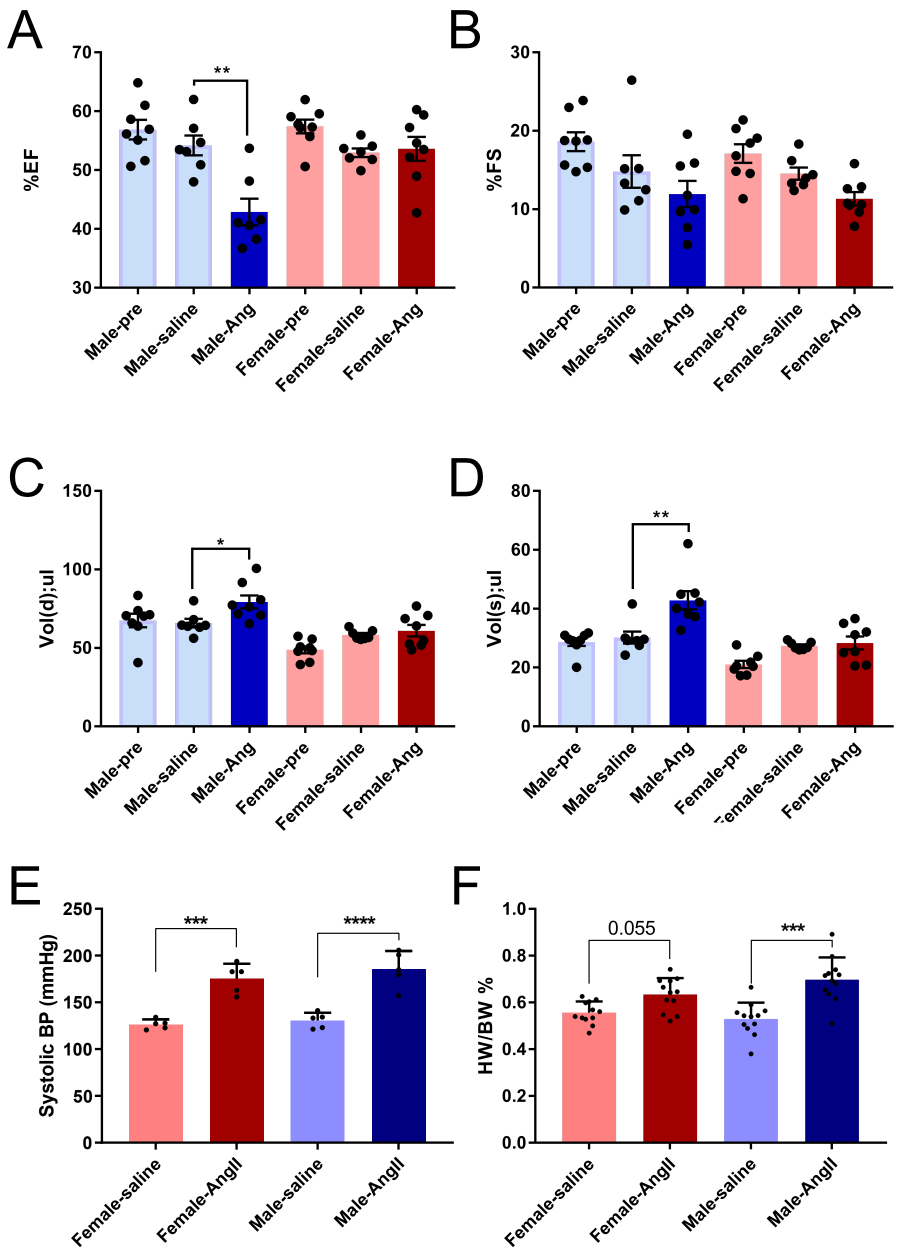


**Online Figure 4. Physiological assessment of cardiac function after AngII reveals sexual dimorphism in functional cardiac decline.**

A-D) Echocardiographic assessment of experimental mice prior to AngII stimulus (Male/Female-pre) and after two weeks AngII treatment, or saline control. Echocardiographs were used to calculate (A) ejection fraction percentage, (B) fractional shortening percentage, (C) diastolic volume, and (D) systolic volume. Statistical analysis was conducted using an unpaired t-test, *P< 0.05, **P< 0.005, ***P< 0.0005. E) Conscious blood pressure assessment after AngII, or saline control, in males and females. *P< 0.05, **P< 0.005, ***P< 0.0005. F) Heart-weight to body-weight ratio after AngII, or saline control, in males and females. ***P< 0.0005

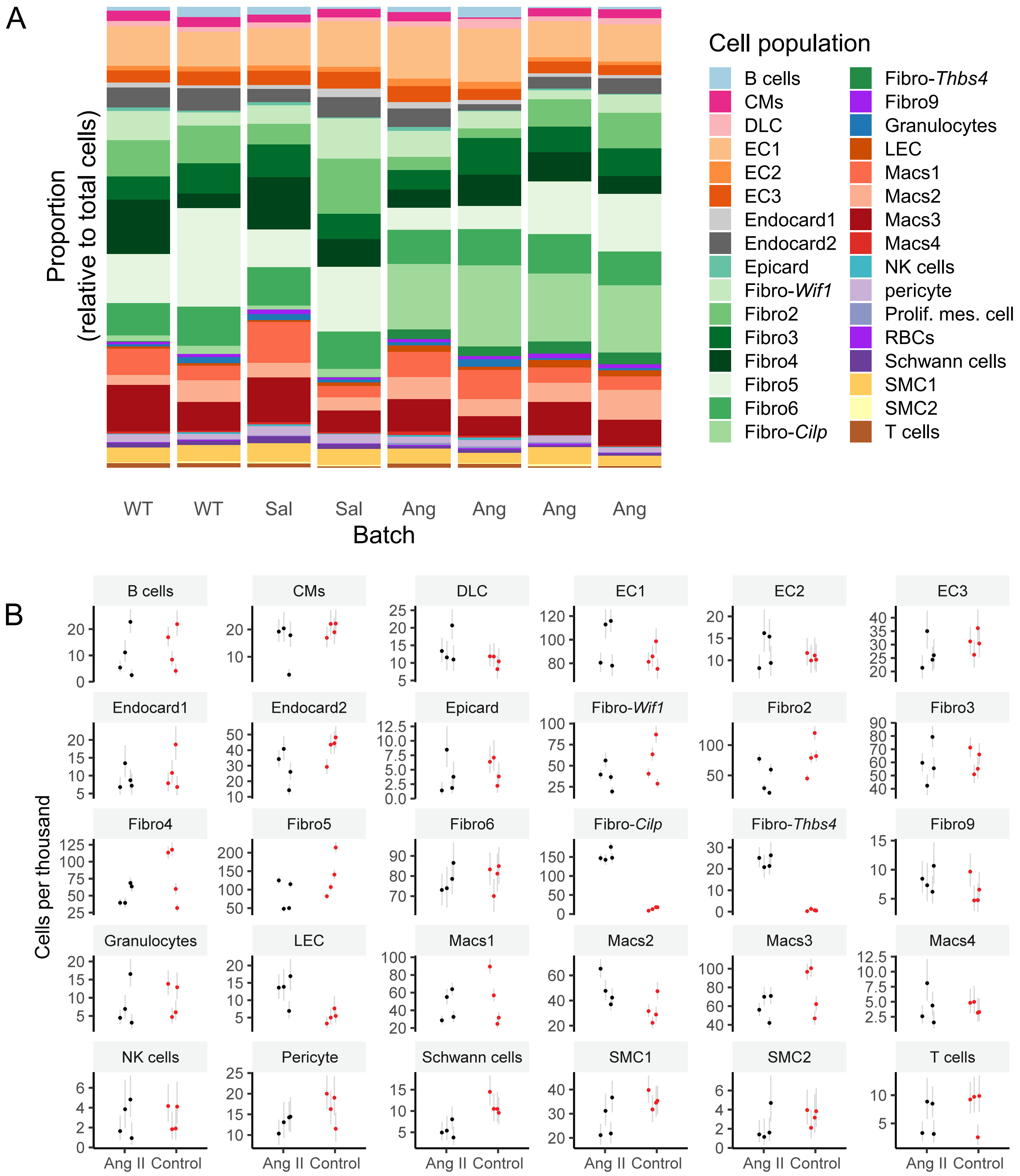


**Online Figure 5. Cellular composition of mouse hearts at homeostasis and after AngII treatment.**
A) Cellular composition of the sampling of sequenced cells derived from mouse hearts at homeostasis, after administration of saline (two weeks), and after administration of AngII (two weeks). Cells from individual samples are shown in columns (batch). Each sample consisted of cells from female and male mice (Materials and Methods). Colors indicate cell populations and sub-populations and are used throughout this study. Heights of individual boxes comprising each bar represent the proportion of cells classified as each cell population. Cellular composition shown here is not directly comparative to hearts isolated from treated and untreated mice due to the sorting strategy employed (Online Figure 1A). B) As in (A) but showing individual points representing observed cell abundances (in units of cells per thousand sequenced) of each population. Individual AngII-treated or control (wild type or saline administration) samples mice are indicated by black and red dots, respectively. Light grey lines indicate binomial proportion confidence intervals and are calculated using Jeffreys interval.

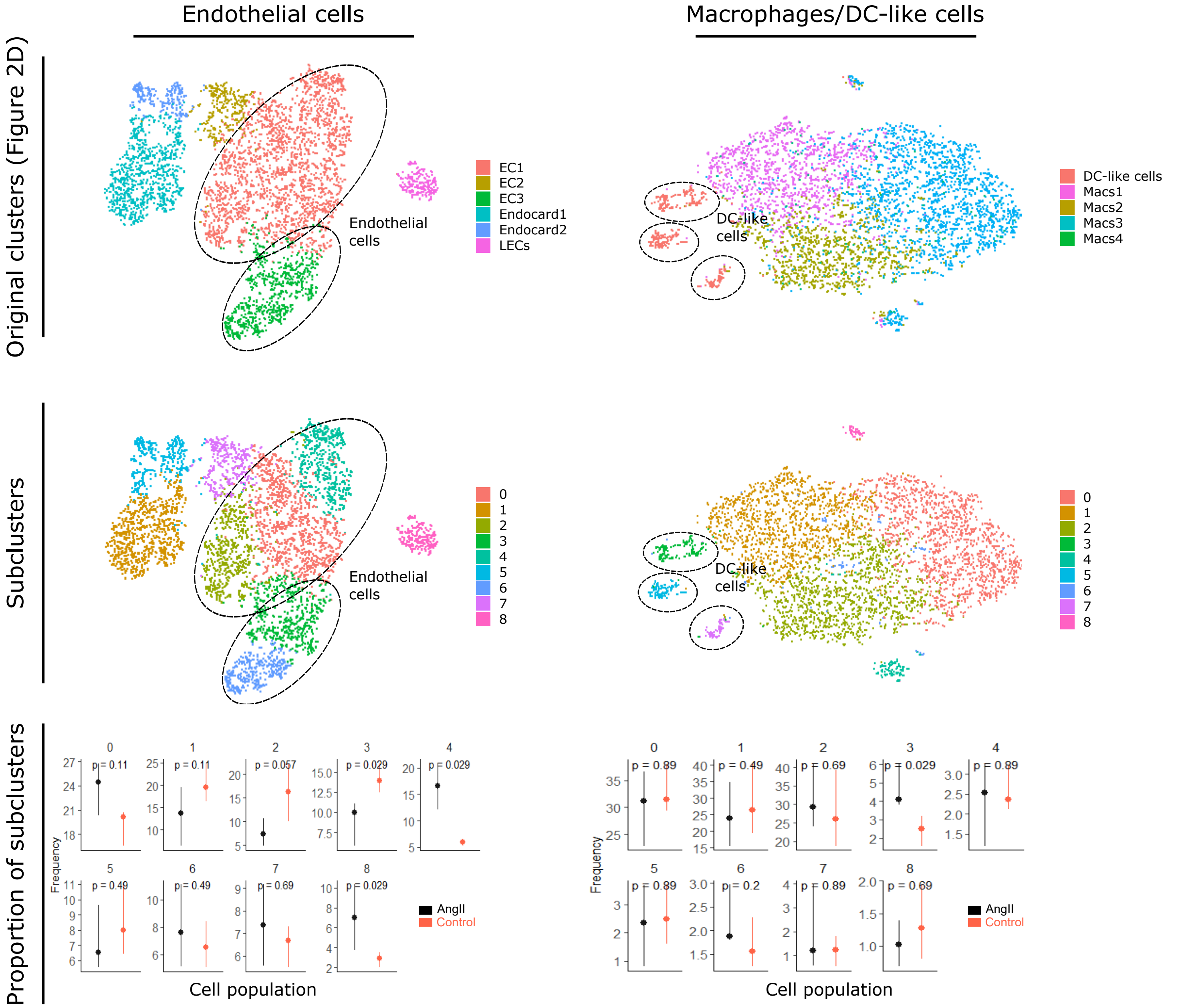


**Online Figure 6. Sub-clustering of endothelial cells and macrophages.**
Top two tSNE projections display endothelial cell and macrophage/DC-like cell populations (left and right, respectively) with cells colored based on clustering shown on Figure 2D. Bottom two tSNE projections display endothelial cells and macrophage populations (left and right, respectively) with cells colored based on new clusters identified following re-clustering. Dot-and-whisker plots summarize relative abundances (in percentage) of new clusters identified following re-clustering of cell populations. The cluster each plot corresponds to is indicated by the number above each plot. Statistical analysis was performed using Wilcoxon-test.

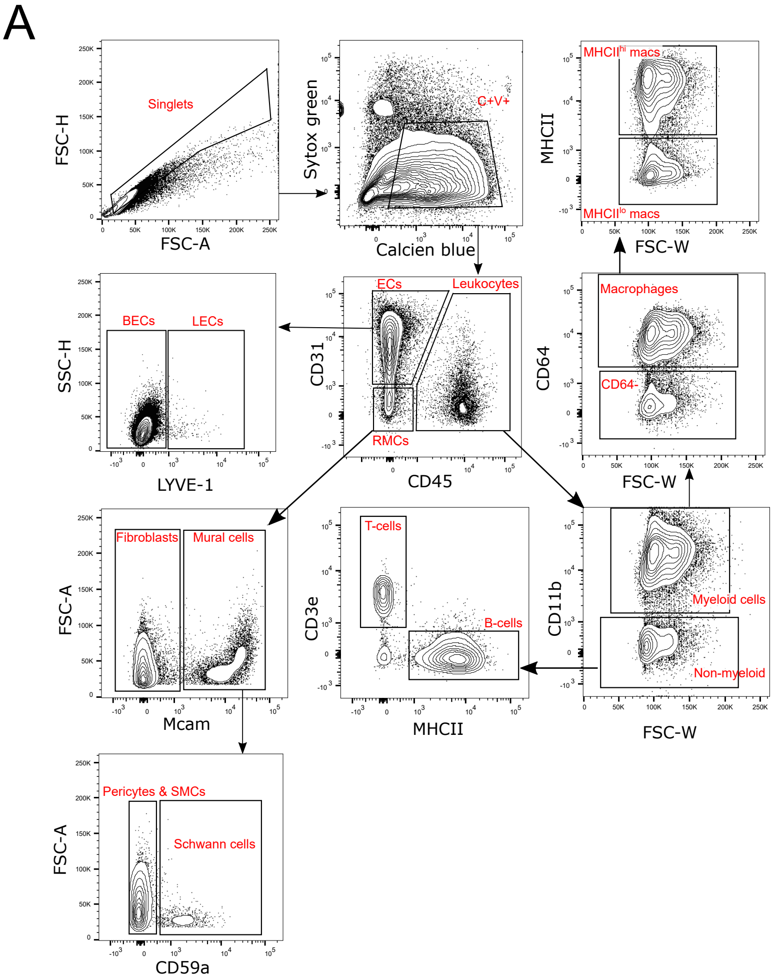


**Online Figure 7. Sexual dimorphism in cardiac cellular composition changes in response to AngII detected by flow cytometry**
A) Flow cytometry gating strategies to identify relative proportions of live, metabolically active (C+V+) cardiac cell types. B) Quantified proportions of each cell type normalized to the mean of each WT control (WT = 1). ECs were significantly elevated in male mice given AngII compared to females given AngII. Leukocytes were also significantly increased in male AngII mice compared to male WT controls within the major cardiac cell compartment. Both vascular and lymphatic ECs are elevated in male AngII mice compared to male WT controls and female AngII counterparts. No changes were observed within the resident mesenchymal cell compartment. However, in the leukocyte compartment male AngII mice exhibited significantly elevated myeloid cells compared to both male WT mice as well as female AngII mice. Consequently, these aforementioned increases in leukocyte proportions were observed in macrophages including their polarization sub-types MHCIIhi and MHCIIlo. Statistical analysis was conducted using an unpaired t-test, where significance was determined as P< 0.05. FSC-H = forward scatter height, FSC-W = forward scatter width, FSC-A = forward scatter area, SSC-H = side scatter height.

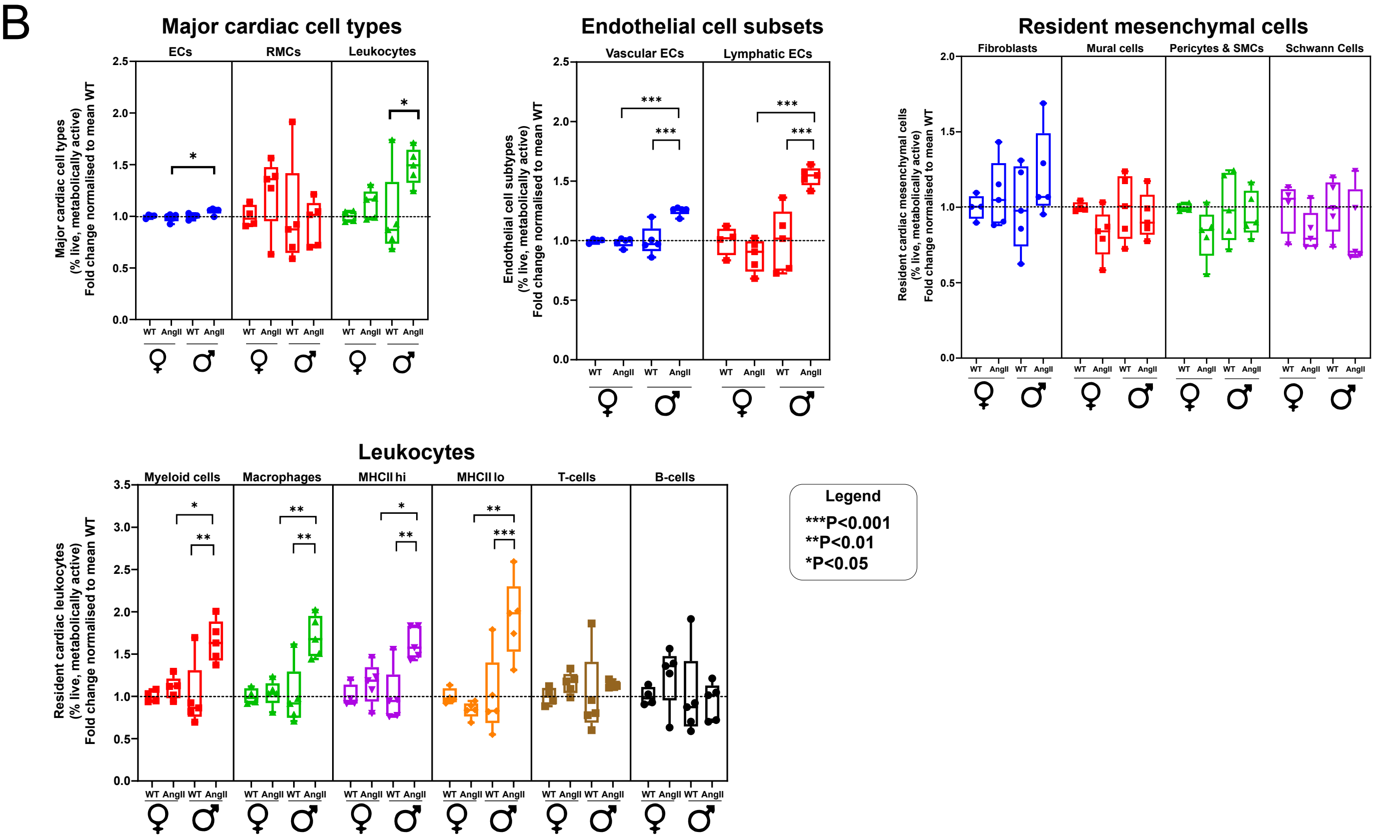

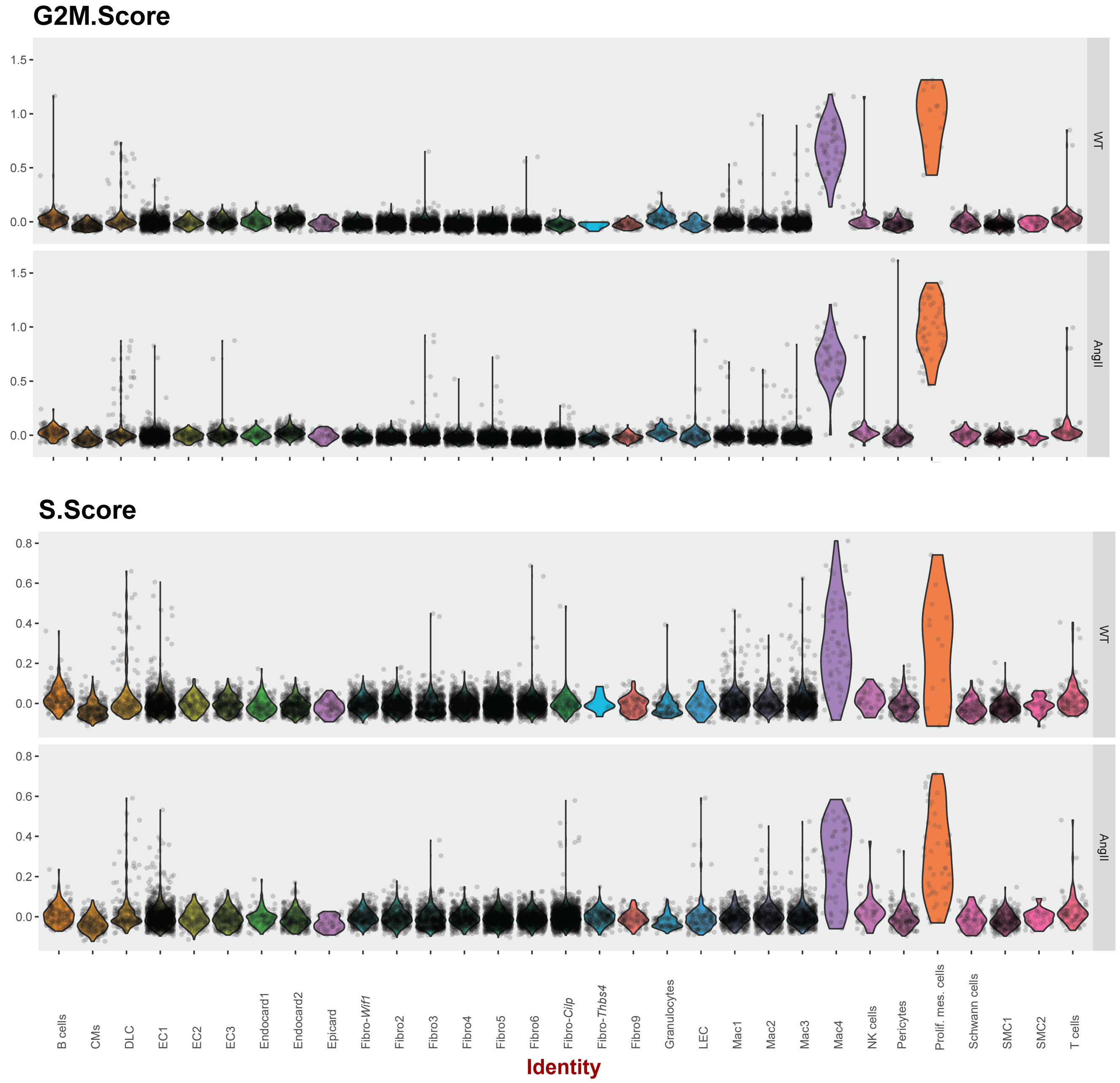


**Online Figure 8. Cell cycle scoring to identify dividing cardiac cells from mice with and without AngII treatment.**
Cell cycle scores are determined by considering expression levels G2/M and S phase marker genes and is based on the strategy by Tirosh et al., 2016 [10].

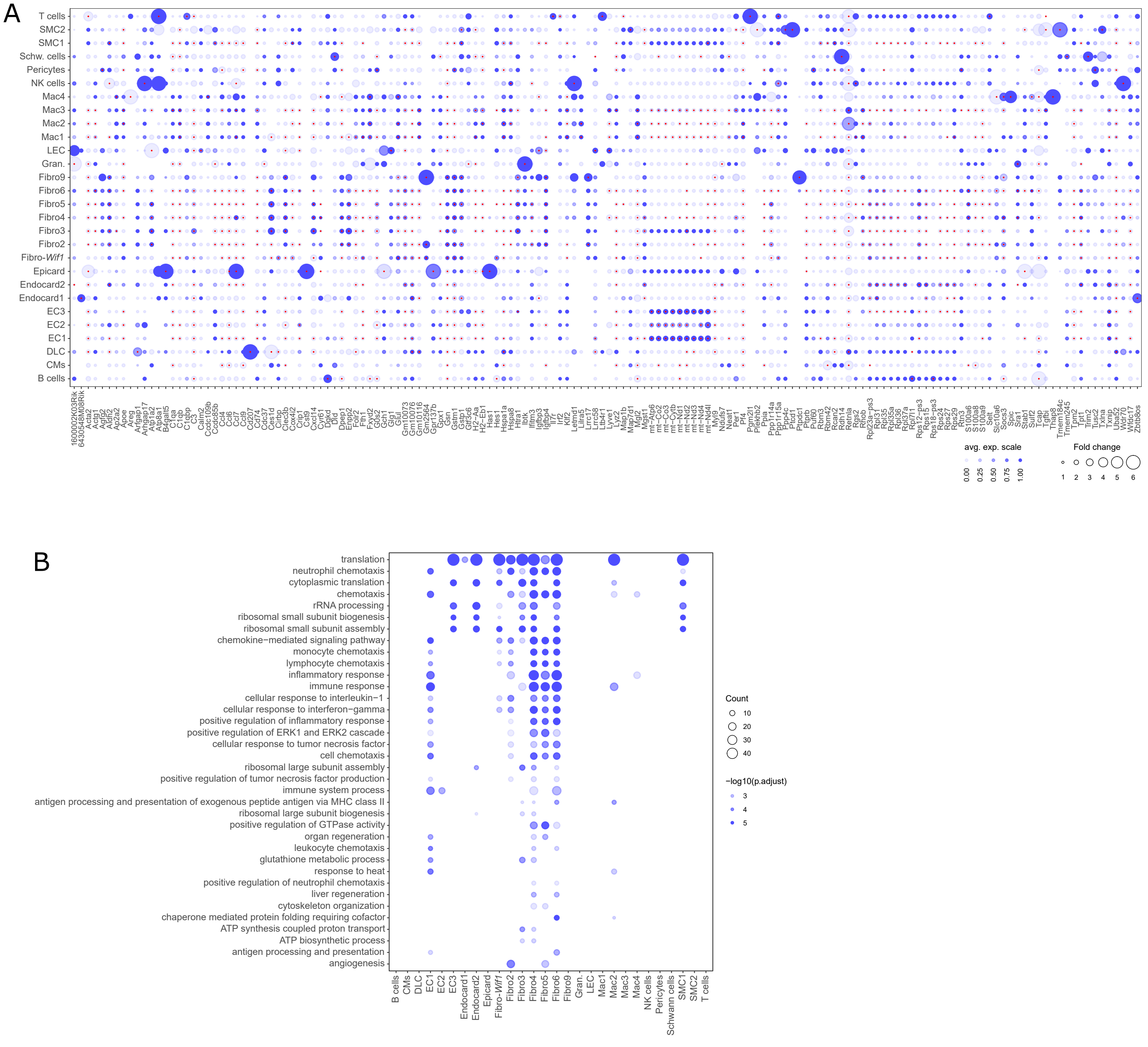


**Online Figure 9. Downregulated genes after induction of fibrosis.**

A) Dot plot summarizing the top 10 downregulated genes for each cell population. Duplicate entries of genes have been removed. Fold change (FC) in gene expression is indicated by circle size. Circle color indicates relative expression level in cells from control samples (expression level before reduction). Red dots indicate reduction in gene expression where p≤0.01 (uncorrected). B) Enrichment of GO terms in downregulated genes. Number of genes downregulated (circle size) and adjusted p-value (circle color) are indicated.

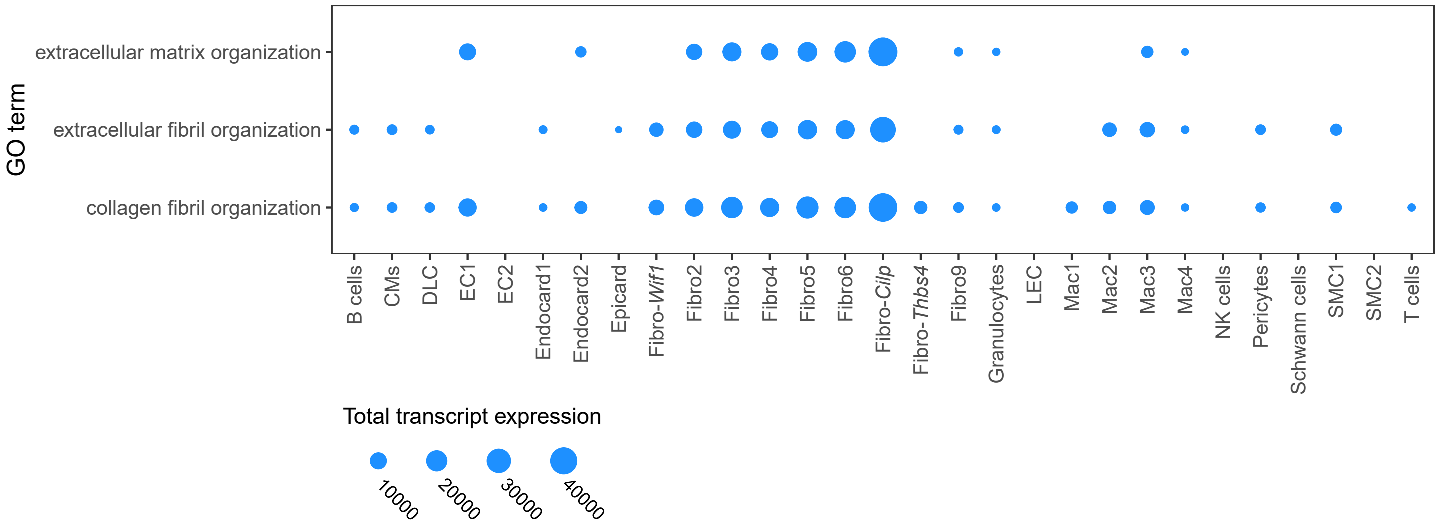


**Online Figure 10. Total transcript corresponding to ECM-related GO terms.**
Dot plot showing expression of genes classified within the GO categories shown on y-axis. Circle size represents the sum of transcripts for genes corresponding to each GO term within each cell population.

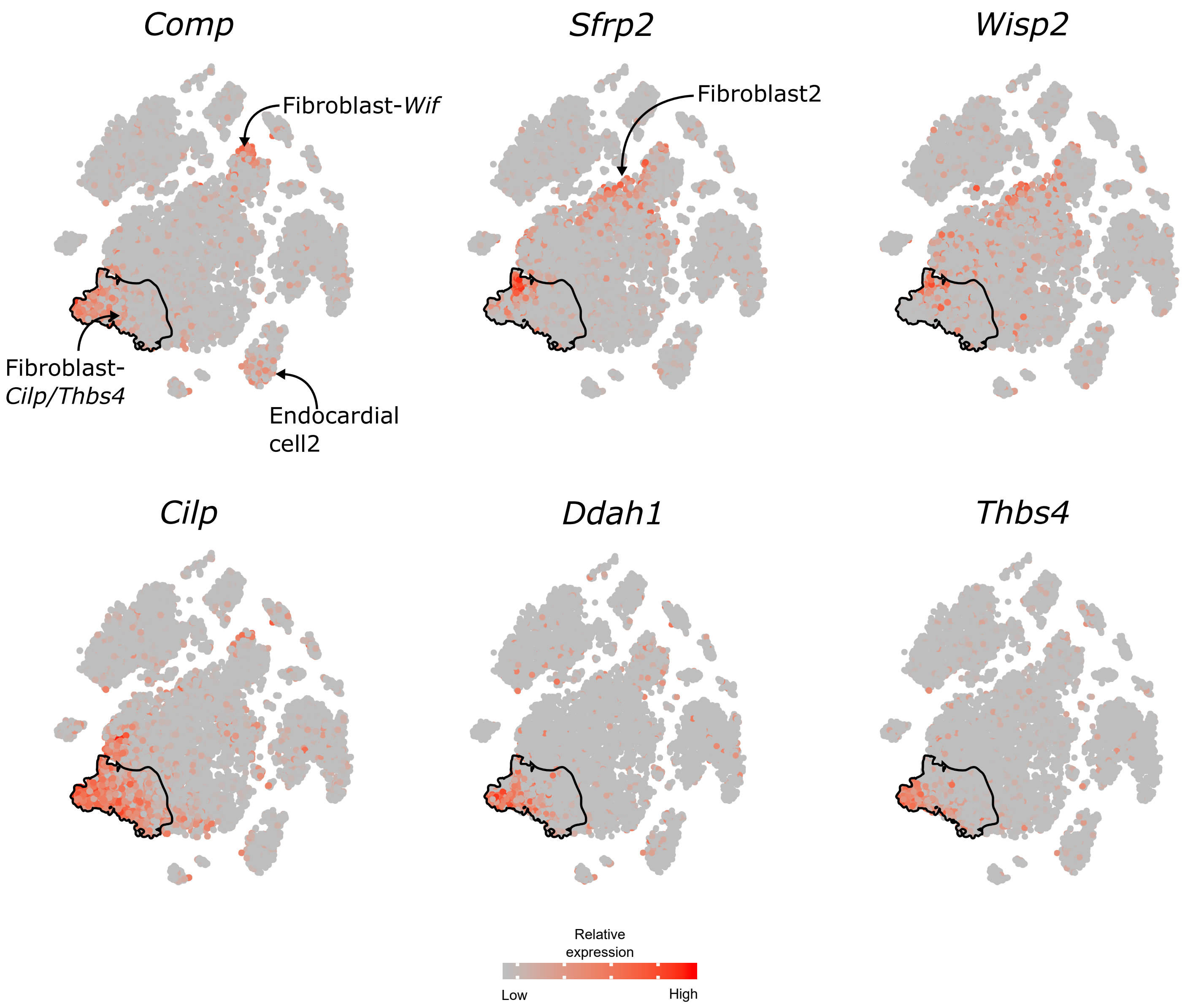


**Online Figure 11. Markers of matrifibrocytes and Fibroblast-Cilp and Fibroblast-Thbs4.**
Figures show FItSNE projections for cardiac cell populations (as shown in Figure 2D) with relative gene expression indicated by color (red=high, gray=low).

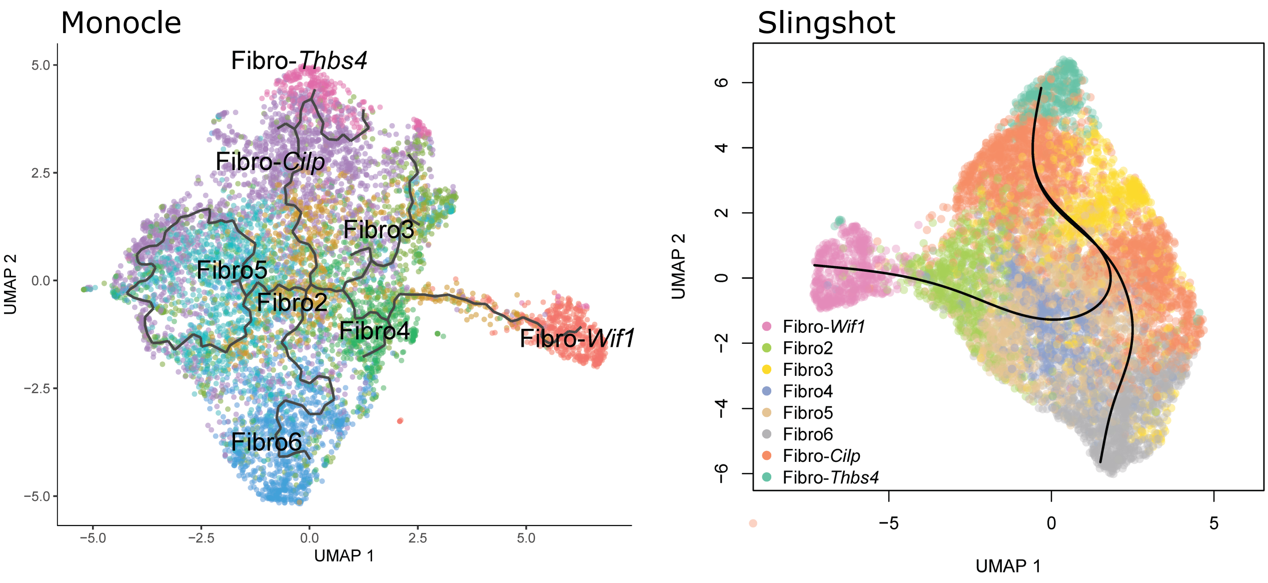


**Online Figure 12. Shifts in fibroblast cell state in response to chronic stress imposed by AngII treatment.**

Left- Pseudotime trajectories of fibroblasts isolated from AngII-treated mice inferred using monocle3. Cells are visualized using UMAP dimensionality reduction and colored according to fibroblast cluster. Gray line indicates inferred pseudotime trajectory. Right- Pseudotime trajectories of fibroblasts isolated from AngII-treated mice inferred using slingshot. Cells are visualized using UMAP dimensionality reduction and colored according to fibroblast cluster. Black line indicates inferred pseudotime trajectory.

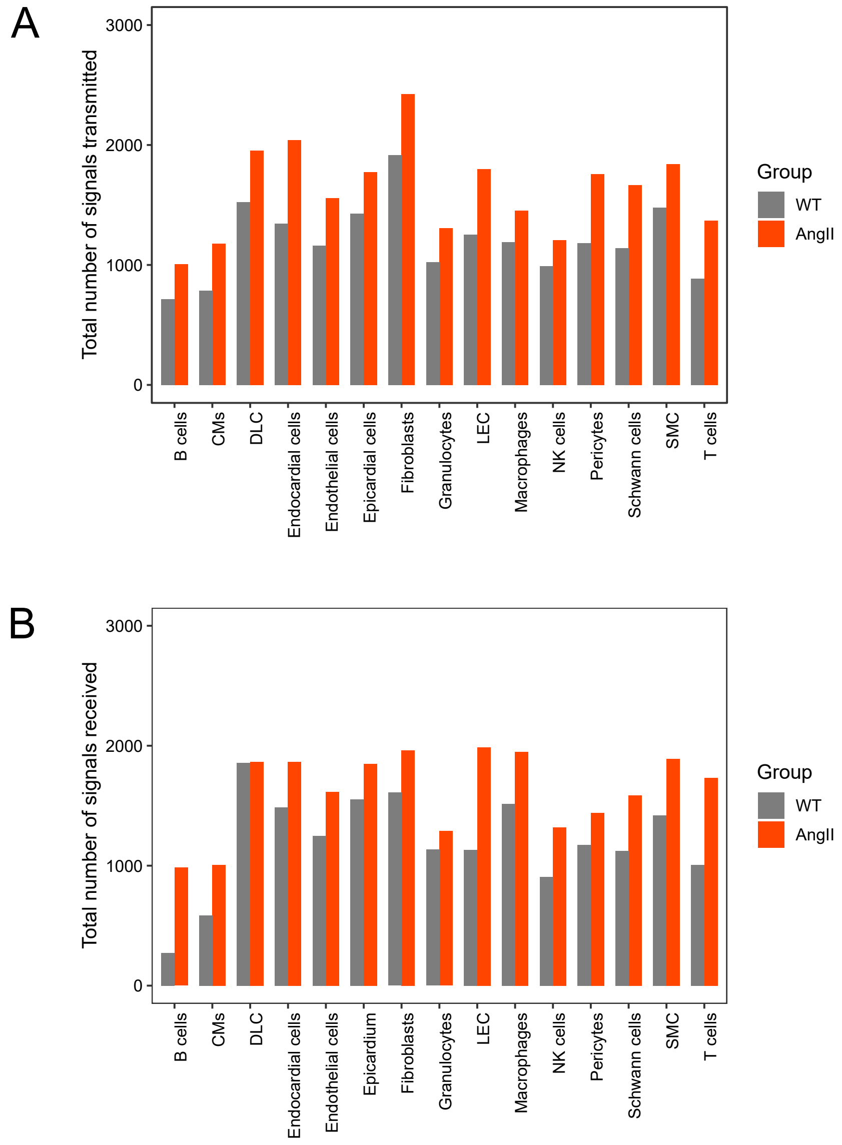


**Online Figure 13. Transmitted and received signals by cardiac cells with and without AngII treatment.**
A) Number of signals transmitted as determined by ligand encoding genes which are expressed by cells which have cognate receptors expressed by other cardiac cell populations. B) Number of signals received as determined by receptor encoding genes expressed by cells which have cognate ligands expressed by other cardiac cell populations.

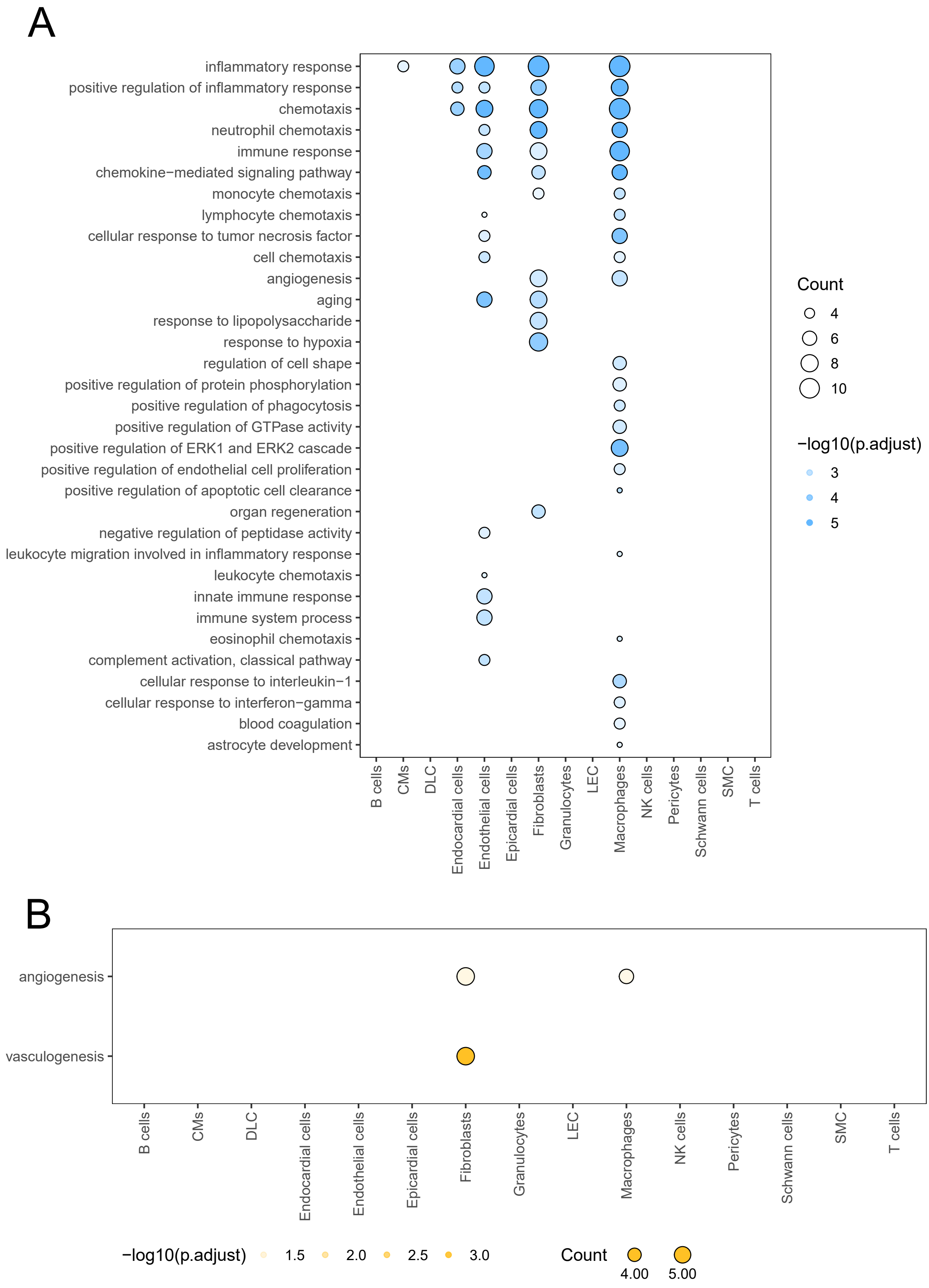


**Online Figure 14. GO terms corresponding to downregulated ligand and receptor genes.**
A) GO terms enriched in downregulated ligand genes. B) GO terms enriched in downregulated receptor genes.

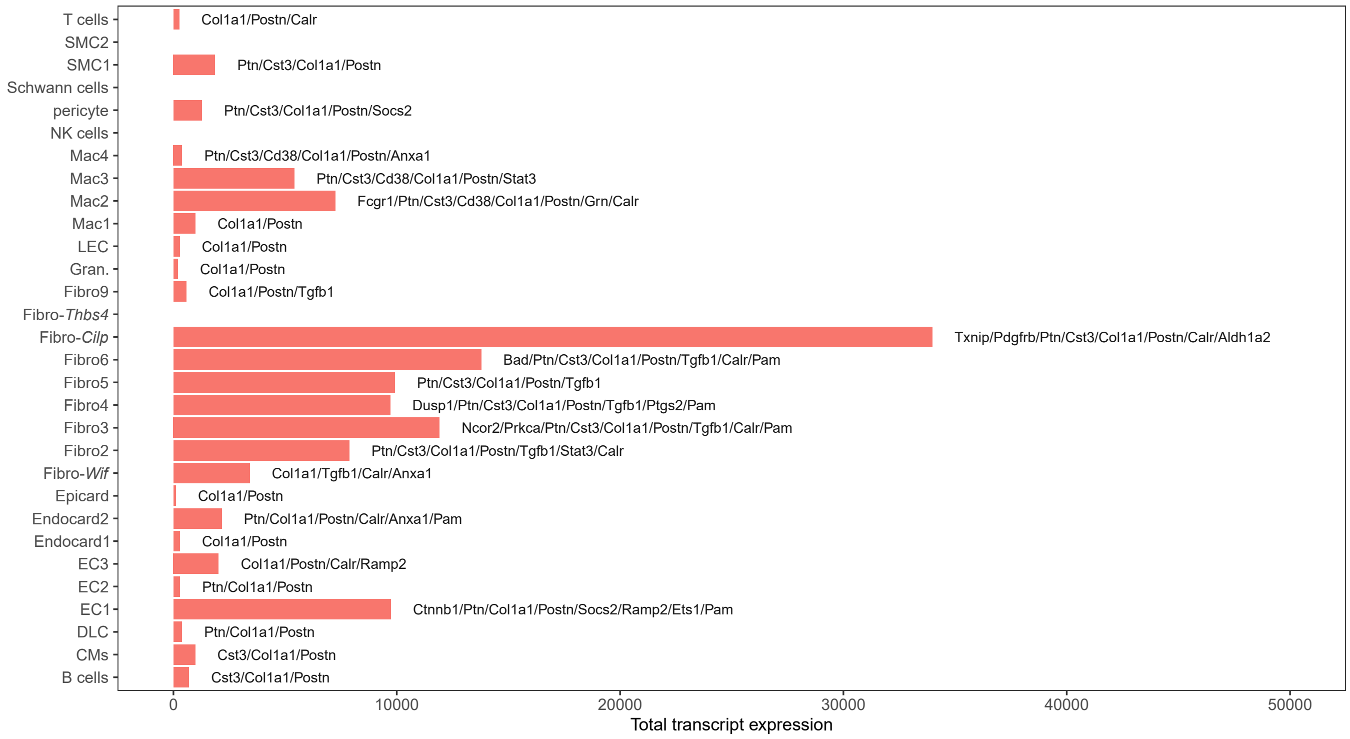


**Online Figure 15. Summed transcript levels of genes annotated as responsive to estradiol that are upregulated in cardiac cells following AngII treatment**

A bar plot which summarizes the total transcript abundance for genes that are upregulated in cardiac cells following AngII treatment and included in the 'response to estradiol' GO term.

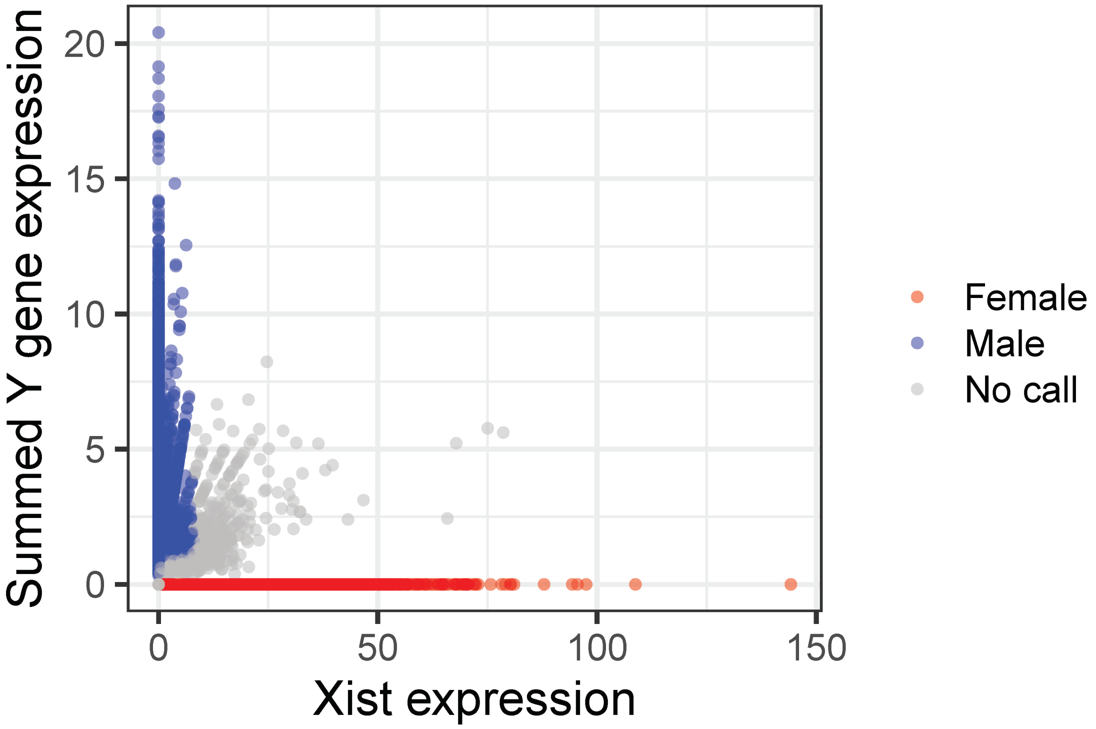


**Online Figure 16. Distribution of males and female cells based on expression of Y chromosome-related genes and Xist.**
Blue and red dots indicate male and female cells, respectively. Grey dots represent cells which could not be discriminated. Genes considered are outside of the pseudoautosomal regions of the X and Y chromosomes (see Online Materials and Methods).

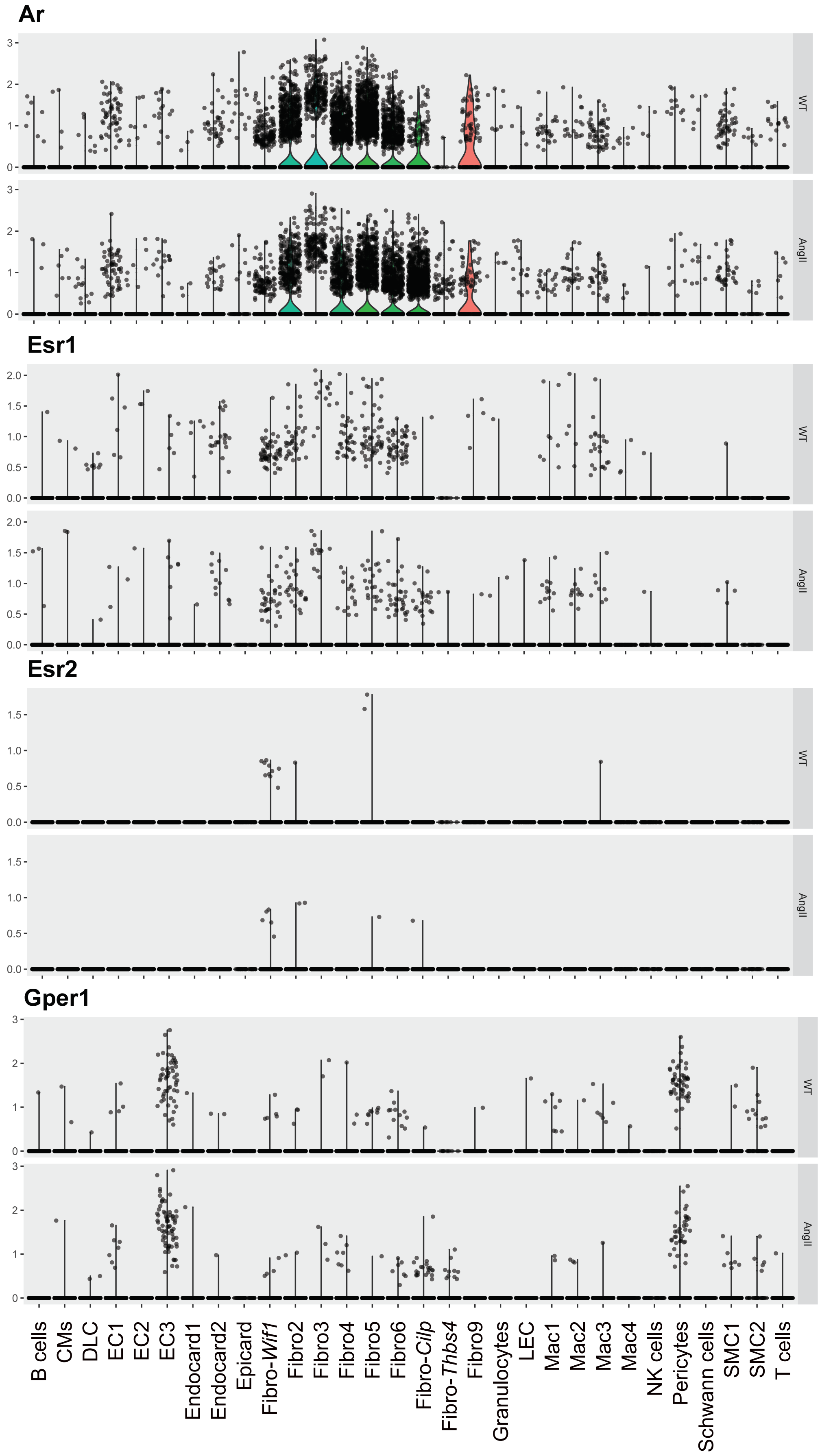


**Online Figure 17. Sex hormone receptors in cardiac cells from mice with and without AngII treatment.** Violin plots generated with Seurat showing the expression of various hormone receptors across all cardiac cell types. Androgen receptor expression (Ar) is greatest in cardiac fibroblasts. Estrogen receptor α and estrogen receptor β (Esr1, Esr2, respectively) are both lowly expressed, but most highly detected in cardiac fibroblasts. The membrane bound G protein-coupled estrogen receptor 1 (Gper1) is also lowly expressed, but greatest in endothelial cells and pericytes.

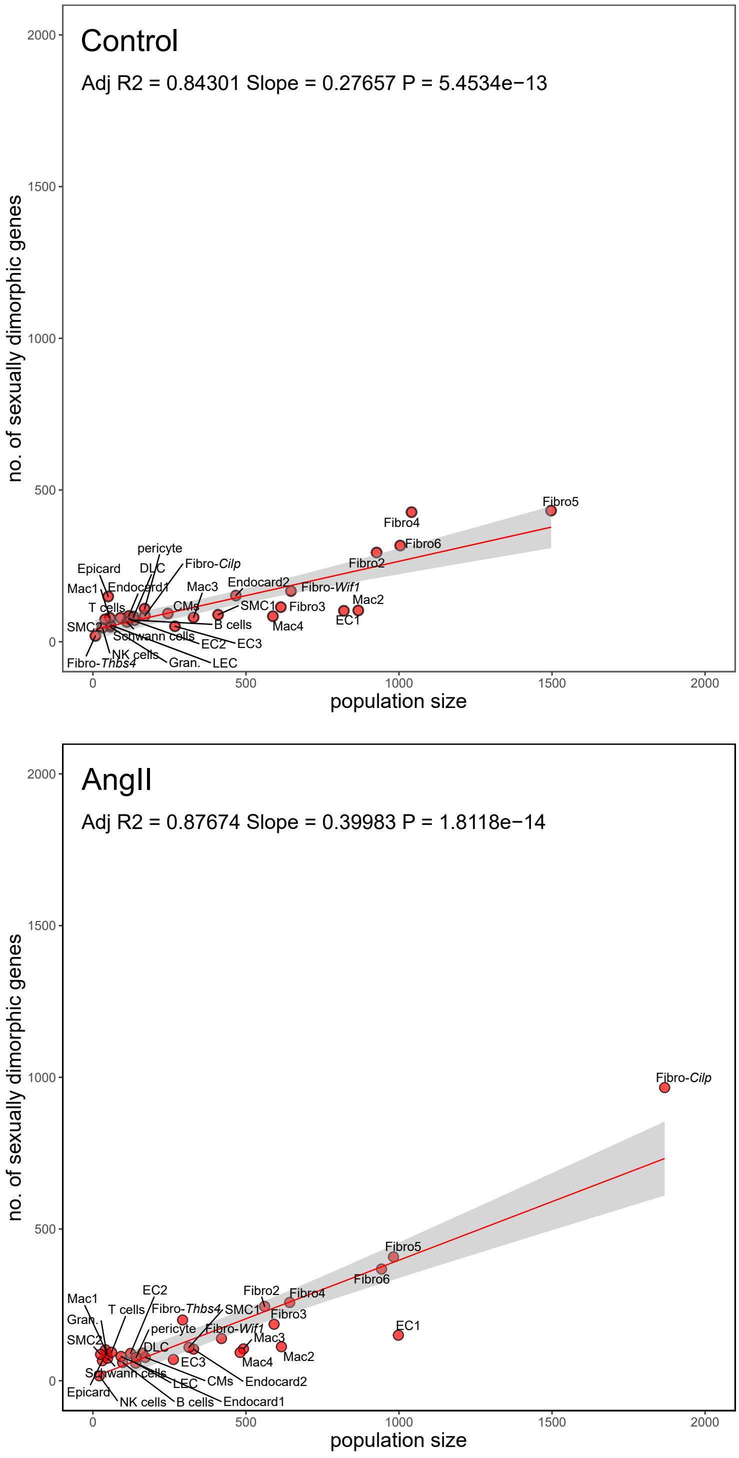


**Online Figure 18. Relationship between number of sexually dimorphic genes discovered and cell population size.**

Top panel summarizes the relationship in cardiac cell populations from control animals. Bottom panel summarizes the relationship in cardiac cell populations from AngII-treated animals.

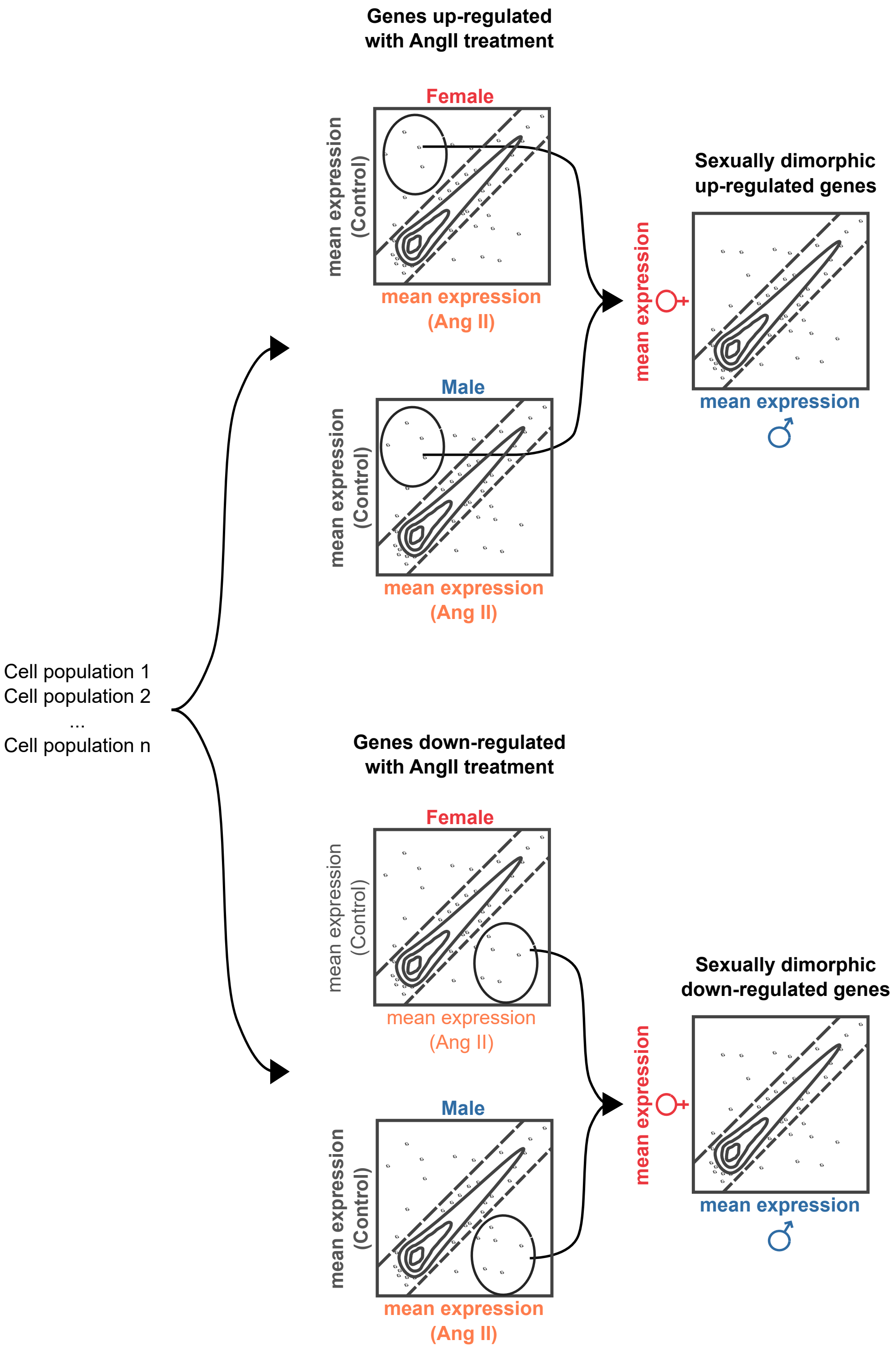


**Online Figure 19. Strategy for determining sexually dimorphic genes regulation following AngII treatment (related to figure 7D-F).**

For each cell population, female and male cells were considered in isolation and genes upregulated or downregulated following AngII treatment determined. For dimorphisms in upregulated genes (results shown Figure 7D-F), a list of genes upregulated in either females or males was generated and sexual dimorphisms in gene expression was examined. For dimorphisms in downregulated genes (results shown Online Figure 20), a list of genes downregulated in either sex was generated and dimorphism in expression was again examined.

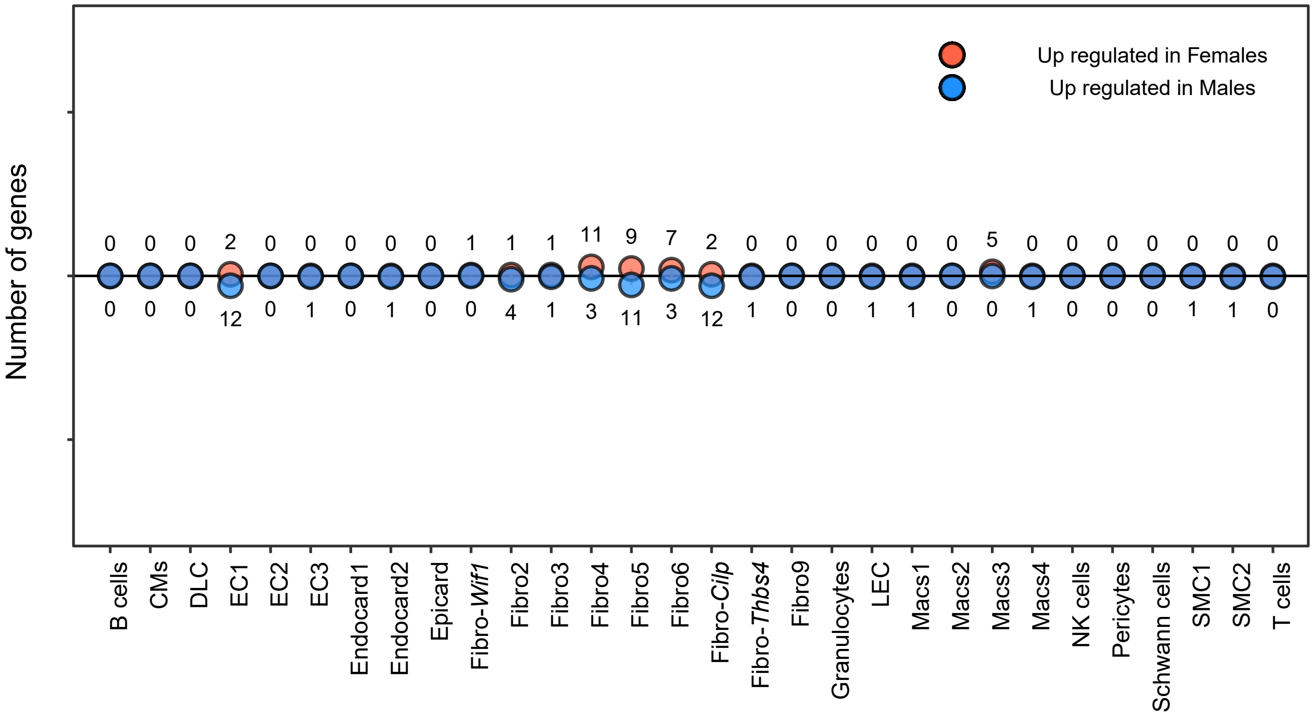


**Online Figure 20. Sexual dimorphism in gene expression amongst genes that are downregulated in cardiac cells following AngII treatment.**

The number of genes which are downregulated in response to AngII as well as exhibiting sexually dimorphic gene expression, presented for each cardiac cell type. In this context upregulated refers to a gene that is greater in either sex, but overall remains lower after AngII. Strategy to determine sexually dimorphisms is outlined in Online Figure 19.

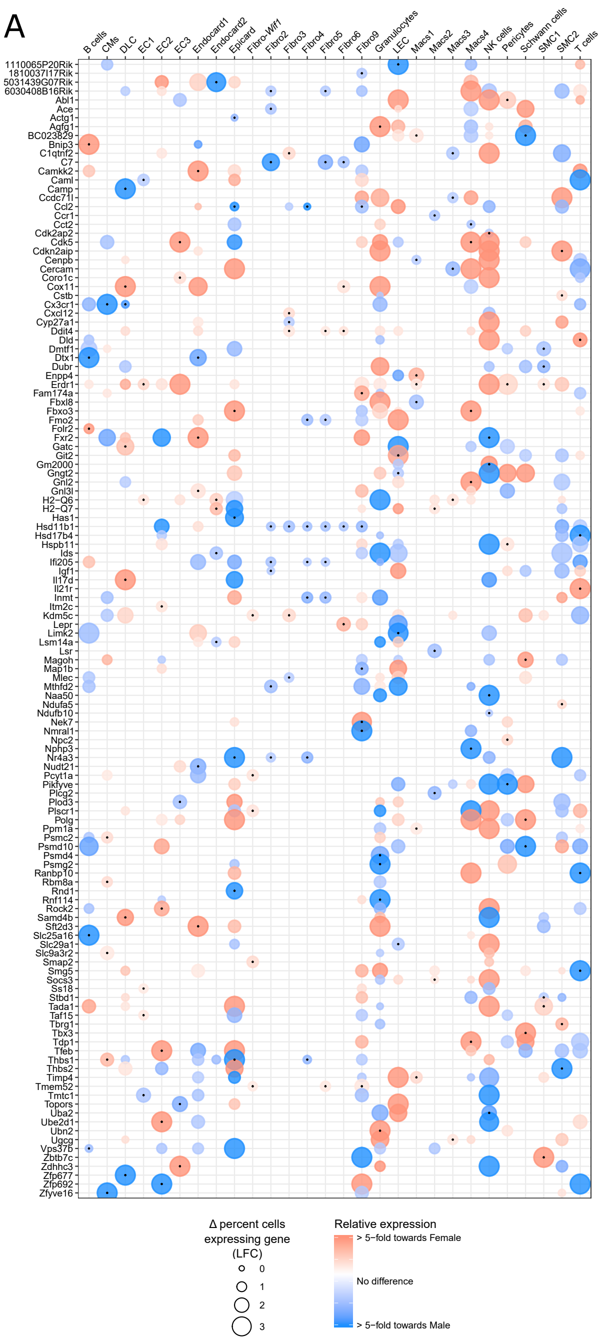


B

| GO ID | Term | p.adj | Genes | Cell population |
|---|---|---|---|---|
| GO:0032870 | cellular response to hormone stimulus | 2.0314E-05 | Jun, Fosb, Fos, Aif1, Dusp1, Jund, Igfbp7, Junb | |
| GO:0051591 | response to cAMP | 0.00986339 | Jun, Fosb, Fos, Dusp1, Jund, Junb | |
| GO:0071277 | cellular response to calcium ion | 0.00795899 | Jun, Fosb, Fos, Jund, Cacybp, Junb | |
| GO:0050873 | brown fat cell differentiation | 0.01707746 | Bnip3, Arl4a, Selenbp1, Fabp4, Cebpa | Cardiomyocytes |
| GO:0042493 | response to drug | 0.01406265 | Jun, ND1, Fosb, Fos, Abca1, Jund, Dpysl2, Hsp90aa1, Lpl, Lgals1, Junb | |
| GO:0006911 | phagocytosis, engulfment | 0.03434078 | Aif1, Gsn, Abca1, Fcgr2b, Mfge8 | |
| GO:0006366 | transcription from RNA polymerase II promoter | 0.0396789 | Jun, Fosb, Fos, Jund, Junb, Egr1, Gtf2h1 | |
| GO:0002479 | antigen processing and presentation of exogenous peptide antigen via MHC class I, TAP-dependent | 0.01633121 | Psma1, H2-Q7, Psmb9, H2-K1, B2m | Endothelial cell 2 |
| GO:0002474 | antigen processing and presentation of peptide antigen via MHC class I | 0.01710388 | H2-T22, H2-Q6, H2-Q7, H2-K1, B2m | |
| GO:0006412 | translation | 1.1114E-38 | Rpl9, Rpl6, Rpl14, Rpl35, Eef1d, Rpl39, Eif4e, Rpl37a, Rps16, Rpl11, Rpl35a, Mrps24, Rpl41, Rps24, Rps29, Rpl23a, Rps8, Eif3g, Rps14, Fau, Rpl22l1, Rps15a, Rps5, Rps28, Rpl13a, Rpl19, Aimp1, Rpl32, Rps27rt, Rps19, Rps9, Rpl5, Eif3h, Rpl37, Eif3k, Rps23, Rpl38, Rps18, Rpl34, Rpl27a, Rps27a, Rps21, Rps2, Eif3f, Rpl17, Rpl18a, Rpl18, Rps6, Rpl8, Fars2, Rpl36a | |
| GO:0000028 | ribosomal small subunit assembly | 1.3894E-05 | Rps5, Rps28, Rps27rt, Rps25, Rps19, Rps14, Rps2 | |
| GO:0042274 | ribosomal small subunit biogenesis | 0.00021328 | Rps28, Rps6, Rps19, Rps24, Rps16, Nob1 | |
| GO:0002474 | antigen processing and presentation of peptide antigen via MHC class I | 0.00022106 | H2-D1, H2-T22, H2-Q4, H2-Q6, H2-Q7, H2-K1, B2m | |
| GO:0006364 | rRNA processing | 0.00057952 | Rpl11, Rpl35a, Lsm6, Rps28, Rpl14, Rps6, Rpl5, Rps19, Rps24, Rps16 | |
| GO:0002479 | antigen processing and presentation of exogenous peptide antigen via MHC class I, TAP-dependent | 0.00119835 | H2-D1, H2-Q7, Psma7, Psmb9, H2-K1, B2m | Endocardial cell 2 |
| GO:0030490 | maturation of SSU-rRNA | 0.00154051 | Lsm6, Rps28, Rps19, Rps14, Nob1 | |
| GO:0002181 | cytoplasmic translation | 0.00170038 | Rplp2, Rpl22l1, Rpl35a, Rpl9, Rpl6, Rpl8 | |
| GO:0001731 | formation of translation preinitiation complex | 0.00775607 | Eif3f, Eif3g, Eif3k, Eif3h, Rps2 | |
| GO:0000462 | maturation of SSU-rRNA from tricistronic rRNA transcript (SSU-rRNA, 5.8S rRNA, LSU-rRNA) | 0.01102751 | Rps8, Rps19, Rps14, Rps24, Rps16 | |
| GO:0019882 | antigen processing and presentation | 0.0119613 | H2-D1, Cd74, H2-DMb1, Psmb9, H2-K1, H2-Eb1 | |
| GO:0032870 | cellular response to hormone stimulus | 0.00753557 | Fos, Dusp1, Jund, Socs2, Igfbp7, Junb, Ramp2 | |
| GO:0006412 | translation | 0.01710525 | Slc25a39, Rps8, Rpl27, Rpl13, Rps3, Rpl27a, Rpl37a, Rpsa, Rps5, Rps28, Rpl18a, Rps3a1, Rps19, Eif3k | Macrophage 2 |
| GO:0002479 | antigen processing and presentation of exogenous peptide antigen via MHC class I, TAP-dependent | 0.04050329 | H2-D1, Psma2, H2-Q7, Psmb8, B2m | |
| GO:0002376 | immune system process | 0.00095306 | H2-DMa, Il4ra, H2-Q7, Arhgef2, Apobec3, H2-Eb1, H2-K1, Hmgb2, Rnf19b, H2-D1, Casp4, Cd74, Irf7, H2-Ab1, C3, B2m, Irf1 | |
| GO:0019882 | antigen processing and presentation | 0.00768406 | H2-D1, H2-DMa, Cd74, H2-Ab1, H2-K1, H2-Eb1, Rab6a | Macrophage 4 |
| GO:0002474 | antigen processing and presentation of peptide antigen via MHC class I | 0.0093927 | H2-D1, H2-Q4, H2-Q6, H2-Q7, H2-K1, B2m | |
| GO:0006457 | protein folding | 4.8066E-07 | Ahsa1, Fkbp4, Hspa8, Dnaja4, Hsp90ab1, Hspe1, Cct2, Dnaja1, Cdc37l1, Hspa4l, Hsp90aa1, Mkks, Nudcd2, Hspd1 | |
| GO:0006986 | response to unfolded protein | 0.00036995 | Hspa1a, Hspa4l, Hsp90aa1, Derl1, Hspb1, Hsph1, Hspe1, Hspd1 | |
| GO:0009408 | response to heat | 0.00053588 | Hspa1a, Hsf1, Hspa1b, Hsp90aa1, Dnaja4, CYTB, Hspd1, Dnaja1 | Endothelial cell 2 |
| GO:0042773 | ATP synthesis coupled electron transport | 0.00529547 | ND5, ND4L, ND4, COX2 | |
| GO:0042026 | protein refolding | 0.01730404 | Hsp90aa1, Hspa8, Dnaja4, Hspd1 | |
| GO:0051085 | chaperone mediated protein folding requiring cofactor | 0.04697373 | Hspa8, Hsph1, Hspe1, Hspd1 | |
| GO:0030198 | extracellular matrix organization | 2.9821E-05 | Lamb2, Lama2, Vwa1, Col4a1, Nid1, Ccdc80, Crispld2, Kazald1, App, Eln, Lamc1, Tnxb | |
| GO:0006986 | response to unfolded protein | 2.8114E-05 | Manf, Herpud1, Hspa4l, Dnajc3, Thbs1, Hsp90aa1, Hspb1, Derl2, Hspe1 | |
| GO:0030199 | collagen fibril organization | 0.00110549 | Col1a2, Lox, P4ha1, Cyp1b1, Serpinh1, Col5a1, Tnxb | |
| GO:0007155 | cell adhesion | 0.00310122 | Lama2, Nid1, Itgb5, App, Fblim1, Thbs1, Cyp1b1, Svep1, Plpp3, Ret, Lamb2, Prkca, Spon1, Tnfaip6, Ccn2, Srpx2, Lamc1, Col5a1 | |
| GO:0070374 | positive regulation of ERK1 and ERK2 cascade | 0.00600562 | Ccl11, Ccl19, Pdgfrb, Prkca, Bmper, F2r, Ccn2, Gm2023, Alkal2, C3, Pten | |
| GO:0010466 | negative regulation of peptidase activity | 0.00567305 | Timp2, Ecm1, Serpinb6a, App, Cst3, Serpine2, Timp3, Serpina3n, Serping1 | |
| GO:0045766 | positive regulation of angiogenesis | 0.00619782 | Ccl11, Ecm1, Prkca, Thbs1, F3, Cyp1b1, Ptgis, Hspb1, C3 | Fibroblast 2 |
| GO:0001525 | angiogenesis | 0.0056342 | Ramp1, Angptl4, Col4a1, Thsd7a, Ccdc80, Arhgap24, Ecm1, Prkca, Ccn2, Srpx2, Cyp1b1, Pten | |
| GO:0006457 | protein folding | 0.00721002 | Fkbp4, Hspa4l, Hsp90aa1, Hspa8, Nktr, Hsp90b1, Hsp90ab1, Hspe1, Dnaja1 | |
| GO:0032355 | response to estradiol | 0.0101001 | Txnip, Penk, Pdgfrb, Prkca, Cst3, Ccnd1, Ccn2, Pten | |
| GO:0034976 | response to endoplasmic reticulum stress | 0.01472181 | Atp2a2, Herpud1, Thbs1, Sel1l, Srpx, Sdf2l1, Hspa5 | |
| GO:0050921 | positive regulation of chemotaxis | 0.04329284 | Ccl19, Pdgfrb, Thbs1, Gm2023 | |
| GO:0045793 | positive regulation of cell size | 0.04896771 | Hsp90aa1, Ret, Kdm1a, Hsp90ab1 | |
| GO:0030970 | retrograde protein transport, ER to cytosol | 0.04896771 | Herpud1, Hsp90b1, Sel1l, Derl2 | |
| GO:0030198 | extracellular matrix organization | 0.00026242 | Lgals3, Nid1, Ccdc80, Crispld2, Comp, Nid2, Vit, Lamc1, Fbln1, Smarca4, Tnxb | |
| GO:0006986 | response to unfolded protein | 0.00043678 | Manf, Herpud1, Hspa4l, Dnajc3, Hsp90aa1, Derl1, Hspb1, Hsph1 | Fibroblast 5 |
| GO:0034976 | response to endoplasmic reticulum stress | 0.00445175 | Ppp2cb, Atp2a2, Herpud1, Srpx, Txndc11, Sdf2l1, Hyou1, Hspa5 | |
| GO:0045766 | positive regulation of angiogenesis | 6.494E-05 | Ccl11, Zc3h12a, Ecm1, Hmox1, Sphk1, Xbp1, Thbs1, F3, Ptgis, Hspb1, Serpine1, C3 | |
| GO:0006986 | response to unfolded protein | 0.0006216 | Syvn1, Manf, Herpud1, Dnajc3, Xbp1, Thbs1, Hsp90aa1, Hspb1 | |
| GO:0034976 | response to endoplasmic reticulum stress | 0.00052992 | Atp2a2, Herpud1, Xbp1, Thbs1, Sel1l, Srpx, Sdf2l1, Hyou1, Hspa5 | |
| GO:0030335 | positive regulation of cell migration | 0.00043512 | Ccl11, Xbp1, Sphk1, Tgfb1, Thbs1, Itga5, Hspa5, Csf1, Tnfaip6, F2r, Lamc2, Hras, Ptp4a1 | |
| GO:0071347 | cellular response to interleukin-1 | 0.00047236 | Ccl11, Zc3h12a, Cxcl2, Ccl7, Nfkb1, Ptgis, Myc, Ccl2, Serpine1 | |
| GO:0006954 | inflammatory response | 0.00060819 | Ccl11, Map2k3, Zc3h12a, Fas, Sphk1, Tgfb1, Thbs1, Rel, Cxcl2, Ecm1, Ccl7, Csf1, F2r, Ccl2, C3, Ptgs2 | |
| GO:0030970 | retrograde protein transport, ER to cytosol | 0.00392432 | Syvn1, Herpud1, Hsp90b1, Sel1l, Selenos | |
| GO:0001525 | angiogenesis | 0.00586329 | Ramp1, Zc3h12a, Col4a1, Ccdc80, Ecm1, Tnfrsf12a, Hmox1, Ccn2, Xbp1, Ccl2, Serpine1, Ptgs2 | |
| GO:0001934 | positive regulation of protein phosphorylation | 0.0174311 | Plaur, Gprc5b, Ccn2, Sphk1, Xbp1, Tgfb1, Fnip1, Hras, Sqstm1, C3 | |
| GO:0032967 | positive regulation of collagen biosynthetic process | 0.01881991 | F2r, Suco, Ccn2, Tgfb1, Ccl2 | Fibroblast 6 |
| GO:0071260 | cellular response to mechanical stimulus | 0.01864462 | Bcl10, Hdac4, Fas, Gja1, Tgfb1, Nfkb1, Ptgs2 | |
| GO:0034097 | response to cytokine | 0.02118437 | Timp2, Rel, Skil, Nfkb1, Serpina3n, Srf, Ptgs2 | |
| GO:0032496 | response to lipopolysaccharide | 0.02073098 | Mgst1, Nfkbia, Penk, Cxcl2, F2r, Fas, Ace, Mta1, Trib1, Ptgs2 | |
| GO:0010628 | positive regulation of gene expression | 0.02297981 | Zc3h12a, Lmna, Plaur, Vim, Tgfb1, Hspa8, Nfkb1, Serpine1, Osr1, Csf1, Ccn2, Gja1, Hras, Mafg | |
| GO:0010629 | negative regulation of gene expression | 0.03723729 | Zc3h12a, Rel, Slit3, Ccn2, Gja1, Tgfb1, Ace, Nfkb1, Hras, Myc, Serpine1 | |
| GO:0006950 | response to stress | 0.03933165 | Eif2ak1, Hsp90aa1, Hsp90b1, Hsp90ab1, Map4k4 | |
| GO:0050729 | positive regulation of inflammatory response | 0.03907691 | Ccl11, Ccl7, Gprc5b, Ace, Ccl2, Serpine1 | |
| GO:0034599 | cellular response to oxidative stress | 0.03907691 | Srxn1, Zc3h12a, Atp2a2, Penk, Xbp1, Selenos | |
| GO:0008285 | negative regulation of cell proliferation | 0.04565623 | Eif2ak1, Timp2, Tgfb1, Srf, Adarb1, Hdac4, Hmox1, F2r, Slit3, Gja1, Serpine2, Hras, Ptgs2 | |
| GO:0050729 | positive regulation of inflammatory response | 0.04577001 | S100a9, Ctss, Ccl4, Fabp4, S100a8, Hspd1, Il33 | Granulocytes |

**Online Figure 21. Sexually dimorphic genes in cell populations from control mouse hearts.**
A) A dot plot showing genes differentially expressed at 2-fold or greater between males and females. Red and blue circles represent genes which are more highly expressed in females or males respectively. Circle size represents the difference in the percentage of cells expressing the gene between sexes. Solid black dots indicate a statistically significant difference in expression with p≤0.01 (unadjusted). B) GO terms enriched in sexually dimorphic gene sets in female and male cardiac cell populations. Red and blue text indicate GO terms enriched in sets of genes that are upregulated in females and males, respectively.

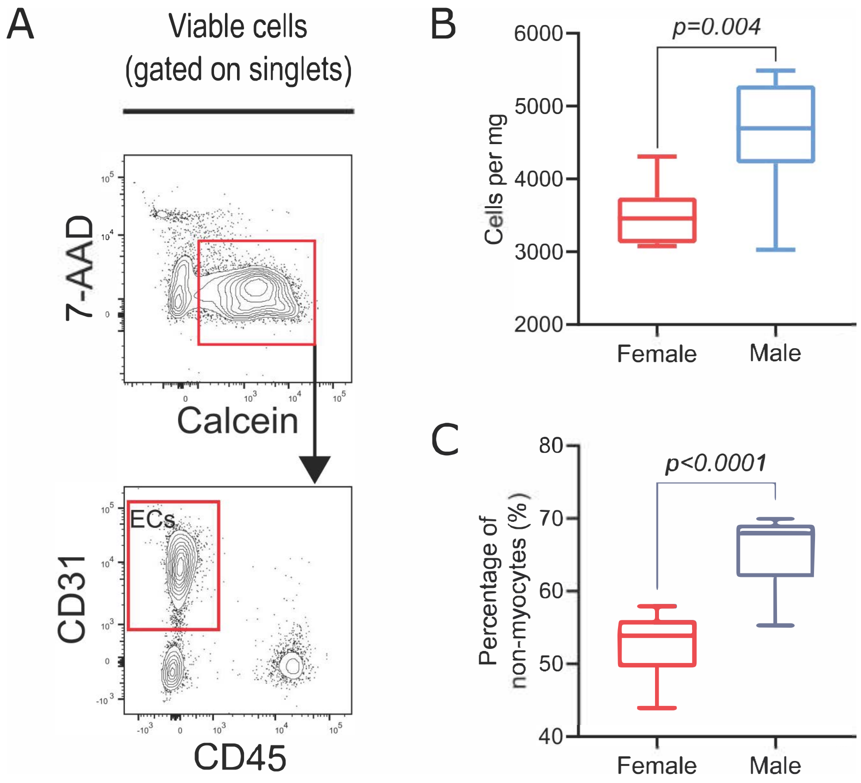


**Online Figure 22. Abundances of endothelial cells in female and male hearts determined by flow cytometry**

A) Flow cytometry contour plots outlining the gating strategy used to determine frequency of endothelial cells. B) Absolute number of endothelial cells normalized to heart tissue mass. C) Proportion of endothelial cells relative to total non-myocytes.

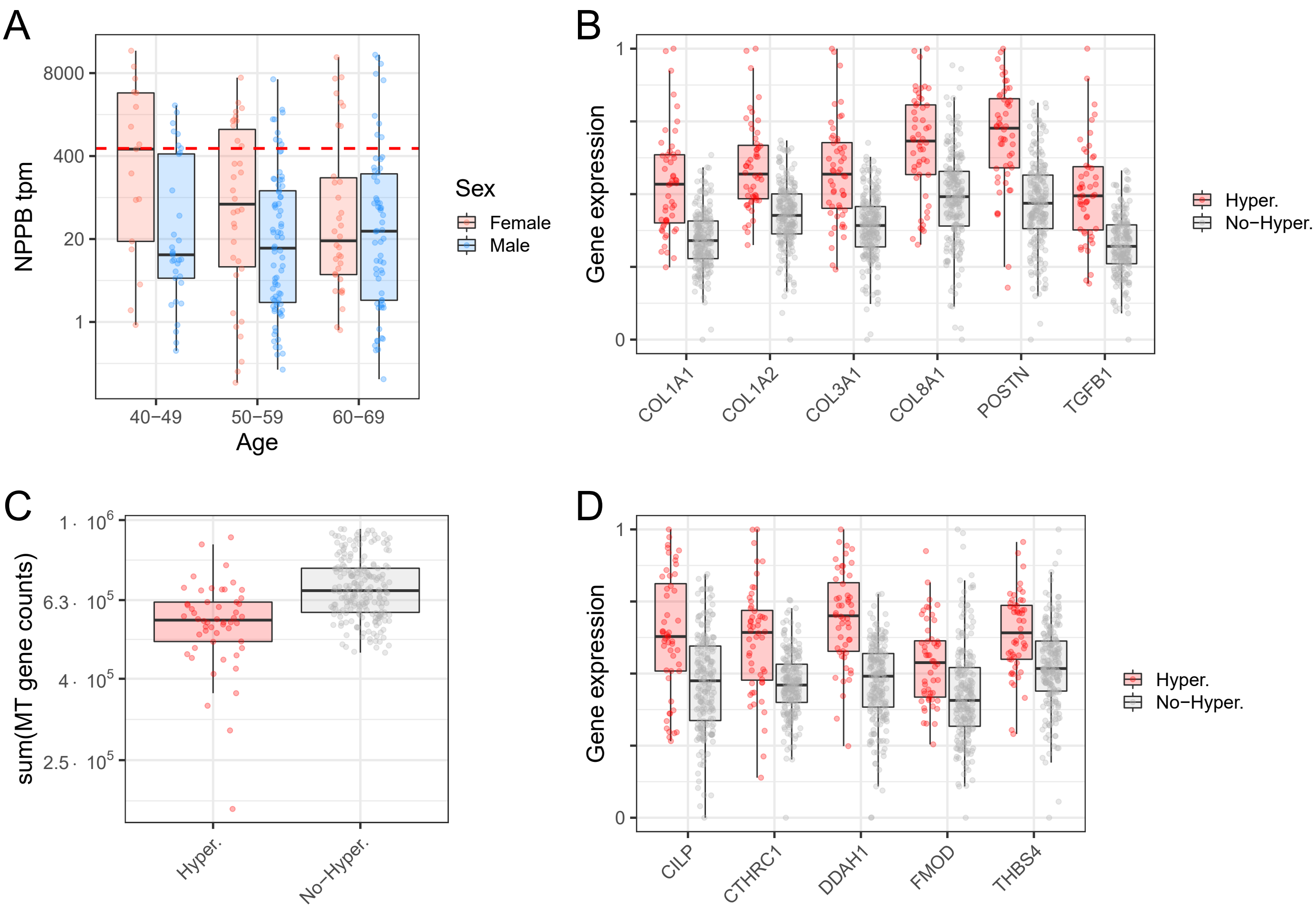


**Online Figure 23. GTEx bulk RNA-seq analysis from human hearts.**

A) Expression of NPPB (in units of transcripts per million; tpm) in 257 female and male human hearts as a function of age and sex. Dotted red horizontal line indicates threshold used for classifying samples as putatively hypertrophic (highest 20% NPPB expression without regard to age or sex). B) Expression of markers of pathological cardiac remodeling in human hearts. Boxplots show identical human samples to (A) classified as positive or negative for putative hypertrophy. To facilitate comparison of patterns for genes that have different absolute expression levels, gene expression levels (in units of tpm) were rescaled to set a maximum of one and minimum of zero. P-values provided above each gene are derived from linear models for the effect of hypertrophy classification while controlling for age and sex. C) Summed expression of transcripts of genes encoded by mitochondrial DNA for identical human samples to (A) classified as positive or negative for putative hypertrophy. P-values provide below the boxplots is derived from a linear model for the effect of hypertrophy classification while controlling for age and sex. D) Expression of human orthologs to markers of Fibroblast-Thbs4 in human hearts. Boxplots show identical human samples to (A) classified as positive or negative for putative hypertrophy. To facilitate comparison of patterns for genes that have different absolute expression levels, gene expression levels (in units of tpm) were rescaled to set a maximum of one and minimum of zero. P-values provided above each gene are derived from linear models for the effect of hypertrophy classification while controlling for age and sex.

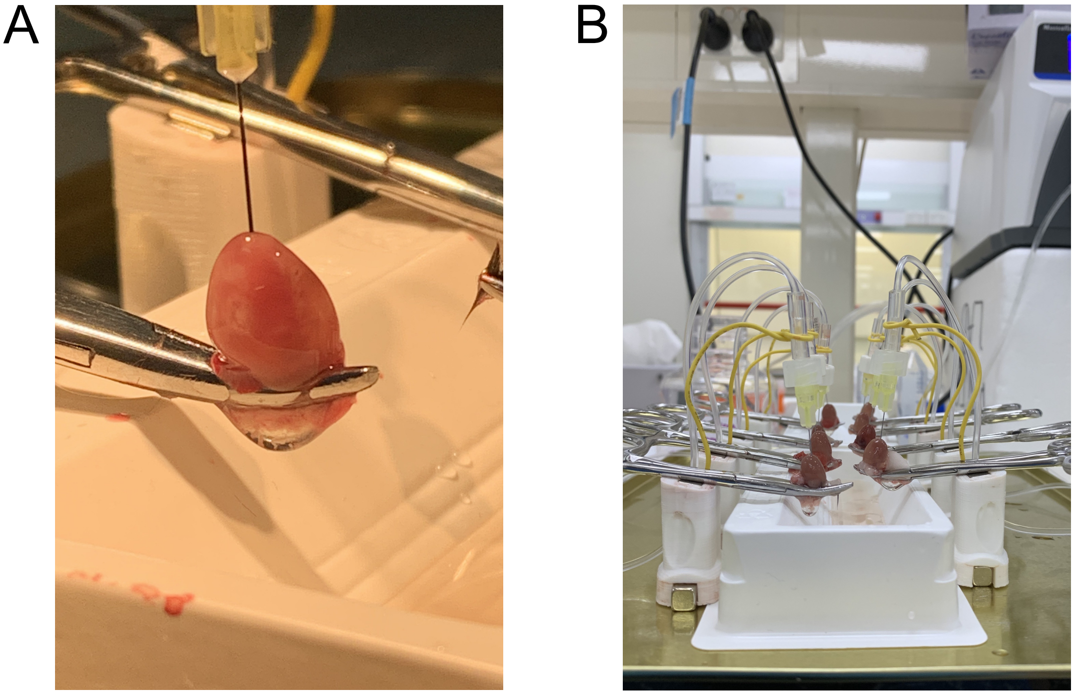


**Online Figure 24. Cardiac tissue perfusion assembly for parallel perfusion-based dissociation of cardiac cells from multiple hearts.**
A) A close-up view of a heart being perfused through the apex by digestion enzymes. B) Equipment layout for simultaneous perfusion of multiple hearts.
